# Therapeutic Blockade of Type 2 Cytokines and PD1 Unleashes Anti-Tumor Immunity Through Coordinated Reprogramming of Innate and Adaptive Immune Surveillance

**DOI:** 10.1101/2025.09.26.678820

**Authors:** Thomas Fabre, Eleanore Hendrickson, Katherine McGourty, Paola M. Marcovecchio, James McMahon, Sara Patti, KaiLi Cage, Stephen Searles, Christella Widjaja, Christopher Stairiker, Kathryn Bound, Roxanne Martino, Edward Pyszczynski, Ying Zhang, Graham Thomas, Samuel A. Stoner, Keith A. Ching, Fang Jin, Christopher P. Dillon, Thomas A. Wynn, Richard L. Gieseck, Alexander M. S. Barron

## Abstract

**Background:** Checkpoint inhibitors improve survival in patients with several types of tumors. However, resistance to checkpoint inhibitors creates an opportunity for patients to benefit from novel immunotherapies. The type 2 cytokines IL-4, IL-13 and TSLP have been implicated in suppressing anti-tumor immune responses through T and myeloid cells. Our current study tested whether combined therapeutic blockade of IL-4, IL-13 and TSLP improved anti-tumor immunity alone and in combination with PD1 antagonism.

**Methods:** We used *in vitro* experiments with primary cells to identify cell types likely to participate in controlling tumors upon IL-4, IL-13, TSLP and PD1 blockade. Therapeutic blockade in the subcutaneous CT26 model tested *in vivo* tumor growth inhibition and associated immunological changes. Bioinformatic analysis of human tumor bulk RNA sequencing data probed for survival associations with IL-4/IL-13 and TSLP transcriptional responses.

**Results:** *In vitro*, IL-4 suppressed T cell-mediated tumor growth inhibition and reduced monocyte-derived dendritic cell expression of proteins associated with anti-tumor immunity. *In vivo*, blocking IL-4, IL-13, TSLP and PD1 improved tumor growth inhibition by creating “hotter” tumors. This was associated with repolarization of CD4 and CD8 T cells and shifts in monocyte, conventional type 1 and type 2 (or monocyte-derived) dendritic cell programs. Transcriptional responses to IL-4/IL-13 and TSLP were associated with poor survival outcomes across patients with several types of cancers.

**Conclusion:** Therapeutic blockade of IL-4, IL-13 and TSLP may drive immunological tumor growth inhibition in subsets of cancer patients alone and in combination with checkpoint inhibitors. Improved tumor growth inhibition was likely driven through augmented cytotoxic T cell priming in secondary lymphoid organs and improved reactivation by repolarized monocytes and dendritic cells in tumors.

## Background

Checkpoint inhibitor therapies benefit subsets of patients with several types of solid tumors (1–3). Up to 61% of melanoma patients treated with nivolumab, an anti-Programmed death 1 (PD1) blocking antibody, have responses lasting at least five years(4). Clinical responses to checkpoint inhibitors are frequently associated with type 1 immunity including Interferon gamma (IFNγ) signaling and CD8 T cell infiltration(5–8). Despite these clinical successes, some solid tumors are intrinsically resistant to checkpoint inhibitors and others acquire resistance over time(4,9–11).

Resistance can occur through multiple mechanisms, and one may be the inhibition of type 1 immunity by type 2 cytokines(12–17). Interleukin-4 (IL-4) and IL-13 are canonical type 2 cytokines that signal through two heterodimeric receptors(18,19). IL-4 can suppress the production of IFNγ by CD4 and CD8 T cells and can impair the cytotoxic activity of effector CD8 T cells(20–24). This is partially the result of transcriptional repression of a type 1 program that includes *Ifng, Il12rb2, Stat4* and *Tbx21* by STAT6 and GATA-3(25–35). IL-4 and IL-13 signaling in myeloid cells can protect tumors through a variety of mechanisms including reducing IL-12 production(12,14,36–42). Consistent with deleterious roles for IL-4 and IL-13 in anti-tumor immunity, inhibiting this cascade frequently improves immunological control of tumor growth(12,14,36,43–50).

Thymic stromal lymphopoietin (TSLP) is an alarmin associated with type 2 immune responses and atopic diseases(18,51). Unlike IL-4 and IL-13, the long isoform TSLP transcriptional program is largely executed by STAT5(51–54). As an alarmin, TSLP matures and polarizes type-2 conventional dendritic cells (cDC2) from humans and mice(55,56). By inducing a program that lacks IL-12 and includes OX40 ligand (OX40L), TSLP creates cDC2 that polarize naïve CD4 T cells into T helper type 2 (T_H_2) and T follicular helper type 2 cells(55,57–61). cDC2 matured by TSLP also potently reactivate memory T_H_2 cells thus triggering secretion of IL-13(56). In addition to polarizing and restimulating T_H_2 responses, naïve CD8 T cells primed by TSLP-matured DC may produce IL-13 and exhibit weak cytotoxicity(62). Although the roles of TSLP are context-specific, data from breast and pancreatic cancers indicate that TSLP-matured DC drive harmful type 2-skewed responses in some patient populations(63–68).

This evidence suggests that IL-4, IL-13 and TSLP can suppress anti-tumor immunity independent of conventional checkpoints like PD-1. We hypothesized that combined blockade of these three cytokines would inhibit tumor growth alone and in combination with PD1 antagonism. Using *in vitro*, *in vivo* and *in silico* experiments, we demonstrated that IL-4, IL-13 and TSLP suppressed anti-tumor immunity. Blocking IL-4, IL-13, TSLP and PD1 improved tumor growth inhibition (TGI) which was associated with repolarization of DC and T cells in the tumor microenvironment (TME) and secondary lymphoid organs (SLO). Together these data support combined blockade of IL-4, IL-13, TSLP and PD1 as a novel cancer immunotherapy.

## Materials and Methods

### In vitro *T cell priming and tumor growth inhibition assay and cytokine secretion measurement*

Total CD4 and CD8 T cells were isolated by negative selection from three healthy donors as described in the online supplemental methods. Apoptosis was induced in A375 human tumor cells by treatment with mitomycin C (Stemcell Technologies). Apoptotic A375 cells were mixed with equal numbers of CD4 or CD8 T cells and cultured with cytokines and antibodies for 11-12 days as described in the online supplemental methods. At the end of this priming phase cytokines were quantified in the conditioned medium using V-plex kits (Meso Scale Diagnostics) per the manufacturer’s instructions.

Expanded CD4 and CD8 T cells from the priming cultures were mixed at a 1:1 ratio and co-cultured with A375 tumor cells as described in the online supplemental methods. TGI was measured by longitudinal GFP imaging and counting on an Incucyte S3 microscope (Sartorius). Growth curves were normalized to the initial number of cells and area under the curves (AUC) calculated using Prism (GraphPad). Lower AUC measurements indicate fewer tumor cells and better TGI.

### In vitro *murine bone marrow-derived DC2-like differentiation and stimulation assay*

Bone marrow was isolated from C57BL/6 femurs by centrifugation as previously described (69). Monocytes or hematopoietic stem cells (HSC) were isolated by negative selection using EasySep kits (Stemcell Technologies) per the manufacturer’s instructions. 7.5×10^6^ HSC/225cm^2^ flask or monocytes at 4×10^6^/25cm^2^ flask were differentiated in RPMI1640 supplemented with 2-mercaptoethanol (55μM, ThermoFisher Scientific), 1× nonessential amino acids (ThermoFisher Scientific), 1mM sodium pyruvate (ThermoFisher Scientific), 10mM HEPES (ThermoFisher Scientific), recombinant murine 20ng/mL GM-CSF (Biolegend) and 100ng/mL recombinant murine Flt3L (Peprotech) for eight days. Media and cytokines were refreshed every 2-3 days. Differentiated bone marrow dendritic cells were isolated by CD11c positive selection kit (Stemcell Technologies) per the manufacturer’s instructions, and the purity of bone marrow derived conventional type 2 dendritic-like cells (BMDC2) determined by flow cytometry.

BMDC2 were plated at 1×10^5^ in flat-bottomed 96-well tissue culture treated plates (BD Falcon). Cells were stimulated with lipopolysaccharide (LPS-EB, Invivogen) at 5 μg/mL to mature BMDC2. Recombinant murine IL-4 (50ng/mL, Biolegend), IL-13 (50ng/mL, Biolegend) and TSLP (50ng/mL, R&D Systems) or a combination of the three cytokines were added concurrently with LPS to the appropriate wells. After 24 hours, conditioned media was collected for secreted protein analysis and cells were collected for flow cytometry by detaching with Accutase (ThermoFisher Scientific). Secreted proteins were quantified using Legendplex kits (Biolegend) per the manufacturer’s instructions.

### Derivation of IL-4/IL-13 and TSLP transcriptional response signatures

Transcriptional response signatures were generated in collaboration with CytoReason (Tel Aviv, Israel) by combining published data detailing genes transactivated in response to IL-4/IL-13 or TSLP signaling with pathway data from public databases. Genes encoding the proteins that combine to form the type I and type II IL-4/IL-13 receptors or the TSLP receptors were added to the respective signature lists. These lists were filtered by calculating Spearman correlation coefficients against Cytoreason’s database of bulk RNAseq from purified cell types and whole tissues from patients with inflammatory diseases. Only genes with ρ>0.70 were included in the final lists. These resulting transcriptional response signature gene lists are shown in Supplemental Table 3.

### In vitro *murine tumor cell stimulations, flow cytometry and secreted protein quantification*

Murine CT26 colon adenocarcinoma cells were cultured in DMEM with 4.5g/L glucose and GlutaMAX-I (ThermoFisher) supplemented with 10% fetal bovine serum (ThermoFisher), penicillin and streptomycin (ThermoFisher) and 10mM HEPES (ThermoFisher). Cells were stimulated with recombinant murine IL-4, IL-13, TSLP, IL-1β or TNFα (all from Biolegend) at 10ng/mL for 18 hours. Media was collected and frozen at -80°C before secreted cytokines were quantified using Legendplex fluid-phase immunoassays (Biolegend). Stimulated cells were washed with PBS and detached with TrypLE Express (ThermoFisher). These were stained with Zombie NIR viability dye and the antibodies listed in Supplemental Table 2.

### *Mice, tumor transplantation,* in vivo *treatments and tissue collection*

Female Balb/C J mice were housed under specific pathogen-free conditions and subcutaneously implanted in the right hind flank with approximately 500,000 CT26 cells. On day 9 when visible tumors averaged 75mm^3^ (as defined by (length*width^2^)/2), mice were randomized into groups (n=10/group) immediately prior to dosing. The treatment groups were: (1) isotype control, (2) anti-IL-4 (clone 11B11, Pfizer, Inc.) plus mIL13Ra2-mFc (Pfizer, Inc.), (3) anti-PD1 (clone F2, Pfizer, Inc.), (4) anti-IL-4 plus mIL13Ra2-mFc plus anti-PD1, (5) anti-IL-4 plus mIL13Ra2-mFc plus anti-TSLP (clone 28F12, ThermoFisher) or (6) anti-IL-4 plus mIL13Ra2-mFc plus anti-TSLP plus anti-PD1. Anti-IL-4 (10mg/kg) and mIL13Ra2-mFc (10mg/kg) were injected subcutaneously while anti-PD1 (10mg/kg) and anti-TSLP (10mg/kg) were injected intraperitoneally, every 3-4 days, for a total of 5 doses. Isotype-matched control antibodies were administered such that the total mass of protein per dose was equivalent across treatment groups. Tumor volumes were longitudinally tracked and animals were sacrificed on day 22 post-tumor implantation. Tumors, tumor-draining axillary lymph nodes and spleens were collected and processed as described in online supplemental methods. All procedures performed on animals were in accordance with regulations and established guidelines and were reviewed and approved by an Institutional Animal Care and Use Committee or through an ethical review process.

#### Flow cytometry

Viability was determined by staining with LIVE/Dead Blue (ThermoFisher Scientific) or Zombie NIR (Biolegend). Fc receptors were blocked with mouse or human TruStain FcX (Biolegend) as appropriate. Surface and intracellular staining was performed using the fluorochrome-labeled antibodies for human cells listed in Supplemental Table 1 or murine cells listed in Supplemental Table 2. Data was acquired and unmixed on a Cytek Aurora flow cytometer and analyzed using OMIQ software by a combination of traditional two-dimensional hierarchical gating and dimensionality reduction (Dotmatics). The two-dimensional gating schemes for murine tumor samples (Supplemental Figure 1), human monocyte-derived macrophages (Supplemental Figure 2) and human T cells (Supplemental Figure 3) can be found in the online supplemental data. All PaCMAP dimensionality reduction and PARC clustering was performed using the settings listed in online supplemental methods. The features used for PaCMAP and PARC are listed in the Results section. Mass-normalized cell counts were generated by dividing the number of cells by the mass of the piece of tumor digested for analysis by flow cytometry.

### Ex vivo *restimulation of dissociated murine tissues and secreted protein quantification*

Single cells from the TME or spleen were restimulated with phorbol 12-myristate 13-acetate (PMA) and ionomycin (ThermoFisher) in complete RPMI (ThermoFisher) for 20 hours as described in the online supplemental methods. Secreted proteins were quantified using fluid-phase Legendplex immunoassays (all from Biolegend) per the manufacturer’s instructions.

### Quantification of proteins in murine serum samples

Frozen murine serum samples were thawed, diluted 1:100 (to quantify IgE) or 1:2 (all other analytes) in Assay Buffer and serum proteins quantified with the mouse IgE or proinflammatory chemokine (13-plex) Legendplex fluid-phase immunoassays (both from Biolegend) per the manufacturer’s instructions.

### RNA isolation, bulk RNA sequencing of murine tissues and data processing

Tumor pieces and TDLN from three mice per treatment group in RNAlater (ThermoFisher Scientific) were transferred into 2mL CKMix tubes (Precellys) and homogenized using a Precellys Evolution (Precellys) on dry ice. Tissues were then resuspended in RLT buffer (Qiagen) supplemented with 2-mercaptoethanol (ThermoFisher Scientific) and RNA isolated using the RNeasy 96 kit (Qiagen) per the manufacturer’s instructions. The RNA concentration was determined using the Qubit RNA High Sensitivity Kit (Thermo Fisher Scientific). RIN measurements were determined by using the 4200 Tapestation System (Agilent). The TruSeq Stranded mRNA library preparation kit (Illumina) was utilized for library preparation. Library QC was determined using the Qubit dsDNA HS assay kits (Thermo Fisher Scientific), and the library size measurement was carried out using the 4200 TapeStation system (Agilent). Libraries were normalized, pooled then sequenced on the NovaSeq 6000 Platform on the NovaSeq S4 300 cycle flowcell (Illumina). 151-8-8-151 sequencing parameters were utilized. Demultiplexing was performed using Illumina’s bcl2fastq2-2.20 tool to generate fastq files. Reference mapping and conversion to log2-transformed transcripts per million (TPM) were performed as described in(70). Briefly, raw reads were mapped to the vM17 reference genome (Gencode) and gene expression quantified using RSEM v1.3.0 and STAR v020201. These tables were uploaded to the Data4Cure, Inc. Biomedical Intelligence Cloud (https://www.data4cure.com) for analysis.

### Analysis of bulk RNA sequencing data

RNA and transcriptional response signature expression analyses on murine cancer cell line and tumor model data, and on The Cancer Genome Atlas (TCGA) Pan-Cancer Atlas data were conducted with the Data4Cure, Inc. Biomedical Intelligence Cloud (https://www.data4cure.com). Comparisons between mouse tumor models used previously published data available from the Sequence Read Archive under accession number <u>PRJNA505989</u>(70). Analyses of patient data used previously published TCGA data available from the <u>National Cancer Institute’s GenomicData Commons</u>(71). Gene expression analysis used either log_2_ TPM (murine data) or upper quantile-normalized fragments per kilobase of transcript per million mapped reads (UQ-FPKM, human data). Transcriptional response signature analysis used either the median (IL-4/IL-13 and TSLP transcriptional response signatures) or sum (T_H_2 signature) of expression of the genes in the signature and were Z-scored. RNA and transcriptional expression data were exported and analyzed in PRISM (GraphPad). Cox proportional hazard models were run on the Data4Cure, Inc. Biomedical Intelligence Cloud using default settings for median split analysis.

### Statistical analyses

All the statistical testing except for the analysis of covariance (ANCOVA) for CT26 tumor volumes and Cox proportional hazard models for TCGA data was conducted using PRISM (GraphPad). Differences between groups were identified with one-or two-way analysis of variance (ANOVA) or Student’s t-test as appropriate (noted in the figure legends). Post-hoc tests for the differences between the means of selected groups were conducted with the tests noted in the figure legends. P-values for post-hoc pairwise comparisons are displayed on each graph.ANCOVA analysis for CT26 tumor volumes was performed with an internal Pfizer statistical analysis platform. Briefly, tumor volumes were log_10_ transformed, with ANCOVA applied to the log transformed data at each time point. Comparison of each treatment group to the isotype control group was performed using a 1-sided test. Contrast comparisons between other treatment groups were performed using a 2-sided test. To determine whether drug combinations resulted in additive, or independent, responses, the Bliss independence model was used. Additive efficacy is confirmed if the Bliss predicted value of combinatorial efficacy is statistically insignificant compared to experimental efficacy using the formula Y_a+b_=Y_a_+ Y_b_− Y_a_Y_b_, where Y_x_ represents the ratio of treatment tumor volume to control. Contrast comparisons between combination treatment groups and predicted Bliss response at each time point were performed using a 2-sided ANCOVA test.

## Results

### Type 2 cytokines suppress T cell-mediated tumor growth inhibition and type 1 polarizing cDC2 maturation in vitro

To dissect suppression of anti-tumor immunity by IL-4, IL-13 and TSLP, we tested effects of these cytokines on primary human T cells, DCs and macrophages *in vitro*. First, we used a T cell tumor growth inhibition (TGI) assay where cytokines and blockades were applied during “priming” and TGI (Figure 1A). A375 growth was unaffected by any of the treatments in the absence of T cells (Supplemental Figure 4A). Recombinant human IL-12 was used as a positive control which enhanced TGI for two donors (Fig. 1B). IL-4 impaired TGI and increased the IL-13/IFNγ secretion ratio (Fig. 1B-D, Sup. Fig. 4B-E). Antibody blockade of IL-4 both improved TGI and decreased the IL-13/IFNγ secretion ratio (Fig. 1B-D, Sup. Fig. 4B-E). Thus, IL-4 suppressed TGI by acting directly on T cells and blocking IL-4 improved TGI and decreased type 2 T cell polarization.

**Figure 1.**
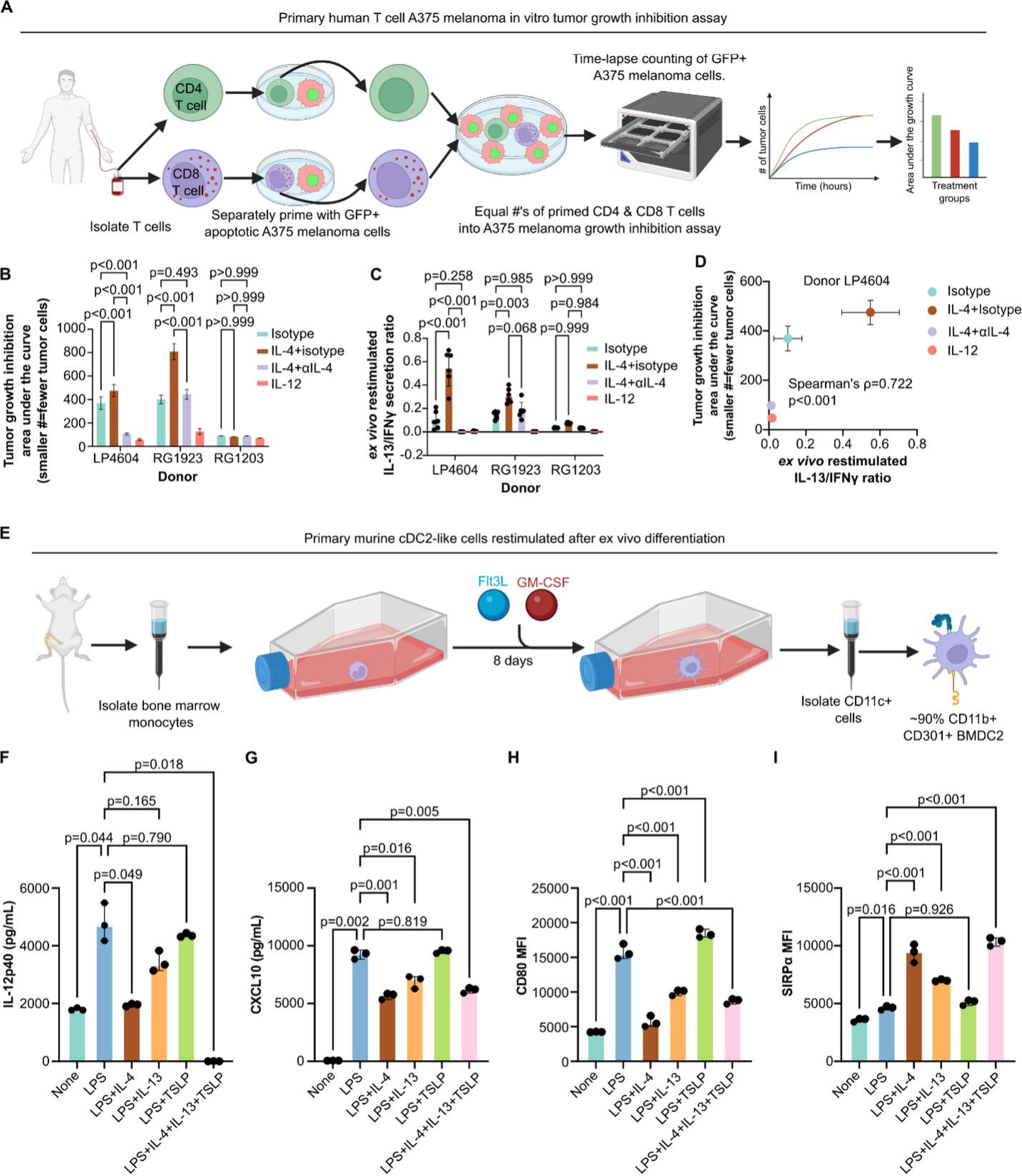
IL-4, IL-13 and TSLP suppress anti-tumor T cell responses directly and indirectly through cDC2 polarization *in vitro*. **(A)** *In vitro* primary human T cell tumor growth inhibition (TGI) experimental workflow cartoon. (**B**) Area under the TGI curves (AUC) for primary human T cells from three donors exposed to recombinant human IL-4 or IL-12 and isotype or IL-4 blocking antibody (αIL-4) during the priming and growth inhibition phases from (A). **(C)** Ratio of IL-13/IFNγ proteins secreted by the same primary T cells from (B) during the priming phase as measured by fluid phase immunoassay. **(D)** Correlation of AUC from (B) with IL-13/IFNγ ratio from (C) for donor LP4604. **(E)** Cartoon depicting generation of murine bone marrow derived cDC2-like cells. **(F-G)** Secretion of IL-12p40 (F) or CXCL10 (G) proteins by murine cDC2-like cells stimulated with lipopolysaccharide (LPS) alone or in combination with recombinant murine IL-4, IL-13, TSLP or the combination of IL-4, IL-13 and TSLP as measured by fluid phase immunoassay. **(H-I)** Surface expression of CD80 (H) or SIRPα (I) by murine cDC2-like cells stimulated as in (F-G). For (B-C) bars and whiskers represent the mean and standard deviation of six biological replicates per treatment per patient. Statistical tests are two-way ANOVAs (treatment p<0.001, donor p<0.001 and interaction p<0.001 for both TGI AUC and IL-13/IFNγ ratio) with post-hoc Tukey’s multiple comparison tests (p-values shown in figure). For (D) points and whiskers represent the mean and standard deviation of six biological replicates for donor LP4604. Spearman’s correlation ρ and p-values are displayed in the figure. For (F-I) bars and whiskers represent the median and 95% confidence interval for three biological replicates. Statistical tests are one-way ANOVAs (IL-12p40 p<0.001, CXCL10 p<0.001, CD80 p<0.001, SIRPα p<0.001) with post-hoc Sidak’s multiple comparison tests (p-values shown in figure). Cartoons made with BioRender.

Next, we tested the hypothesis that IL-4, IL-13 and TSLP suppress anti-tumor immunity through DCs. We generated murine bone marrow-derived type-2 conventional dendritic-like cells (BMDC2) as shown in Fig. 1E, which resulted in approximately 90% immature BMDC2s (Sup. Fig. 5). Maturation with lipopolysaccharide (LPS) induced secretion of IL-12p40 and CXCL10, two cytokines associated with anti-tumor immunity (Fig. 1F-G). IL-4 suppressed secretion of both IL-12p40 and CXCL10 while IL-13 suppressed CXCL10 secretion (Fig. 1F-G). Although TSLP alone did not alter the secretion of either cytokine, combined IL-4, IL-13 and TSLP completely abrogated secretion of IL-12p40 (Fig. 1F). Similarly, IL-4 inhibited secretion of IL-12p40 and IL-12p70 by primary human monocyte-derived DCs (Sup. Fig. 6). IL-4 and IL-13 also reduced surface expression of the co-stimulatory molecule CD80 and increased SIRPα expression (Fig. 1H-I). These data are consistent with IL-4, IL-13 and TSLP suppressing the production of factors necessary for type 1 T cell polarization and TGI by DCs.

Prior work suggested that IL-4 and IL-13 signaling in macrophages (i.e. M2 or M_IL-4_ or M_IL-13_) suppressed anti-tumor T cell responses(36,48). We tested this *in vitro* using primary human monocyte-derived macrophage (MDM)-T cell co-cultures followed by TGI (Sup. Fig. 7A). Surprisingly, MDM polarized with IL-4 enhanced TGI (Sup. Fig. 7B-C). Consistent with the enhanced TGI, MDM polarized with IL-4 had increased expression of MHC-II, CD86, and CD40 (Sup. Fig. 7D-F). Collectively, these data suggest that IL-4, IL-13 and TSLP suppress anti-tumor immunity by acting on T cells and DCs and not through macrophages.

### TGI from blocking IL-4, IL-13 and TSLP is comparable to PD1 antagonism, and quadruple inhibition yields additive efficacy in CT26 tumors

Next, we sought a murine tumor model to test our hypothesis that blocking IL-4, IL-13 and TSLP would enhance anti-tumor immunity. Bulk RNA sequencing (RNAseq) from several subcutaneous tumor models identified 4T1 and CT26 as the only two with detectable transcripts for all three cytokines (Sup. Fig. 8A)(70). To address signaling downstream of these cytokines, we quantified levels of IL-4/IL-13 and TSLP transcriptional response signatures and a T_H_2 deconvolution signature. When we combined the responses to IL-4/IL-13 and TSLP with the T_H_2 signature, CT26 tumors had the highest score despite a modest TSLP response (Sup. Fig. 8B). It was unlikely that CT26 cells were directly responsible for these responses as pure cultures were unresponsive *in vitro* (Sup. Figs. 9-12). This, combined with a prior publication demonstrating efficacy for IL-4 blockade in CT26 tumors, led us to select this model for our study(36).

Our subcutaneous CT26 tumor study was designed to test several therapeutic hypotheses: first, that blocking IL-4, IL-13 and TSLP (αIL-4/IL-13/TSLP) would demonstrate superior TGI to blocking IL-4 and IL-13 (αIL-4/IL-13) alone or an isotype control; second, that the TGI from quadruple blockade of IL-4, IL-13, TSLP and PD1 (αIL-4/IL-13/TSLP/PD1) would be superior to the isotype control, αIL-4/IL-13/TSLP, αPD1, and αIL-4/IL-13/PD1 groups. Individual longitudinal growth curves showed variation with three mice in the αIL-4/IL-13/TSLP/PD1 group having large tumors (Fig. 2A-F, these are marked with open circles in subsequent graphs). These varied responses were likely due to biological heterogeneity rather than a lack of target coverage as serum levels of the antagonists were comparable within and across treatment groups (Sup. Fig. 13). Median growth curves and terminal tumor volumes demonstrated that the αIL-4/IL-13/TSLP/PD1 and αIL-4/IL-13/TSLP treatments improved TGI compared to the isotype (Fig. 2G-H, Sup. Table 4). Using the Bliss independence model, we demonstrated that the αIL-4/IL-13/TSLP/PD1 combination had additive efficacy versus αIL-4/IL-13/TSLP or αPD1 alone (Fig. 2I). Together, these data indicate that αIL-4/IL-13/TSLP and αIL-4/IL-13/TSLP/PD1 induced TGI with the quadruple blockade yielding additive TGI in the CT26 model.

**Figure 2.**
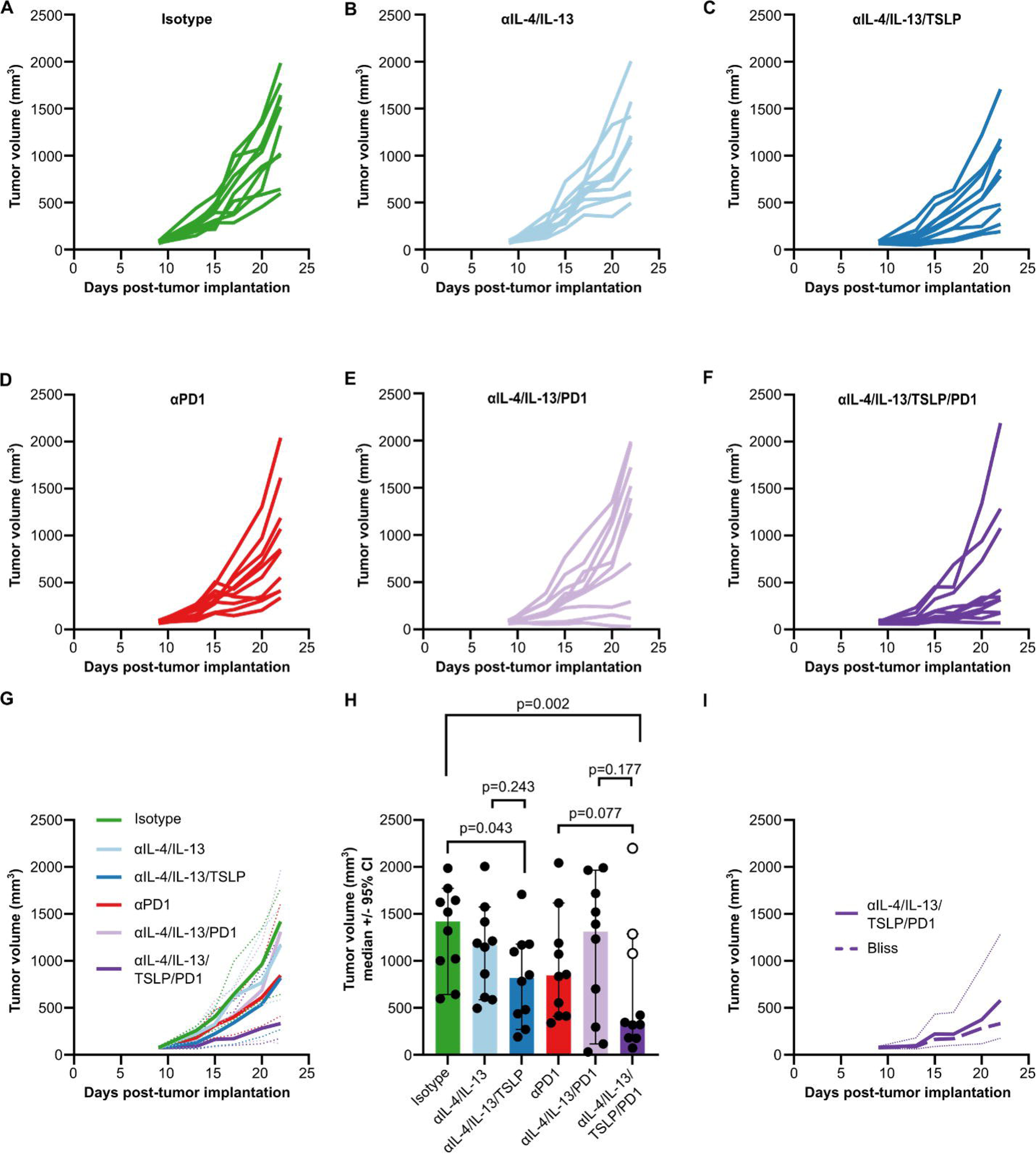
Combined therapeutic blockade of IL-4, IL-13 and TSLP demonstrated additive tumor growth inhibition in combination with PD1 antagonism. (A-F) Longitudinal subcutaneous CT26 tumor volume curves for individual mice per treatment group: (A) isotype; (B) combined blockade of IL-4 and IL-13 (αIL-4/IL-13); (C) combined blockade of IL-4, IL-13 and TSLP (αIL-4/IL-13/TSLP); (D) PD1 blockade (αPD1); (E) combined blockade of IL-4, IL-13 and PD1 (αIL-4/IL-13/PD1); (F) quadruple blockade of IL-4, IL-13, TSLP and PD1 (αIL-4/IL-13/TSLP/PD1). (G) Median (solid lines) longitudinal subcutaneous CT26 tumor volume curves for the isotype (green), αIL-4/IL-13 (light blue), αIL-4/IL-13/TSLP (dark blue), αPD1 (red), αIL-4/IL-13/PD1 (lavender) and αIL-4/IL-13/TSLP/PD1 (purple) groups with 95% confidence intervals (dotted lines). **(H)** Terminal subcutaneous CT26 tumor volumes on day 22 of the study. Each dot represents an individual mouse, the bar represents the group median and the whiskers show the 95% confidence intervals. ANCOVA was used to test for differences between treatment groups. (I) Comparison of the median (thick solid purple line) and 95% confidence intervals (thin dotted purple lines) of the experimental tumor volume data for the αIL-4/IL-13/TSLP/PD1 group against the Bliss independence model (thick dashed purple line).

### Therapeutic CT26 tumor growth inhibition was associated with elevated tumor infiltrating leukocytes and systemic changes in T and myeloid cells

To identify changes associated with the TGI of the αIL-4/IL-13/TSLP and αIL-4/IL-13/TSLP/PD1 treatments, we investigated tumor infiltration by leukocyte lineages using flow cytometry. αIL-4/IL-13/TSLP/PD1 enhanced T and myeloid cell densities in the TME with αIL-4/IL-13/TSLP driving a smaller increase (Fig. 3A-G). However, only the numbers of CD8 T cells, classical monocytes, pDC and DC2 were negatively correlated with terminal tumor volumes (Fig. 3H-K). Neither αIL-4/IL-13/TSLP nor αIL-4/IL-13/TSLP/PD1 altered the frequency of CD8 T cells in the tumor draining lymph node (TDLN) nor was this correlated with tumor volumes (Sup. Fig. 14A-B). Similarly, the frequencies of migratory cDC2 and cDC1 in the TDLN were unchanged in both groups, and neither was correlated with tumor volume (Sup. Fig. 14C-F). Splenic CD8 T cell frequencies increased in both the αIL-4/IL-13/TSLP and αIL-4/IL-13/TSLP/PD1 groups, and these were inversely correlated with tumor volumes (Sup. Fig. 14G-H). Together, these data suggest that enhanced CD8 T cell emigration from TDLN or increased CD8 T cell proliferation outside of the TDLN is responsible for the increase in tumor-infiltrating CD8 T cells in the αIL-4/IL-13/TSLP/PD1 group.

**Figure 3.**
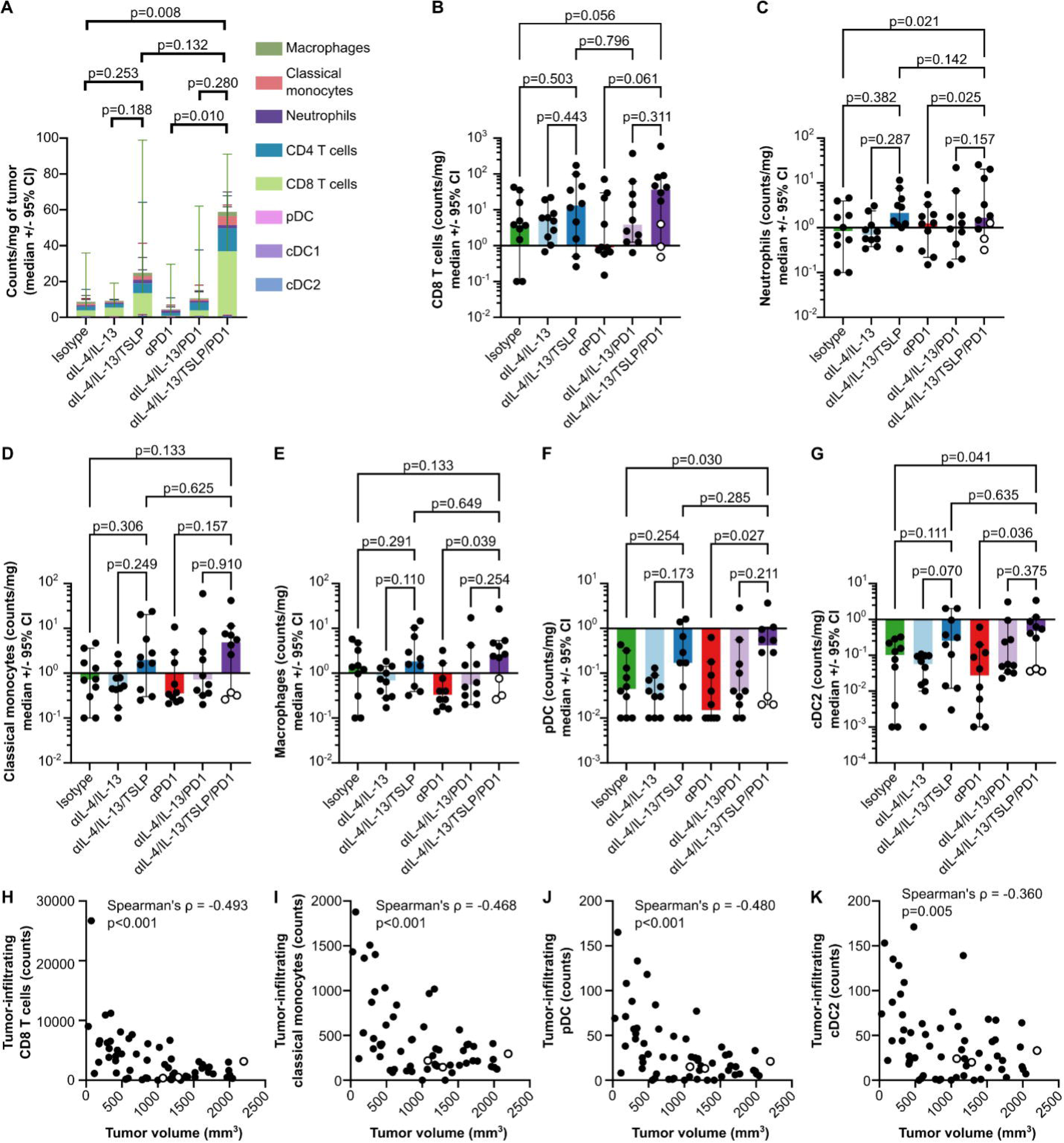
Therapeutic subcutaneous CT26 tumor growth control was associated with an increased density of tumor infiltrating leukocytes. **(A)** Mass-normalized counts of selected tumor-infiltrating leukocyte lineages as measured by flow cytometry. Bars represent the median value of each group (n=10 mice/group), and whiskers represent the 95% confidence intervals. Two-way ANOVA testing for differences between cell types within treatments (p<0.001) and total cell counts/mg of tumor between treatments (p=0.030) followed by uncorrected Fisher’s LSD (p-values for select groups shown) were used. **(B-G)** Individual plots of mass-normalized tumor infiltrating counts from (A). Each dot represents an individual mouse, bars represent the median and whiskers the 95% confidence intervals. Ordinary one-way ANOVA (CD8 T cells p=0.302, neutrophils p=0.122, classical monocytes p=0.353, macrophages p=0.211, pDC p=0.127 and cDC2 p=0.097) were followed by uncorrected Fisher’s LSD tests (p-values shown in figure). **(H-K)** Correlations of tumor infiltrating counts of CD8 T cells, classical monocytes, pDC or cDC2 with terminal tumor volumes. Spearman’s ρ and p-values are shown in the figure. Each dot represents an individual mouse.

Extramedullary hematopoiesis occurs in some cancer patients, and immunosuppressive splenic myelo- and granulopoiesis occurs in the subcutaneous CT26 model(72,73). Both αIL-4/IL-13/TSLP and αIL-4/IL-13/TSLP/PD1 treatments decreased the frequency of splenic neutrophils which was positively correlated with tumor volume (Sup. Fig. 15A-B). Splenic macrophages were also positively correlated with tumor volume and decreased in the αIL-4/IL-13/TSLP/PD1 group (Sup. Fig. 15C-D). αIL-4/IL-13/TSLP/PD1 treatment increased the frequencies of splenic classical monocytes which was negatively correlated with tumor volumes (Sup. Fig. 15E-F). These data suggest that the therapeutic TGI by αIL-4/IL-13/TSLP/PD1 treatment may be partially due to normalization of splenic extramedullary hematopoiesis.

### Increased type 1 immune polarization in the TME and decreased type 2 polarization in TDLN are associated with therapeutic TGI

To test our hypothesis that TGI in the αIL-4/IL-13/TSLP and αIL-4/IL-13/TSLP/PD1 groups was associated with type 2 to type 1 repolarization, we used flow cytometry, cytokine secretion and bulk RNAseq. Although the αIL-4/IL-13/TSLP and αIL-4/IL-13/TSLP/PD1 treatments increased combined tumor infiltration by T_H_1 (CD44+Tbet+) and T_H_2 (CD44+GATA3+) cells, neither increase was significant on its own and, surprisingly, T_H_2 counts were inversely correlated with tumor volume (Fig. 4A-H). Changes in T_H_1 and T_H_2 infiltration do not measure their capacity to produce cytokines which we tested by restimulating TME single cell suspensions with PMA and ionomycin. Because CT26 tumors produce high levels of the decoy IL-13 receptor α2 (IL-13Rα2) we could not accurately quantify IL-13 levels as in Fig. 1 (Sup. Fig. 16). Fewer mice are represented in these graphs due to technical errors in data acquisition. αIL-4/IL-13/TSLP/PD1 increased in the ratio of secreted IFNγ/IL-4, largely due to decreased IL-4 secretion, and IFNγ secretion was inversely correlated while IL-4 secretion was positively correlated with tumor volumes (Fig. 4I-J, Sup. Fig. 17A-D). IFNγ transcriptional responses in a subset of tumors corroborated its inverse correlation with tumor volumes (Fig. 4K-L) (74). These data indicate that TGI induced by the αIL-4/IL-13/TSLP/PD1 treatment was associated with a generally “hotter” TME rather than repolarization.

**Figure 4.**
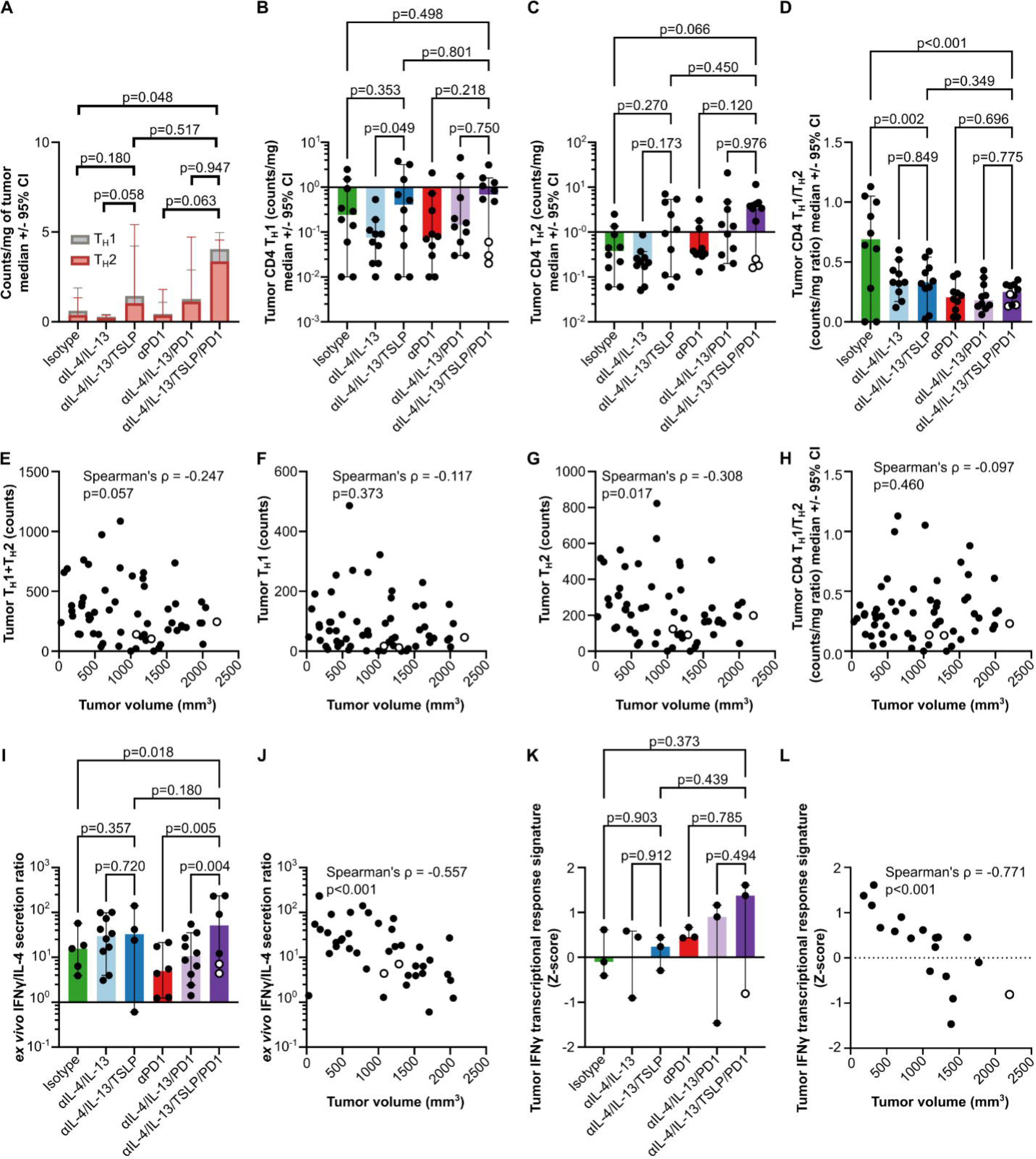
Therapeutically induced tumor growth inhibition was associated with repolarization of the CT26 TME from type 2 towards type 1 immune skewing. **(A)** Mass-normalized counts of tumor infiltrating T_H_1 and T_H_2. Bars represent the median and whiskers the 95% confidence interval of n=10 mice/group. Two-way ANOVA (treatment p=0.042 and cell type p=0.007) followed by uncorrected Fisher’s LSD (p-values shown in figure) were used. **(B-C)** Individual plots of the mass-normalized tumor-infiltrating T_H_1 and T_H_2 counts shown in (A). **(D)** Ratio of mass-normalized tumor infiltrating T_H_1/T_H_2 counts from (A-C). For (B-D) each dot represents an individual mouse. One-way ANOVA (T_H_1 p=0.337, T_H_2 p=0.127, T_H_1/T_H_2 p<0.001) were followed by uncorrected Fisher’s LSD tests (p-values shown in figure). **(E-H)** Correlations between the tumor-infiltrating counts of (E) combined T_H_1+T_H_2, (F) T_H_1, (G) T_H_2 or (H) ratio of T_H_1/T_H_2 with terminal tumor volumes. Spearman’s ρ and p-values are shown in the figure. Each dot represents an individual mouse. (I) The ratio of IFNγ/IL-4 proteins secreted by total single cells isolated from subcutaneous CT26 tumors and restimulated with PMA and ionomycin *ex vivo*. Dots represent data from individual mice, bars show group medians and whiskers display 95% confidence intervals. One-way ANOVA (p=0.047) followed by post-hoc Tukey’s multiple comparisons tests (p-values shown in figure) were used. (J) Spearman’s correlation between terminal tumor volumes and the *ex vivo* IFNγ/IL-4 ratio shown in (A). Each dot represents values from an individual mouse. **(K)** Levels of an IFNγ transcriptional response signature (Z-scored) in a subset of tumors using bulk RNAseq. One-way ANOVA (p=0.911) followed by uncorrected Fisher’s LSD (p-values shown in graph) was used. (L) Correlation of tumor IFNγ transcriptional response signature levels with terminal tumor volume. Spearman’s ρ and p-value are shown in the figure.

To test whether therapeutic type 2-to-type 1 repolarization occurred elsewhere, we examined the same parameters in the TDLN and spleen. T_H_1 were largely undetectable in TDLN by flow cytometry, while T_H_2 frequencies decreased in the αIL-4/IL-13/TSLP and αIL-4/IL-13/TSLP/PD1 treated groups and were positively correlated with tumor volume (Sup. Fig. 18A-C). Although TDLN T_H_1 were largely undetectable, bulk RNAseq demonstrated that αIL-4/IL-13/TSLP/PD1 increased IFNγ transcriptional response levels but these were not correlated with tumor volumes (Sup. Fig. 18D-E). αIL-4/IL-13/TSLP/PD1 treatment increased the splenic T_H_1 frequencies, which were inversely correlated with tumor volume (Sup. Fig. 19A-B). The frequency of splenic T_H_2 was decreased by αIL-4/IL-13/TSLP, but splenic T_H_2 frequencies were not significantly correlated with tumor volume (Sup. Fig. 19C-D). This translated to elevated T_H_1/T_H_2 ratios for both groups which was inversely correlated with tumor volumes (Sup. Fig. 19E-F). There were no changes in the IFNγ or IL-4/IL-13 response signature levels in the αIL-4/IL-13/TSLP or αIL-4/IL-13/TSLP/PD1 groups, but both were inversely correlated with tumor volumes (Sup. Fig. 19G-J). Collectively, these data indicated that T_H_2 suppressed anti-tumor immunity in TDLN and that expanded type 1 polarized immune cells need to exit the TDLN for therapeutic efficacy.

We also tested for differences in dupilumab and tezepelumab pharmacodynamic serum biomarkers and type 1 associated chemokines(75–78). CCL11 appeared to be the only specific biomarker as it was unaffected by the isotype and αPD1 treatments but decreased in all groups containing αIL-4/IL-13 (Sup. Fig. 20A-J). Neither CXCL9 nor CXCL10, both associated with type 1 skewing, were significantly affected by any treatment (Sup. Fig. 20K-L). Thus, therapeutic TGI by αIL-4/IL-13/TSLP and αIL-4/IL-13/TSLP/PD1 was associated with reduced type 2 skewing without systemic type 1 rebound inflammation.

### Expanded effector-like CD8 T cells and elevated CD8 T cell Perforin and CD38 expression are associated with therapeutic αIL-4/IL-13/TSLP/PD1 efficacy

Expansion of CD8 T cells in the TME (Fig. 3) and spleen (Sup. Fig. 14) led us to test whether αIL-4/IL-13/TSLP and αIL-4/IL-13/TSLP/PD1 altered CD8 T cell subsets or protein expression. Using expression levels of the markers CD38, Tbet, MHC-II, Ly6C, TIM-3, CD44, SlamF6, Gata3 and Perforin for PaCMAP dimensionality reduction and PARC clustering we identified 17 tumor-infiltrating CD8 T cell clusters (Fig. 5A). Cluster CD8-09 was enriched in αIL-4/IL-13/TSLP-treated tumors, while cluster CD8-06 was only expanded in tumors treated with αIL-4/IL-13/TSLP/PD1 (Fig. 5B-C). Tumor-infiltrating counts of CD8-06 were negatively correlated with tumor volume, whereas counts of CD8-09 were not (Fig. 5D-E). Clusters CD8-09 and CD8-06 were largely differentiated by their levels of SlamF6, TIM-3, Ly6C, CD38 and Perforin, with cluster CD8-06 appearing more cytotoxic effector-like and CD8-09 more stem-memory-like (Fig. 5F, Sup. Fig. 21). Concordant with the increase in CD8-06 cells, total tumor-infiltrating CD8 T cell expression of Perforin and CD38 was higher in the αIL-4/IL-13/TSLP/PD1 group and inversely correlated with tumor volumes (Fig. 5G-J). Recombinant human IL-4 inhibited Perforin induction in primary human CD8 T cells, which was reversed by antibody blockade thus demonstrating that this mechanism crosses the species barrier (Sup. Fig. 22).

**Figure 5.**
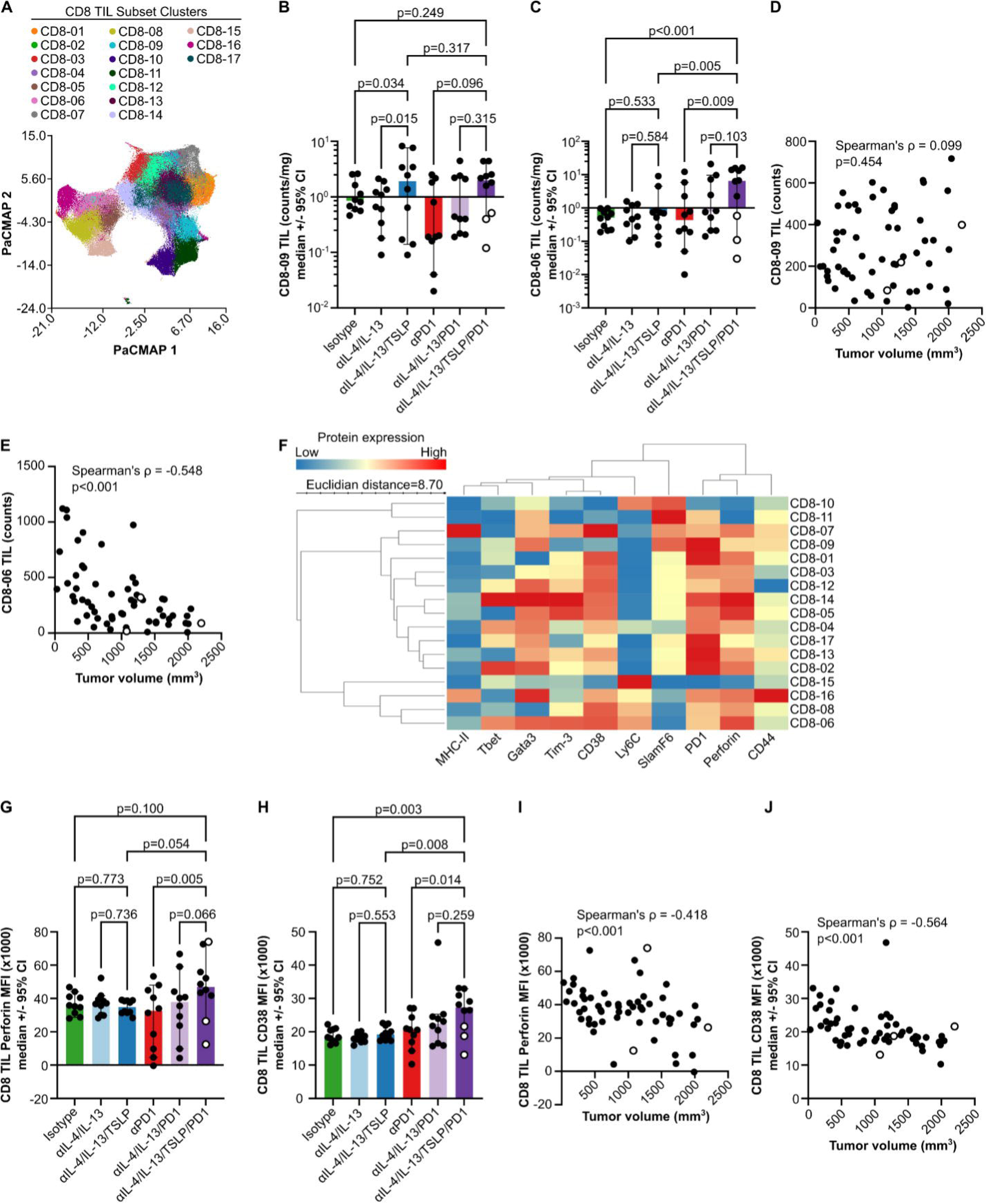
Therapeutic tumor growth inhibition was associated with elevated tumor infiltrating effector-like CD8 T cells. **(A)** PaCMAP dimensionality reduction and PARC clustering of tumor infiltrating (TIL) CD8 T cell subsets using protein expression of CD38, Tbet, MHC-II, Ly6C, Tim-3, CD44, SlamF6, Gata3 and Perforin as measured by flow cytometry. **(B-C)** Mass-normalized counts of tumor infiltrating CD8 clusters -09 and -06 from (A). One-way ANOVAs (CD8-09 p=0.075, CD8-06 p=0.006) followed by Fisher’s uncorrected LSD (p-values show in figure) were used. **(D-E)** Correlations of tumor infiltrating counts of CD8-09 and CD8-06 subsets with terminal tumor volumes. **(F)** Heatmap showing the scaled expression of each protein used in the dimensionality reduction and clustering in (A) across tumor infiltrating CD8 T cell subsets**. (G-H)** Median fluorescence intensities (MFI) of Perforin or CD38 protein expression in total tumor infiltrating CD8 T cell populations as measured by flow cytometry. One-way ANOVA (Perforin p=0.122, CD38 p=0.009) followed by uncorrected Fisher’s LSD tests (p-values shown in figure) were used. **(I-J)** Correlations of total tumor infiltrating CD8 T cell Perforin or CD38 MFI with terminal tumor volumes. Spearman’s ρ and p-values shown in figure.

Performing similar analyses in the TDLN yielded 22 CD8 T cell clusters (Sup. Fig. 23A). Frequencies of six clusters were increased in the αIL-4/IL-13/TSLP/PD1 group with clusters TDLN CD8-07 and -10 showing the largest differences between the αPD1 and αIL-4/IL-13/TSLP/PD1 groups (Sup. Fig. 23B-G). Comparing protein expression levels between these groups demonstrated that TDLN CD8-07 is more stem-memory-like while TDLN CD8-10 is more effector-like (Sup. Fig. 23H). However, none of these frequencies were correlated with tumor volumes (Sup. Fig. 23I-N). Similar analyses on the spleen identified 21 CD8 T cell clusters (Sup. Fig. 24A). Of these, frequencies of six clusters were increased in the αIL-4/IL-13/TSLP/PD1 group (Sup. Fig. 24B-G). Three of these clusters, Spleen CD8-01, Spleen CD8-07 and Spleen CD8-11, were inversely correlated with tumor volumes (Sup. Fig. 24H-J). These three splenic CD8 T cell clusters had higher Perforin and CD44 expression, and lower PD1 levels, than the three that were not correlated with tumor volumes (Sup. Fig. 24K). These data indicate that the enhanced generation of effector memory-like CD8 T cells in secondary lymphoid organs and their increased infiltration into the TME may partially explain the therapeutic TGI seen with αIL-4/IL-13/TSLP/PD1.

### Mature tumor-infiltrating CD40+OX40L-DC2 and multiple cDC1 states were associated with therapeutic TGI by αIL-4/IL-13/TSLP/PD1 treatment

Given the increased tumor-infiltrating DC2 seen with αIL-4/IL-13/TSLP/PD1 treatment, we next looked for alterations in DC2 activation states. We subsampled DC2 from the total CD45+ population, performed PaCMAP dimensionality reduction and clustering with PARC. Using expression levels of PD-L1, CD86, SIRPα, MHC-II, Tim-3, OX40L and CD40 we identified 21 clusters of tumor-infiltrating DC2 (Fig. 6A). Six of these clusters were increased in the αIL-4/IL-13/TSLP/PD1 group (Sup. Fig. 25). Four of these clusters were inversely correlated with tumor volumes (Fig. 6B-E). These four clusters were largely distinguished by higher MHC-II and lower OX40L expression (Fig. 6F). Consistent with this observation, total tumor-infiltrating DC2 expression of OX40L was positively correlated with tumor volumes while MHC-II, CD40 and SIRPα were inversely correlated (Fig. 6G-J). Collectively, these changes in DC2 activation states in the TME may drive the differential reactivation of tumor-infiltrating T_H_1 and T_H_2 associated with αIL-4/IL-13/TSLP/PD1-driven TGI.

**Figure 6.**
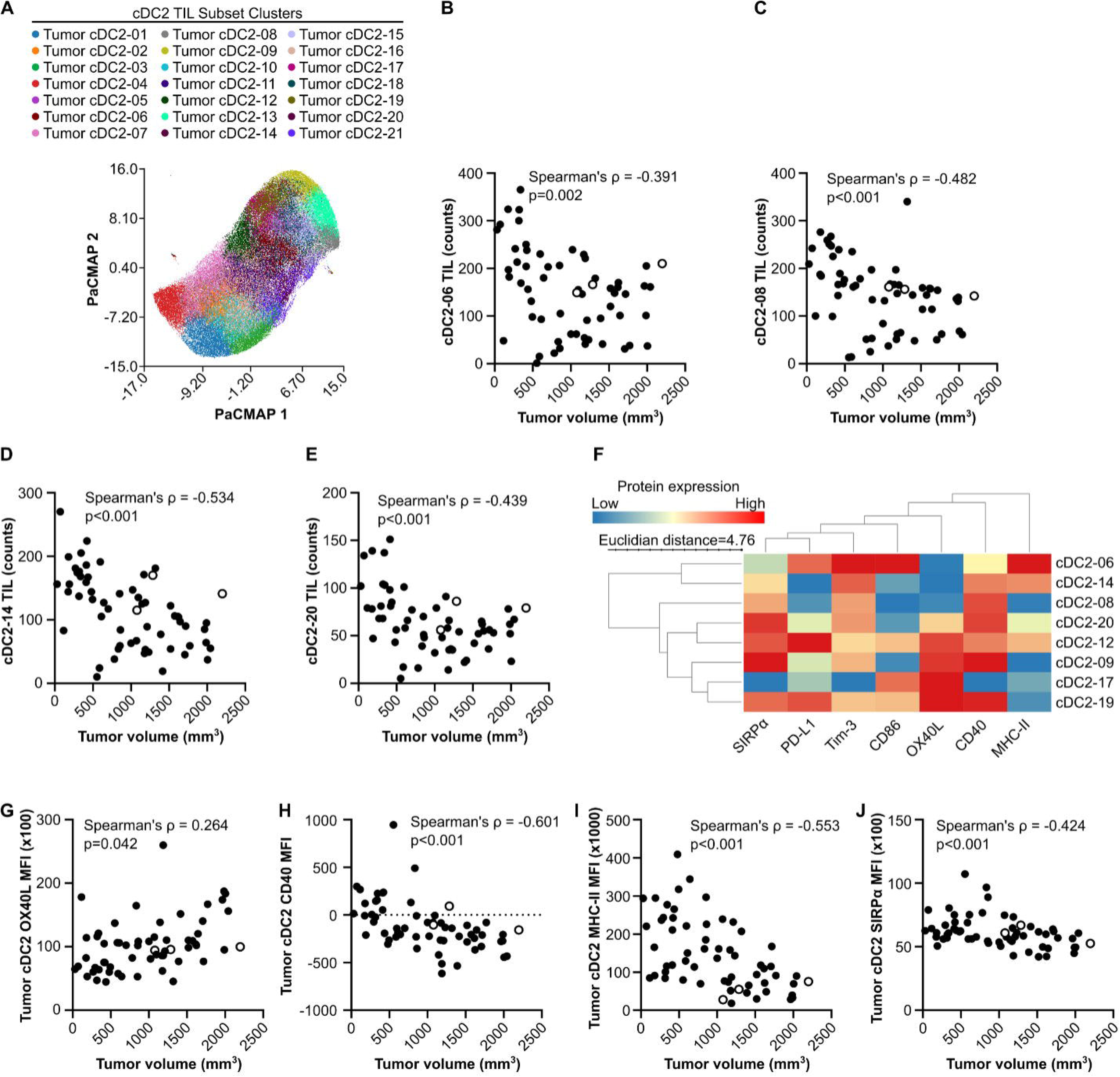
Therapeutic tumor growth inhibition was associated with elevated cDC2 subsets expressing CD40, MHC-II and SIRPα. **(A)** PaCMAP dimensionality reduction and PARC clustering of tumor infiltrating (TIL) cDC2 subsets using protein expression of SIRPα, PD-L1, Tim-3, CD86, OX40L, CD40 and MHC-II as measured by flow cytometry. **(B-E)** Correlations of tumor infiltrating counts of cDC2-06, -08, -14 and -20 subsets with terminal tumor volumes. **(F)** Heatmap showing the scaled expression of each protein used in the dimensionality reduction and clustering in (A) across selected tumor infiltrating cDC2 subsets. **(G-J)** Correlations of total tumor infiltrating cDC2 OX40L, CD40, MHC-II or SIRPα MFI with terminal tumor volumes. Spearman’s ρ and p-values shown in figure.

As classical monocytes are potential precursors for DC2, the increase in tumor-infiltrating DC2 (Fig. 3G) may have been due to increased tumor infiltrating classical monocytes (Fig. 3D) or greater monocyte-to-DC2 differentiation. PaCMAP dimensionality reduction and PARC clustering on subsampled tumor infiltrating classical monocytes identified 15 clusters, five of which were increased in the αIL-4/IL-13/TSLP/PD1 group (Sup. Fig. 26A-F). Three of these clusters were distinguished by higher CD11c, SIRPα, MHC-II and CD86 expression, and higher numbers of these monocytes subsets in the TME were correlated with smaller tumor volumes (Sup. Fig. 26G-J). Consistent with these data, expression of CD11c, MHC-II, CD40 and CD86 by total tumor-infiltrating classical monocytes was inversely correlated with tumor volumes (Sup. Fig. 26K-N). These data suggest that increased classical monocyte infiltration and differentiation into DC2 may have driven therapeutic TGI by the αIL-4/IL-13/TSLP/PD1 treatment.

Although the number of tumor-infiltrating cDC1 did not change with treatment, differences in cDC1 activation states may have driven the changes in TME CD8 T cell phenotypes. To address this, we subsampled cDC1 from the total tumor CD45+ population and performed PaCMAP dimensionality reduction and PARC clustering using the same markers as for cDC2. This approach identified 18 cDC1 clusters, six of which were increased in the αIL-4/IL-13/TSLP/PD1 treated group and three of these were inversely correlated with tumor volumes (Sup. Fig. 27A-J). The three clusters that were inversely correlated with tumor volumes had heterogeneous protein expression, but total cDC1 expression of PD-L1 and CD86 had the strongest inverse correlation while Tim-3 and OX40L had the strongest positive correlation with tumor volumes (Sup. Fig. 27K-O). Thus, changes in TME cDC1 activation states associated with T cell reactivation or priming may be partly responsible for the increase in cytotoxic effector-like CD8 TIL and the therapeutic TGI driven by αIL-4/IL-13/TSLP/PD1.

### High IL-4/IL-13, TSLP and T_H_2 transcriptional signatures identify a subset of tumors and are broadly associated with poor overall survival outcomes

To identify patient populations that might benefit from αIL-4/IL-13/TSLP/PD1 therapy, we asked whether these cytokines are associated with poor survival outcomes in human cancer patients. First, we quantified transcripts for IL-4, IL-13 and TSLP in bulk RNAseq data from a variety of tumor types using The Cancer Genome Atlas (TCGA) data. Lung squamous cell (LUSC), head and neck squamous cell (HNSC) and esophageal carcinoma (ESCA) had the highest combined levels of these transcripts, while diffuse large B cell lymphoma (DLBC) and thymoma (THYM) had the highest IL-4 RNA (Fig. 7A). Across the whole TCGA cohort, all three transcripts were associated with worse overall survival with IL-4 having the highest hazard ratio (Fig. 7B-D). To identify tumor types with active responses to these cytokines, we quantified levels of transcriptional response signatures to IL-4/IL-13 and TSLP and the transcriptional T_H_2 deconvolution signature in these data. DLBC, pancreatic adenocarcinoma (PAAD), stomach adenocarcinoma (STAD), LUSC and HNSC had the highest combined IL-4/IL-13 and TSLP response signature expression (Fig. 7E). Levels of all three response and deconvolution signatures were associated with poor overall survival in the total TCGA cohort (Fig. 7F-H). Together, these data indicate that active responses to IL-4, IL-13 and TSLP are associated with poor overall survival across multiple patient cohorts.

**Figure 7.**
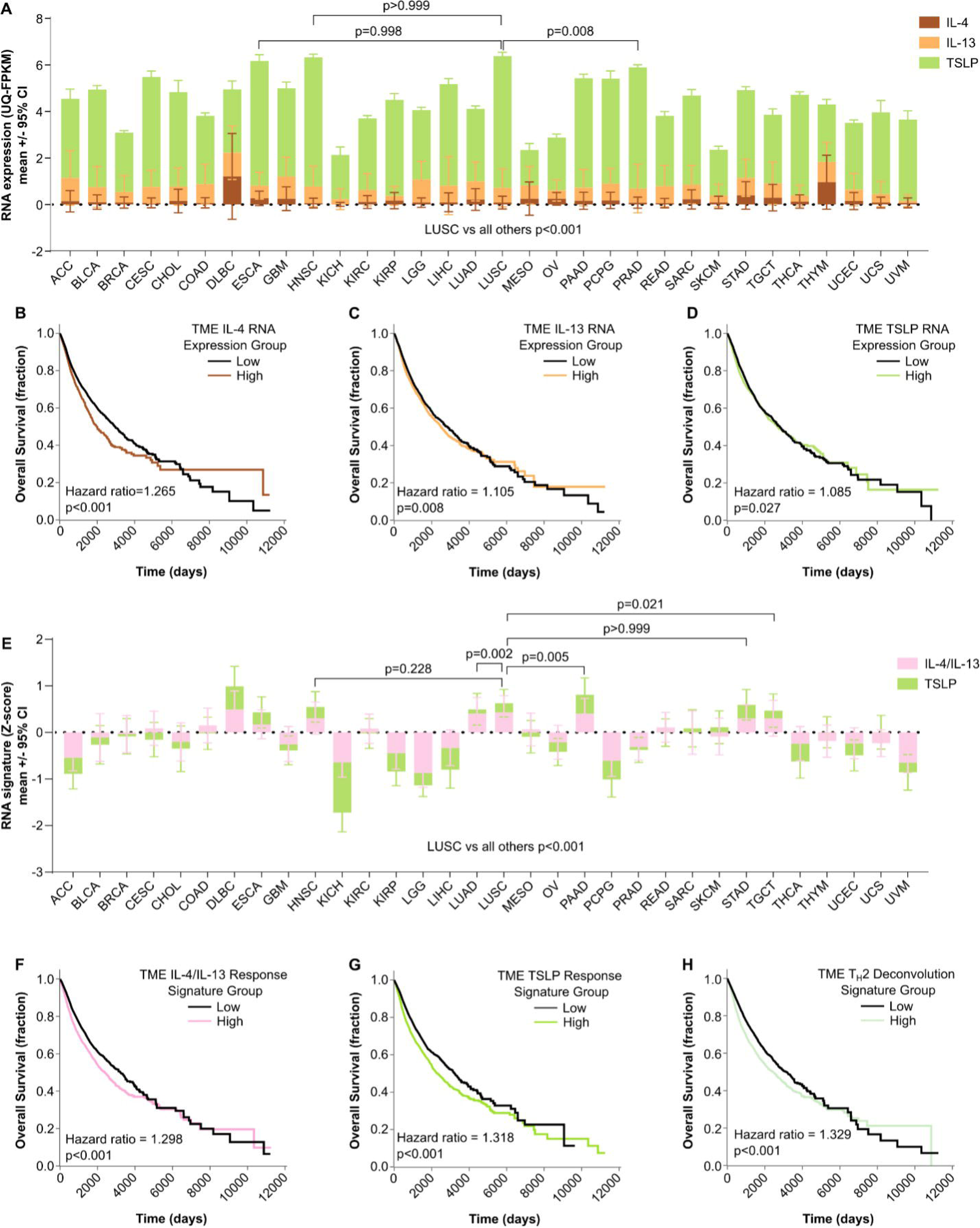
Transcriptional response signatures to IL-4, IL-13 and TSLP and a transcriptional T_H_2 deconvolution signature are associated with worse overall survival outcomes across multiple tumor types. **(A)** Upper quartile normalized fragments per kilobase of transcript per million mapped fragments (UQ-FPKM) from total tumor bulk RNAseq for IL-4 (brown), IL-13 (orange) or TSLP (lime green) by tumor type in The Cancer Genome Atlas (TCGA) cohorts. Two-way ANOVA (TCGA cohort p<0.001 and cytokine transcript p<0.001) followed by Dunnett’s multiple comparisons test (p-values for LUSC shown in figure) were used. **(B-D)** Associations between median-split tumor RNA expression of IL-4 (B), IL-13 (C) or TSLP (D) and overall survival in the total TCGA cohort regardless of tumor type. **(E)** Z-scored IL-4/IL-13 (pink) or TSLP (lime green) transcriptional response signatures from the same data in (A). Two-way ANOVA (TCGA cohort p<0.001 and signature p=0.523) followed by Dunnett’s multiple comparisons tests (p-values for LUSC shown in figure) were used. (F-H) Associations between IL-4/IL-13 (F) or TSLP (G) transcriptional signatures or T_H_2 (H) deconvolution signature and overall survival in the total TCGA cohort regardless of tumor type. Cox proportional hazard ratios and p-values for differences between the low and high expression groups are shown in the figure.

Examining the overall survival hazard ratios for each type of cancer across cytokine transcripts, response signature and cell deconvolution signature levels identified four classes of outcome associations. First, those with high hazard ratios for both IL-4/IL-13 and TSLP transcriptional responses which included testicular germ cell cancer (TGCT), papillary kidney carcinoma (KIRP), uveal melanoma (UVM) and STAD (Sup. Table 5). Second, tumors with high hazard ratios to only the IL-4/IL-13 transcriptional response signature including low-grade glioma (LGG), THYM, chromophobe renal cell carcinoma (KICH), glioblastoma multiforme (GBM) and LUSC (Sup. Table 5). Third, tumors with high hazard ratios to only the TSLP transcriptional response signature such as prostate adenocarcinoma (PRAD), papillary thyroid carcinoma (THCA), bladder urothelial carcinoma (BLCA), liver hepatocellular carcinoma (LIHC) and lung adenocarcinoma (LUAD) (Sup. Table 5). Fourth, cancers with low hazard ratios to both transcriptional response signatures, which included cutaneous melanoma (SKCM), breast cancer (BRCA), DLBC, rectum adenocarcinoma (READ) and PAAD (Sup. Table 5). These data demonstrate the broad potential applicability of trispecific antibodies blocking IL-4, IL-13 and TSLP in multiple solid tumor patient populations including those where checkpoint inhibition is the standard of care.

## Discussion

In this study we demonstrated that therapeutic α-IL-4/IL-13/TSLP treatment improved TGI. Although we did not prove what caused this TGI, CT26 tumors in our α-IL-4/IL-13/TSLP group appeared “hotter” than those of the isotype, α-IL-4/IL-13 and α-PD1 mice. This included higher TME densities of T_H_1, T_H_2 and CD8 T cells, which is consistent with the effects of α-IL-4 in several tumor models(12,14,36). It is likely that the increased tumor infiltrating T_H_1 and T_H_2 arose largely from local proliferation as their frequencies in the TDLN and spleen were unchanged or reduced. Given the increase in splenic stem-memory CD8 T cells in the α-IL-4/IL-13/TSLP group, these likely contributed to the higher density of CD8 T cells in the TME. Splenic stem-memory CD8 T cells expressing Tcf1 also arise during viral infection(79). Given the importance of stem-memory CD8 T cells expressing Tcf1 or SlamF6 to anti-tumor immunity, the splenic stem-memory CD8 T cells in α-IL-4/IL-13/TSLP mice likely fed fresh effector CD8 T cells to the TME to maintain TGI like a previous observation in the TDLN (7,80–83).

The additive TGI of α-IL-4/IL-13/TSLP/PD1 treatment was also associated with increases in stem memory-like CD8 T cells in SLO and the TME. In our study, neither increased frequencies of stem memory-like CD8 T cell subsets nor IFNγ transcriptional response in TDLN were correlated with tumor volumes. However, expanded stem memory-like CD8 T cells in the spleen, which represents both circulation and local priming, were correlated with better TGI. This expansion of stem memory-like CD8 T cells in SLO was likely due to blocking PD1 on newly activated naïve T cells from interacting with PD-L1 or PD-L2 on DCs(84–89). α-IL-4/IL-13/TSLP/PD1 also increased the density of stem memory-like CD8 T cells in the TME. Although this increase was not associated with better TGI the expanded pool of stem memory-like CD8 T cells in the TME likely fed the higher densities of effector CD8 T cells. Blocking IL-4 during CD8 T cell priming directly improves their Perforin- and Granzyme-mediated cytotoxicity and IFNγ secretion(20,21,23,24). Thus, combined blockade of IL-4 and PD1 was likely responsible for the greater numbers of effector-like CD8 T cells and increased TME IFNγ transcriptional response in the α-IL-4/IL-13/TSLP/PD1 group. Both the higher density of effector-like CD8 T cells and increased TME IFNγ transcriptional response correlated with smaller tumors suggesting one potential mechanism of action for α-IL-4/IL-13/TSLP/PD1. Our data is consistent with observations that anti-tumor stem memory CD8 T cells primed in SLO are necessary but must extravasate in or around the tumor then be reactivated by DCs in the TME for full effector differentiation and TGI(90).

Improved reactivation and effector differentiation of tumor-infiltrating CD8 T cells in our α-IL-4/IL-13/TSLP/PD1 group may have been driven by repolarized cDC2- or monocyte-derived DCs (moDC). Higher densities of OX40L-low, CD40+ and MHC-II high cDC2/moDC and monocytes expressing cDC2 markers were associated with α-IL-4/IL-13/TSLP/PD1-induced TGI. Inhibiting IL-4, IL-13 and TSLP activates a DC program including IL-12 and CD40 and dampens the TSLP-associated DC program that includes OX40L, thus improving the priming of anti-tumor T cells(12,42,57,59,63,64). PD-L1 expression, at least by cDC1, is independent of IL-4(12). Both MHC-II and PD-L1 can be stimulated by IFNγ, and increased IFNγ transcriptional responses were only observed in TME where IL-4, IL-13 and PD1 were blocked. Thus, the reprogramming of TME cDC2/moDC appeared to result from combined blockade of IL-4, IL-13 and TSLP which acted orthogonally to α-PD1.

In addition to repolarizing TME cDC2/moDC, α-IL-4/IL-13/TSLP/PD1 also increased the density of these cells in CT26 tumors. These two types of DCs share expression of several markers used for flow cytometric identification in our study but although we did not prove that the expanded cDC2 were monocyte derived, this seems likely given prior observations(91–96). If the expanded cDC2/moDC seen upon α-IL-4/IL-13/TSLP/PD1 treatment are monocyte derived this would partially reconcile our findings with those of LaMarche et al(14). Their group demonstrated that improved TGI in several metastatic lung tumor models was due to deletion of IL-4 and IL-13 signaling in Ms4a3-cre expressing monocyte precursors, but not pre-DC derived Zbtb46-expressing cells(14). Cells traced by the Ms4a3 mouse line used by LaMarche and colleagues can become cDC2/moDC, and moDC appear to arise independent of pre-DC and Zbtb46(91–96). MoDCs can be potent inducers of IFNγ and can drive cytotoxic control of tumors by CD8 T cells(93,95–97). Thus, increased monocyte differentiation into cDC2/moDC in response to α-IL-4/IL-13/TSLP/PD1 could have resulted in the expanded cDC2/moDC densities and increased effector CD8 T cell differentiation in the TME.

An alternative hypothesis is that repolarization of cDC1 in the TME was responsible for the improved effector-like CD8 T cell differentiation in the α-IL-4/IL-13/TSLP/PD1 group. Although α-IL-4/IL-13/TSLP/PD1 did not increase cDC1 density in the TME, it expanded CD40+ and MHC-II high cDC1 subsets. Expression of CD40 and MHC-II by cDC1 is associated with T_H_1 help and CD8 T cell-mediated TGI(98–102). As IL-4Rα signaling inhibits CD40 expression by cDC1 in the non-small cell lung cancer TME, it is likely that blocking these type 2 cytokines was partially responsible for the TME cDC1 repolarization we observed. Given the importance of PD-L1 and PD-L2 expression by DC to CD8 T cell activation, additional PD1 blockade may have augmented reactivation and effector differentiation of TME infiltrating CD8 T cells(84–89). Together, our data suggests that combined blockade of IL-4, IL-13 and TSLP drove synergistic TGI with PD1 antagonism by creating “hotter” tumors through altering orthogonal programs in CD4 and CD8 T cells in multiple tissues and cDC1, cDC2/moDC and classical monocytes in the TME (Sup. Fig. 28).

Widespread clinical use of dupilumab has led to on-label treatment of patients with coincident cancers with mostly positive outcomes and four clinical trials combining dupilumab with PD1/PD-L1 blockade (NCT03886493, NCT06088771, NCT05967884 and NCT05013450)(14,17,103–107). The beneficial TGI effects we report from blocking IL-4, IL-13, TSLP and PD1 are generally in line with these studies, and with a systematic review which indicated that blocking IL-4 and IL-13 is safe for patients with malignancies(108). Despite IL-4/IL-13 and TSLP transcriptional response signatures being associated with poor survival outcomes across the total TCGA cohort, they were not universally detrimental. For example, in cutaneous melanoma patients higher IL-4/IL-13 and TSLP transcriptional signature levels were associated with better overall survival. Atopic dermatitis and the cutaneous T cell lymphomas mycosis fungoides (MF) and Sézary syndrome have similar manifestations occasionally leading to misdiagnosis and treatment of lymphoma patients with dupilumab(109). Given the mixed clinical outcomes for MF patients treated with dupilumab, a detailed understanding of how type 2 immune polarization affects cancer progression is necessary for safe and effective use of these drugs(110,111).

## Figure Legends

**Supplemental Figure 1.**
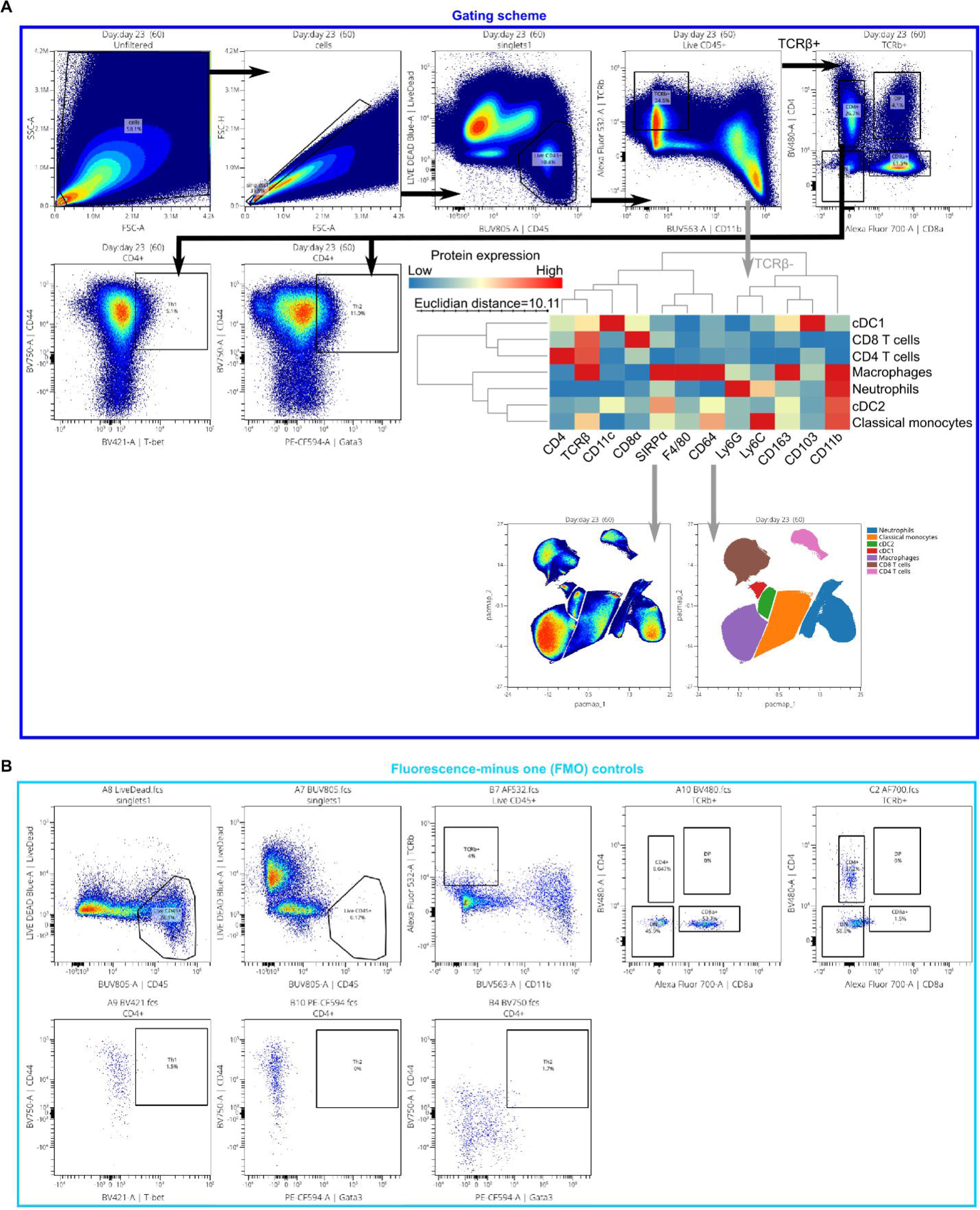
Traditional 2-dimensional gating scheme and fluorescence minus one (FMO) controls for CT26 tumor flow cytometry. **(A)** Traditional 2-dimensional gating scheme used for CT26 tumor, spleen and tumor draining lymph node samples. Images show concatenations of all tumor samples. Gating hierarchy follows the arrows to the “Live CD45+” gate at which point the black arrows denote gates from the TCRβ+ population and gray arrows denote gates from the TCRβ-population. For the TCRβ-population, expression levels of the markers shown in the heatmap viewed on PaCMAP dimensionality reduction plots were used to draw gates directly on the plot for the indicated populations. The union gate and cluster annotations are shown below the heatmap. Subsequent analyses were conducted using subsampling of each gated population. **(B)** FMO controls for each of the gated populations. The population names “singlets1”, “Live CD45+”, “TCRb+” and “CD4+” match the analogous graphs in (A).

**Supplemental Figure 2.**
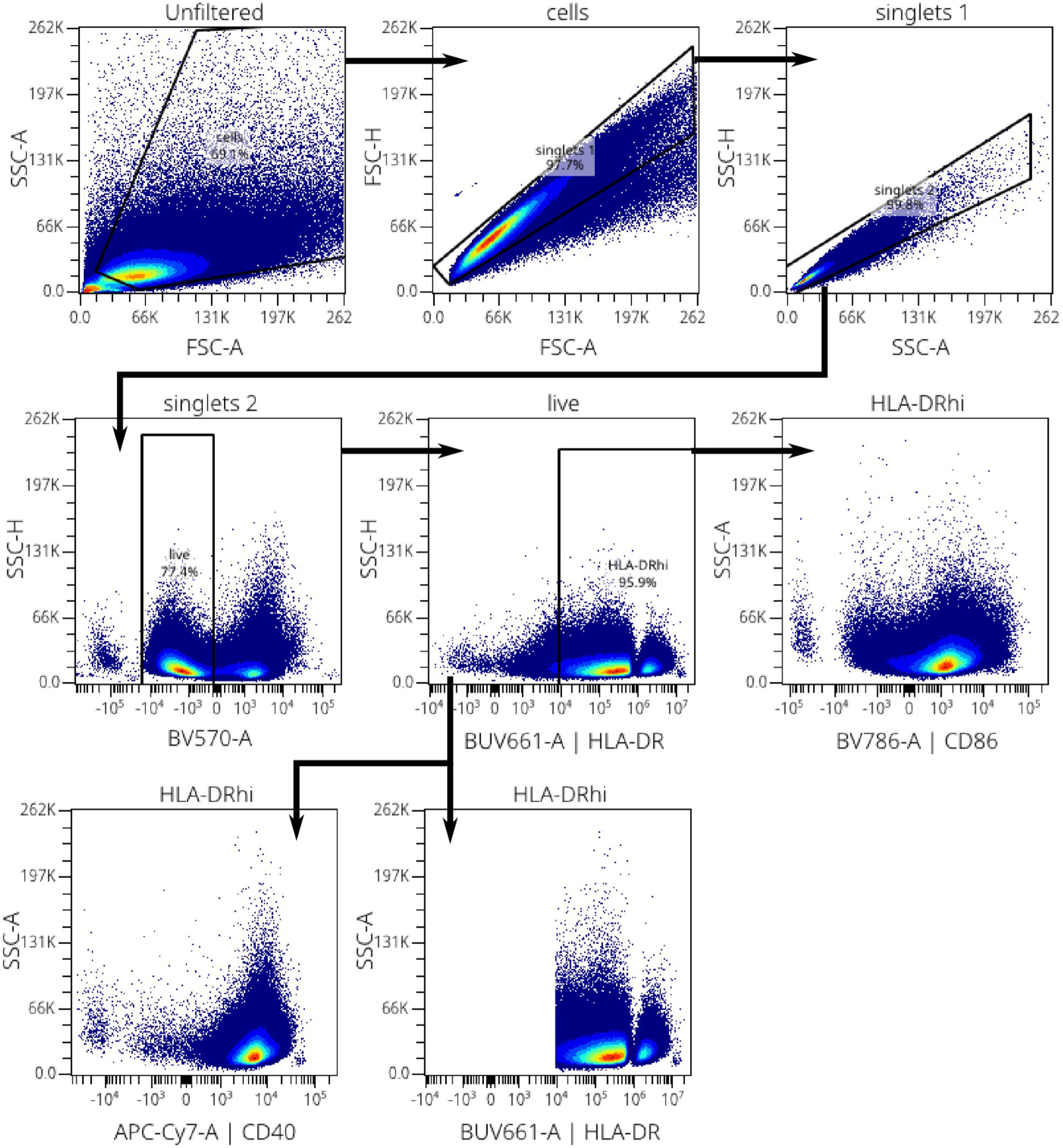
Flow cytometry gating strategy for human monocyte-derived macrophages. Gating hierarchy for quantifying CD86, CD40 and MHC-II MFI on human monocyte-derived macrophages (MDM). Arrows denote the sequential gating hierarchy. CD86, CD40 and MHC-II expression was measured as the median fluorescence intensity of cells within the live HLA-DR+ gate. The titles at the top of each graph denote the parental gate.

**Supplemental Figure 3.**
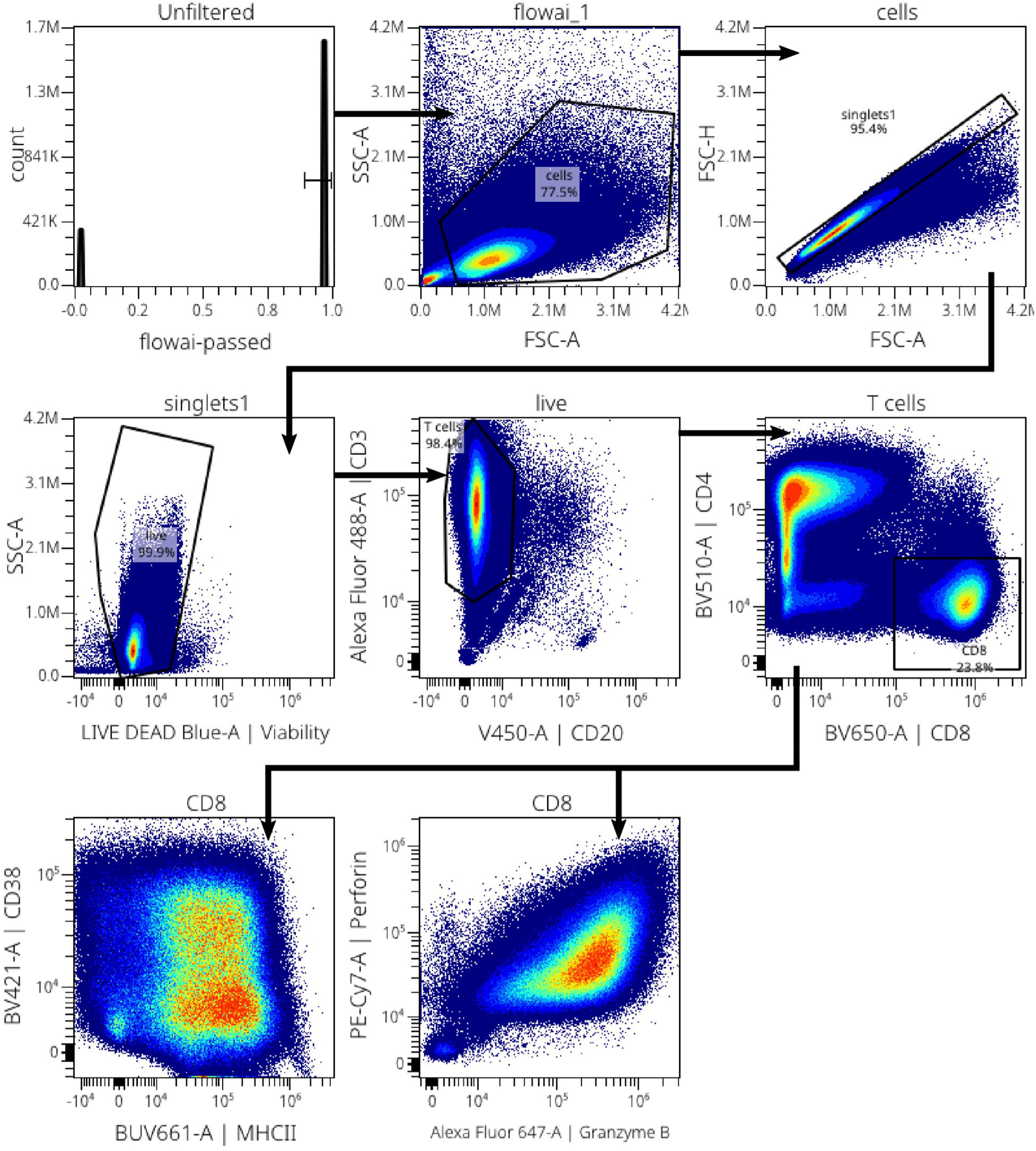
Flow cytometry gating scheme for human T cells. Flow cytometry gating scheme used for quantifying CD38 and Perforin expression using median fluorescence intensity (MFI) by *in vitro* activated primary human CD8 T cells. Plots show concatenated data from all the samples acquired in a single biological replicate experiment.

**Supplemental Figure 4.**
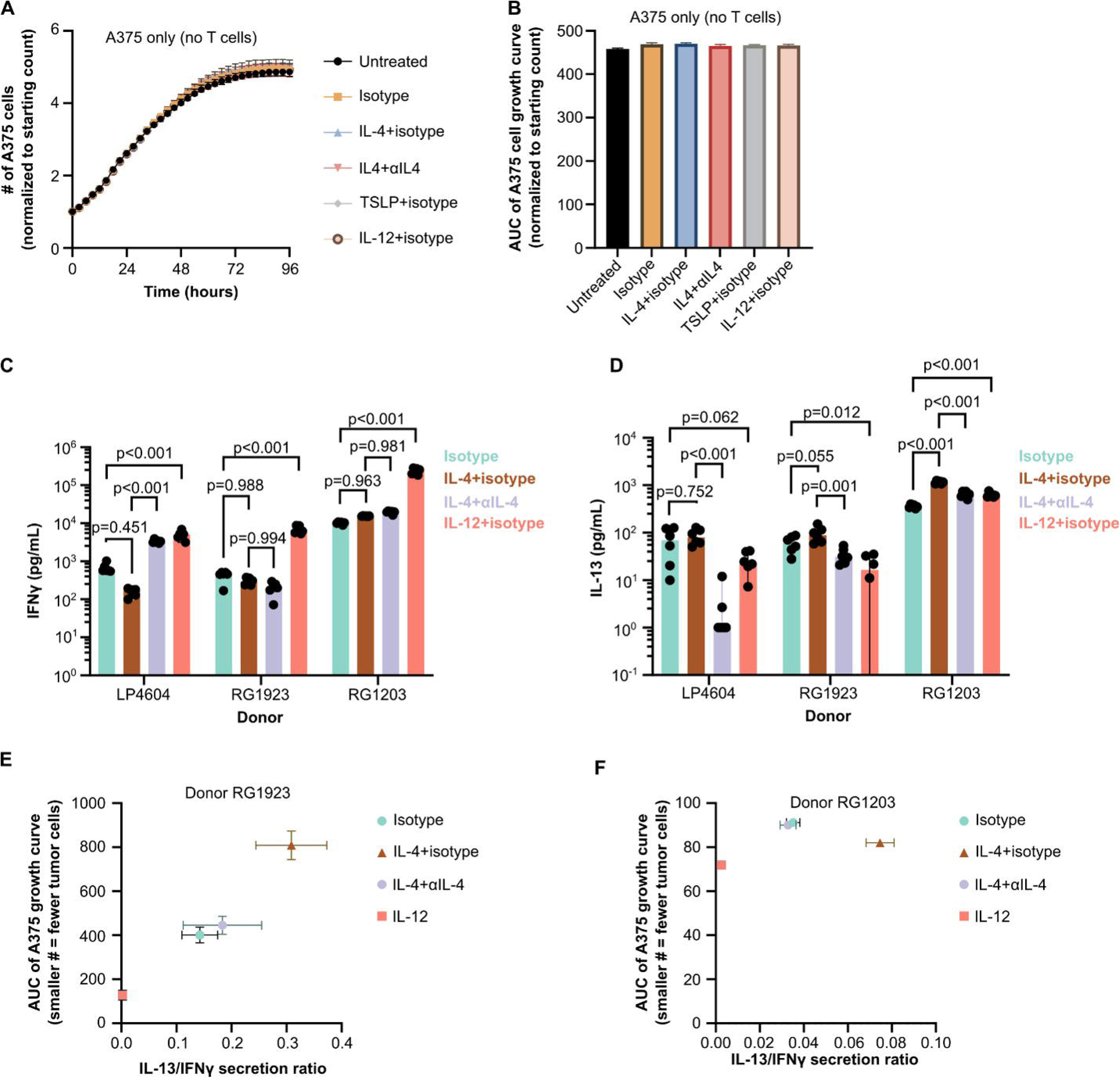
IL-4 inhibits TGI by acting on T cells, not A375 cells. **(A)** A375 melanoma cell line growth curves in the presence of cytokines and blocking antibodies, but in the absence of primary human T cells. **(B)** AUC of the growth curves from (A). **(C-D)** Levels of IFNγ and IL-13 secreted by T cells in the *in vitro* tumor growth inhibition assay related to Fig. 1C-D. Two-way ANOVA for differences between treatments (p<0.001), donors (p<0.001) and the interaction (p<0.001) were followed by Tukey’s multiple comparisons tests (p-values shown in figure). **(E-F)** Correlations of IL-13/IFNγ secretion and tumor growth inhibition AUC for donors RG1923 (E, Spearman’s ρ=0.771, p<0.001) and RG1203 (F, Spearman’s ρ=0.107, p=0.617), related to Fig. 1D.

**Supplemental Figure 5.**
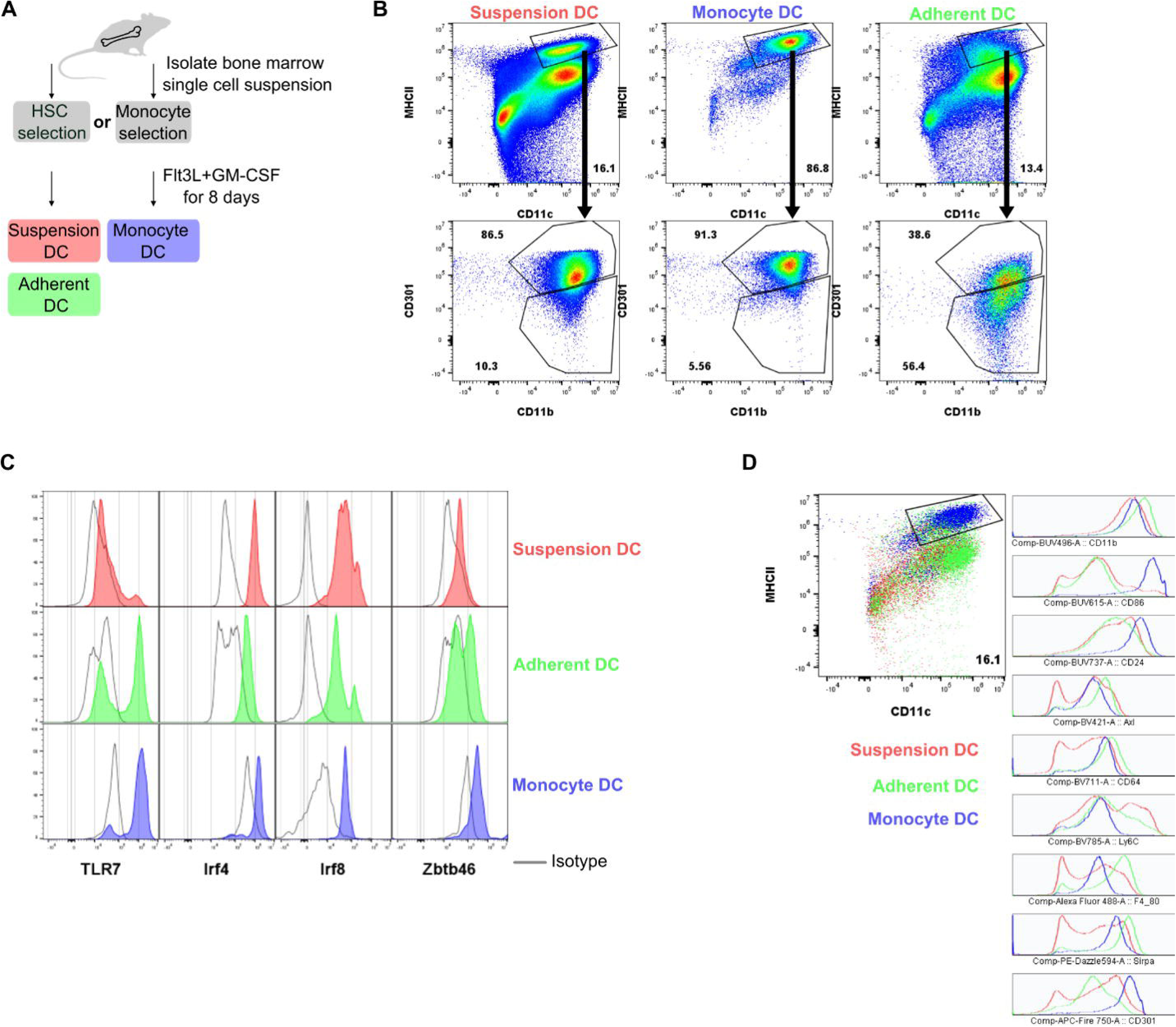
Differentiation and purity of murine bone marrow-derived cDC2-like cells. **(A)** Cartoon of the three murine bone marrow cDC2-like differentiation schemes tested. **(B)** Comparison of the frequency of cDC2-like cells (defined as MHC-II+CD11c+CD301+CD11b+) derived from each differentiation technique as determined by flow cytometry. The numbers on each plot indicate the percentage of cells that fall within the associated gates. **(C)** Expression of factors associated with anti-tumor cDC responses (TLR7), cDC2 (Irf4), cDC1 (Irf8) or total cDC (Zbtb46) in cDC-like cells from the three differentiation methods as measured by flow cytometry. Filled histogram traces show staining for the proteins indicated on the X-axis while the unfilled black line traces represent isotype controls. **(D)** Expression levels of several markers associated with cDC2 (CD24, SIRPα and CD301), macrophages (CD64 and F4/80) and monocytes (Ly6C) by cDC-like cells from the three differentiation methods as measured by flow cytometry.

**Supplemental Figure 6.**
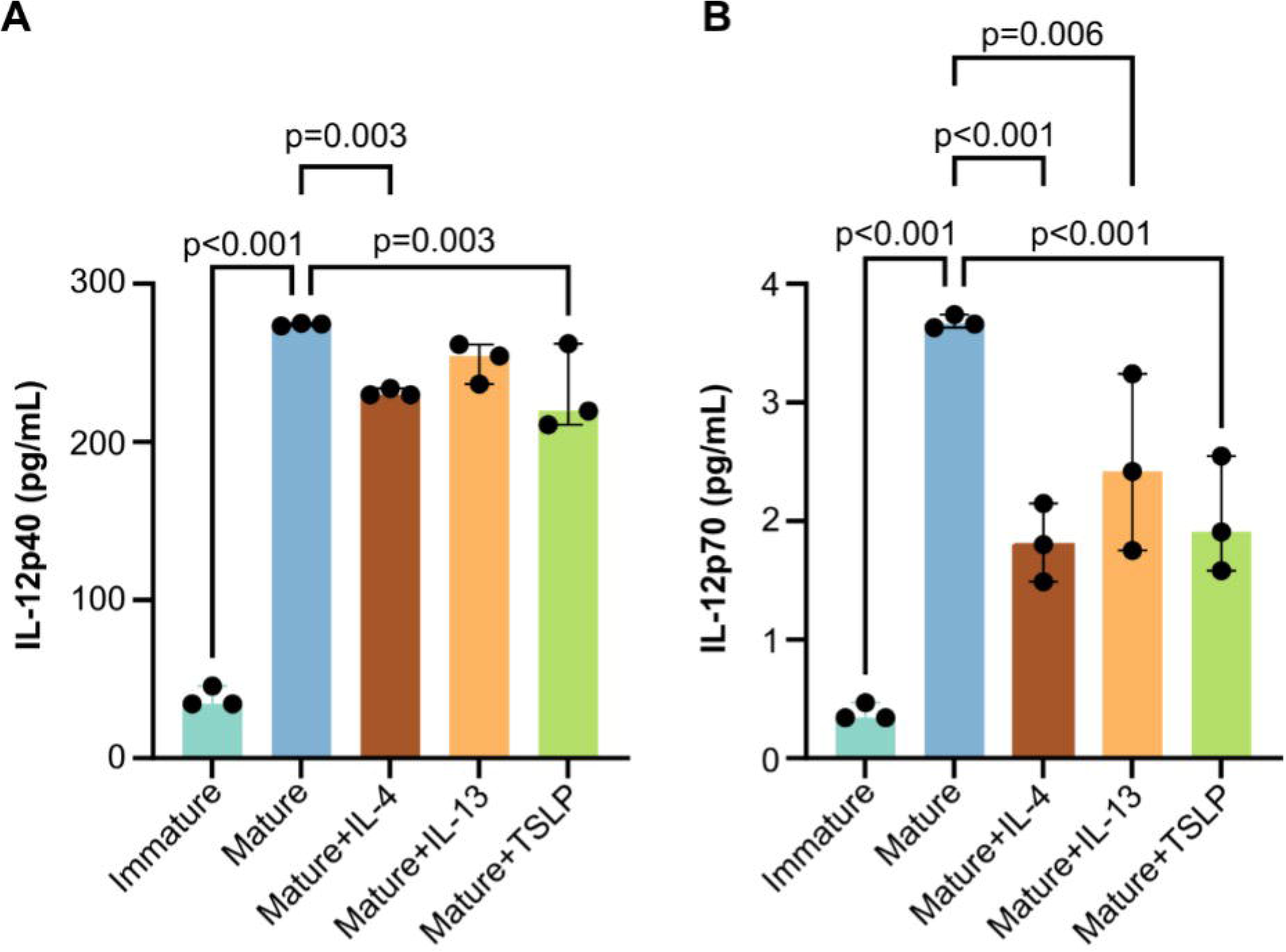
Recombinant human IL-4 and TSLP inhibit maturation-induced secretion of IL-12p40 and IL-12p70 by human monocyte-derived DCs (moDC). (A-B) Secretion of IL-12p40 (A, p<0.001) or IL-12p70 (B, p<0.001) by moDC in response to a maturation cocktail and the recombinant cytokines listed as determined by fluid phase immunoassay. Dots represent biological replicates from one donor performed on different days. Bars represent the medians and whiskers the 95% confidence intervals of each group. One-way ANOVA (p-values listed above) followed by uncorrected Fisher’s LSD tests (p-values shown in figure) were used to test for statistical differences between groups.

**Supplemental Figure 7.**
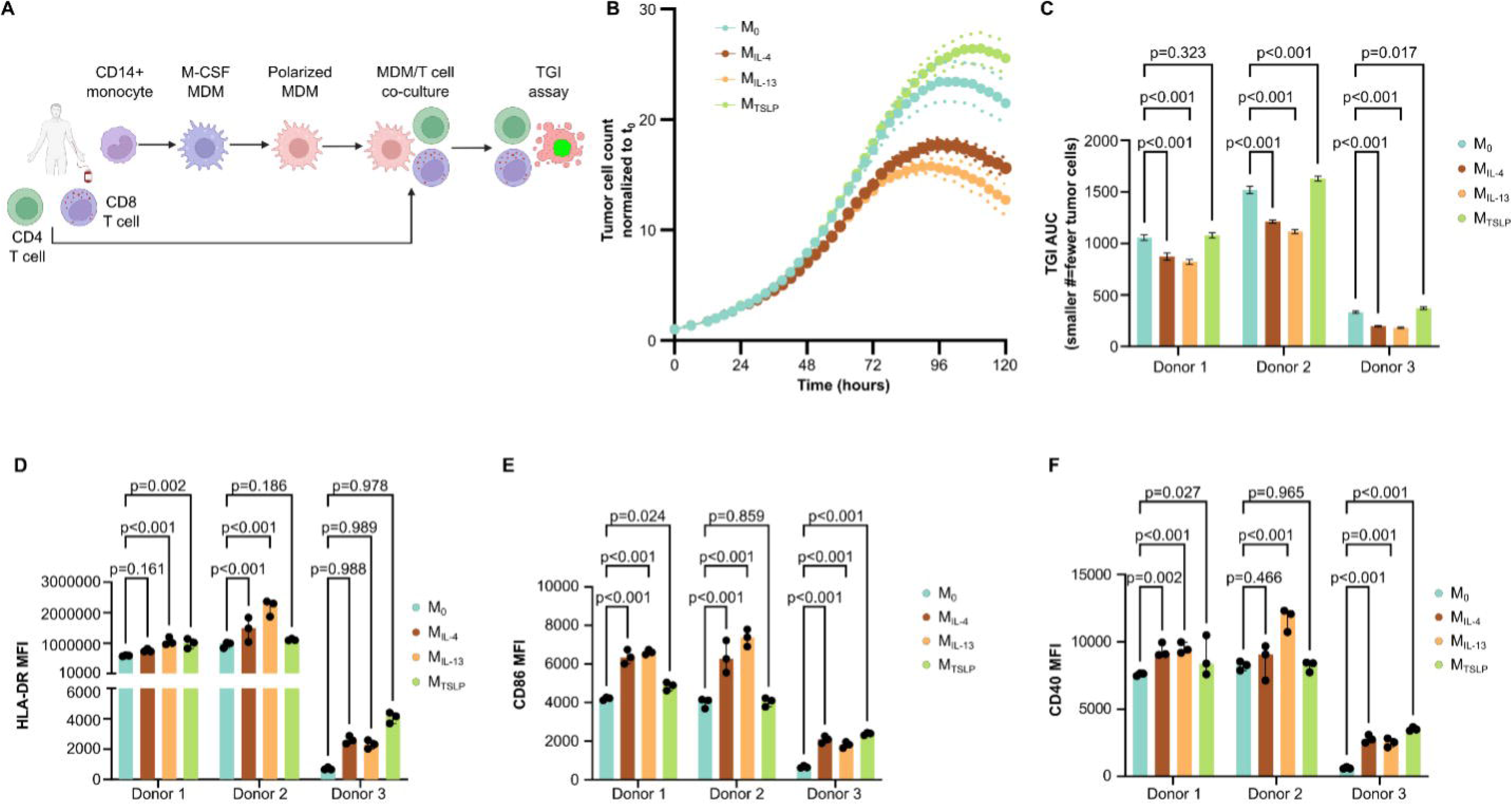
Human monocyte-derived macrophages (MDM) polarized with IL-4 and/or IL-13 enhance T cell-mediated tumor growth inhibition. **(A)** Cartoon describing the assay. Created with BioRender.com. **(B)** A375 growth curve normalized to the number of cells at the initiation of the assay (t_0_). M_0_ are unpolarized MDM, M_IL-4_ were polarized with IL-4, M_IL-13_ were polarized with IL-13, while M_TSLP_ were polarized with TSLP. **(C)** AUC for the growth curves in (B). Two-way ANOVA testing (polarization p<0.001, donor p<0.001 and interaction p<0.001) were followed by Dunnett’s multiple comparisons tests (p-values shown in figure). Six biological replicates were conducted per donor and the bars and whiskers show group means and standard deviations, respectively. **(D-F)** Median fluorescence intensities (MFI) of HLA-DR (D), CD86 (E) and CD40 (F) proteins expressed by the polarized macrophages as measured by flow cytometry. Two-way ANOVAs (polarization p<0.001, donor p<0.001 and interaction p<0.001 for all three proteins) were followed by uncorrected Fisher’s LSD tests (p-values shown in figure). Three biological replicates were conducted for each donor.

**Supplemental Figure 8.**
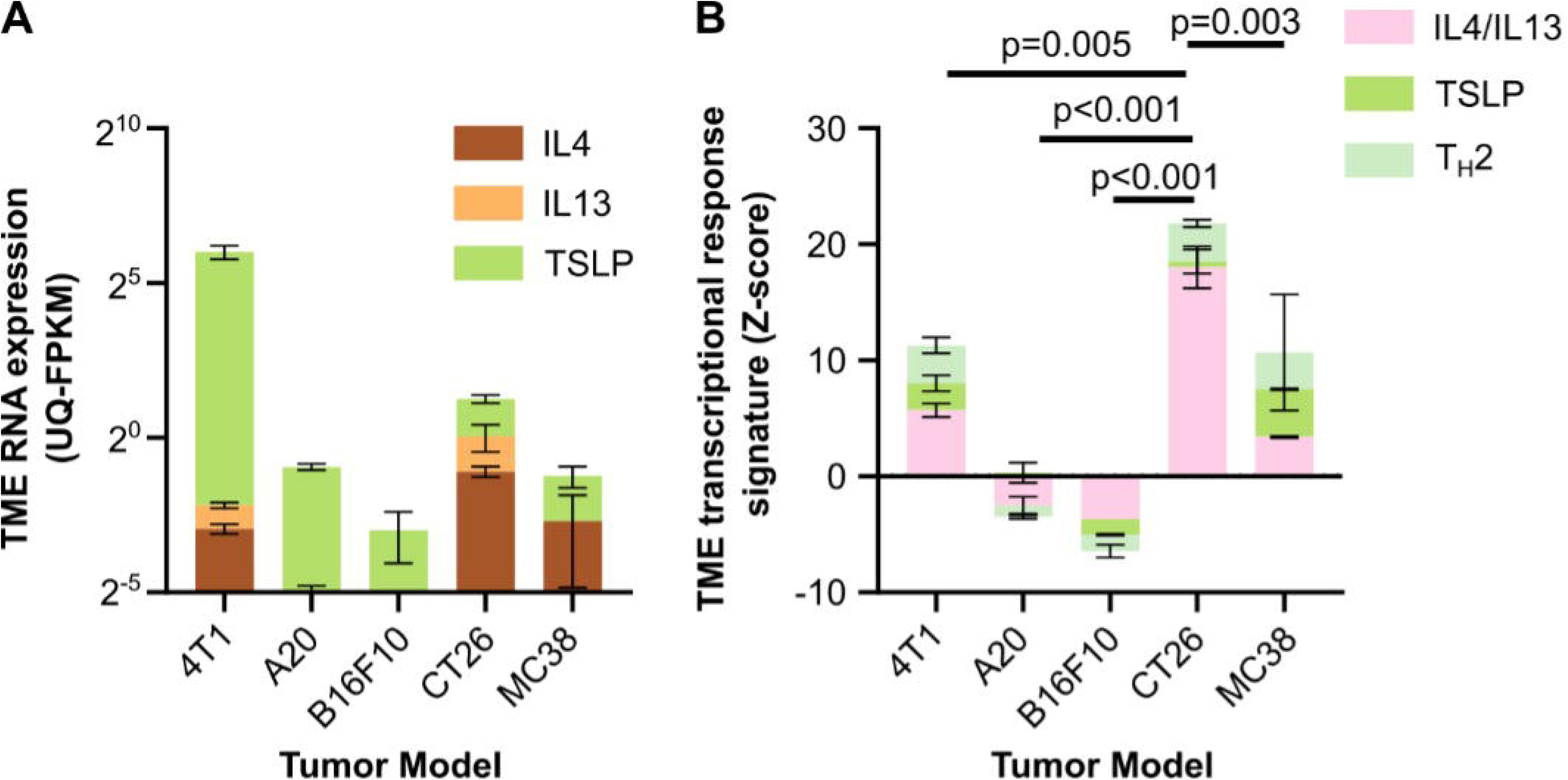
Identification of a murine subcutaneous tumor model with IL-4, IL-13 and TSLP responses. **(A)** Upper quartile normalized fragments per kilobase of transcript per million mapped fragments (UQ-FPKM) of IL-4 (yellow), IL-13 (orange) and TSLP (lime green) from bulk RNA sequencing (RNAseq) of the tumor microenvironments (TME) from the subcutaneous tumor models listed on the X-axis. **(B)** Z-scored transcriptional response signatures for IL-4 and IL-13 (pink) or TSLP (lime green) and a T_H_2 deconvolution signature (seafoam green) from the same bulk RNAseq data as in (A). TME from 3-5 mice were sequenced per model with bars representing the median and the whiskers representing the 95% confidence intervals. Statistical testing in (B) is from a two-way ANOVA (transcriptional signature p=0.001, tumor type p<0.001, interaction p<0.001) followed by Dunnett’s multiple comparison tests versus the CT26 group (p-values shown in figure).

**Supplemental Figure 9.**
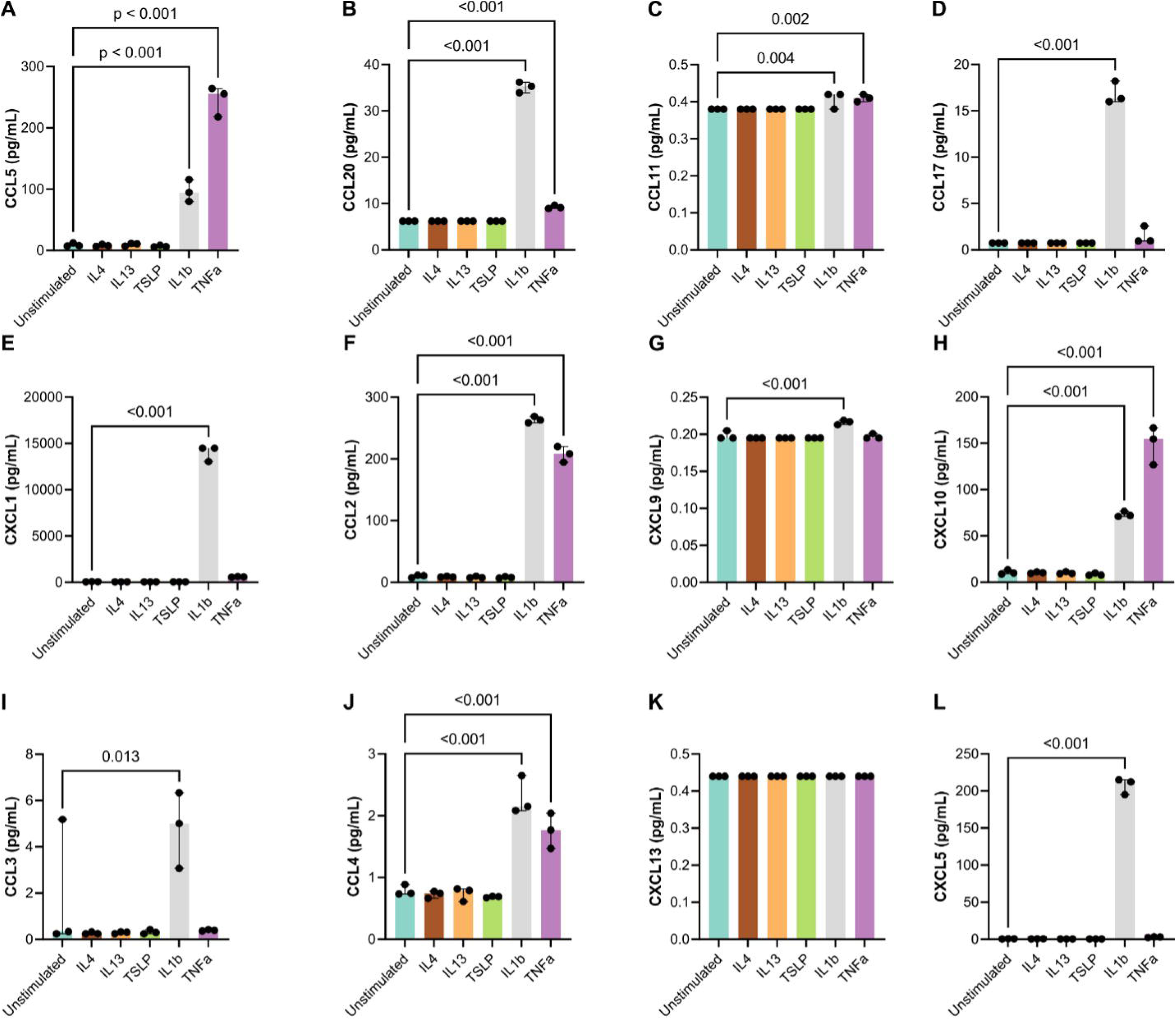
Neither IL-4, IL-13 nor TSLP stimulate secretion of 12 chemokines by the CT26 cell line *in vitro*. (A-L) Secretion of the chemokines noted on the Y-axes as measured by fluid phase immunoassay in conditioned media from monocultures of the CT26 cell line stimulated with recombinant murine IL-4, IL-13, TSLP, IL-1β or TNFα. One-way ANOVAs (CCL5 p<0.001, CCL20 p<0.001, CCL11 p=0.002, CCL17 p<0.001, CXCL1 p<0.001, CCL2 p<0.001, CXCL9 p<0.001, CXCL10 p<0.001, CCL3 p=0.003, CCL4 p<0.001, CXCL13 not tested, CXCL5 p<0.001) were followed by uncorrected Fisher’s LSD tests for differences between groups (p-values shown in figure). Each dot represents a biological replicate run on a different day at a different passage number. Bars indicate the medians and whiskers the 95% confidence intervals of each group.

**Supplemental Figure 10.**
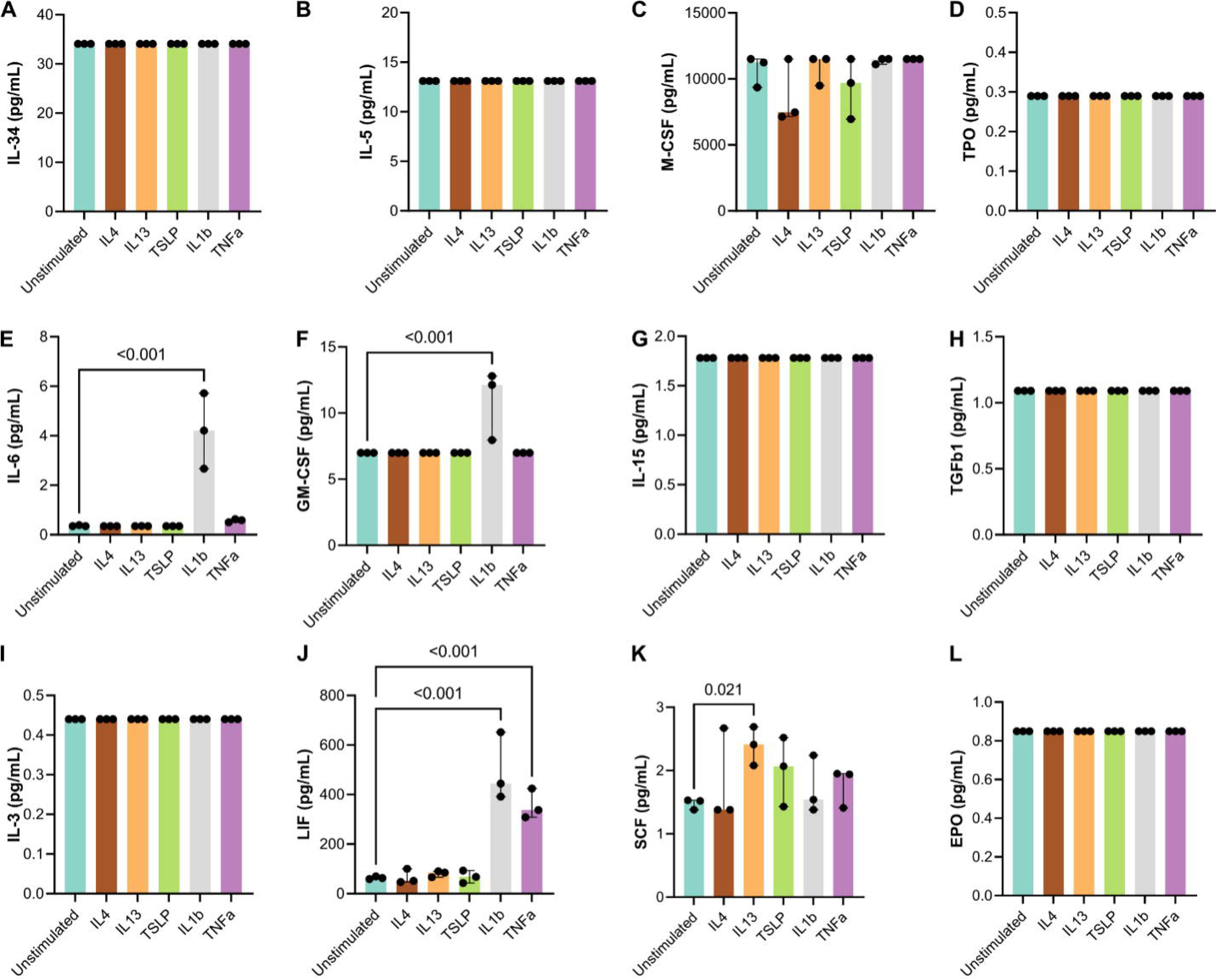
Neither IL-4, IL-13 nor TSLP stimulate secretion of 12 chemokines by the CT26 cell line *in vitro*. (A-L) Secretion of the cytokines noted on the Y-axes as measured by fluid phase immunoassay in conditioned media from monocultures of the CT26 cell line stimulated with recombinant murine IL-4, IL-13, TSLP, IL-1β or TNFα. One-way ANOVAs (IL-34 not tested, IL-5 not tested, M-CSF p=0.247, TPO not tested, IL-6 p<0.001, GM-CSF p=0.001, IL-15 not tested, TGFβ1 not tested, IL-3 not tested, LIF p<0.001, SCF p=0.319, EPO not tested) were followed by uncorrected Fisher’s LSD tests for differences between groups (p-values shown in figure). Each dot represents a biological replicate run on a different day at a different passage number. Bars indicate the medians and whiskers the 95% confidence intervals of each group.

**Supplemental Figure 11.**
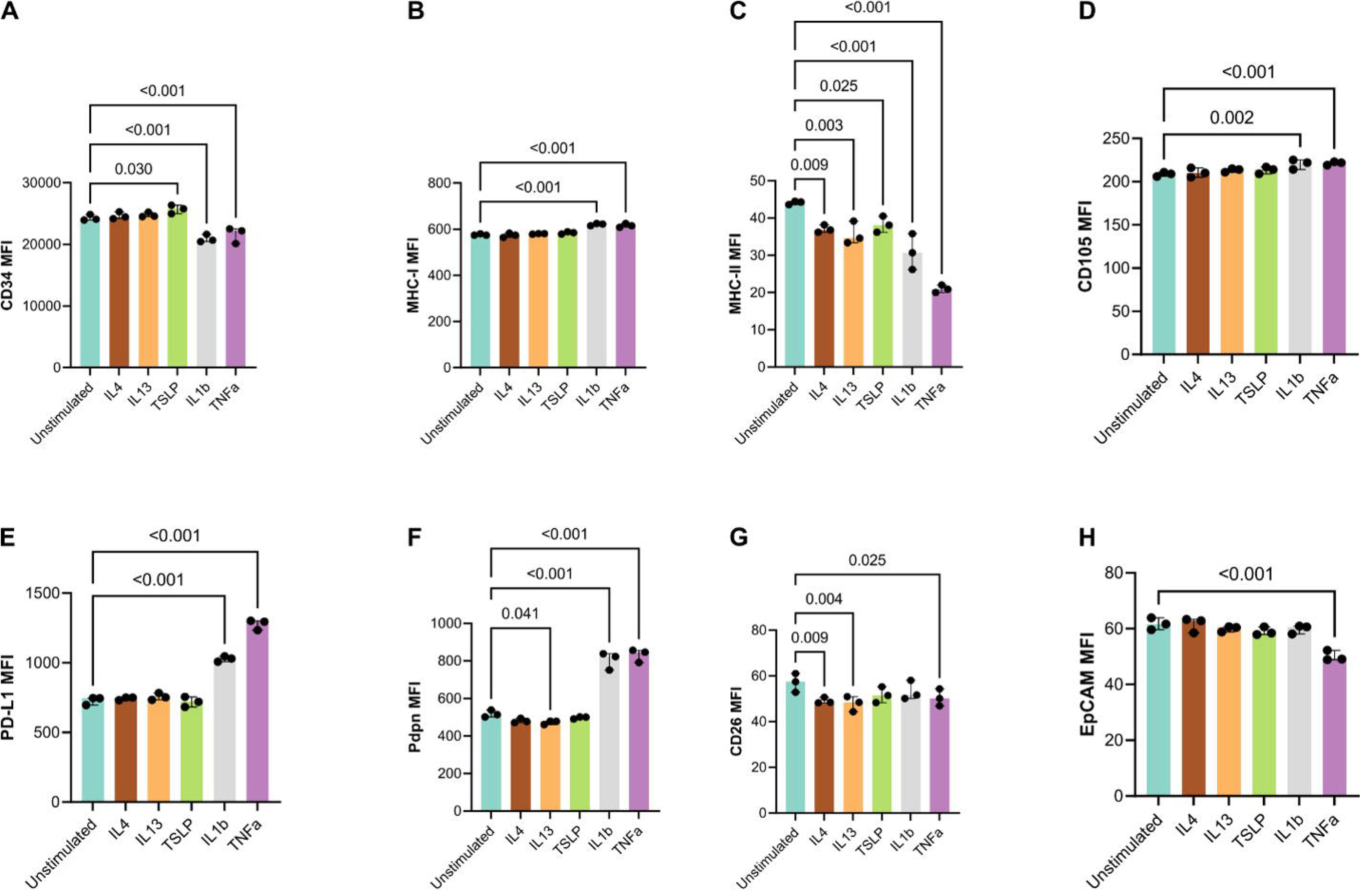
IL-4 and IL-13 slightly decrease MHC-II and CD26 expression by the CT26 cell line. (A-H) Surface expression of the proteins noted on the Y-axes by CT26 cell line monocultures stimulated with the cytokines on the X-axes as measured by flow cytometry median fluorescence intensities (MFI). One-way ANOVAs (CD34 p<0.001, MHC-I p<0.001, MHC-II p<0.001, CD105 p=0.007, PD-L1 p<0.001, Pdpn p<0.001, CD26 p=0.029, EpCAM p<0.001) followed by Fisher’s uncorrected LSD were used to test for differences between treatments. Each dot represents a biological replicate run on a different day at a different passage number. Bars indicate the medians and whiskers the 95% confidence intervals of each group.

**Supplemental Figure 12.**
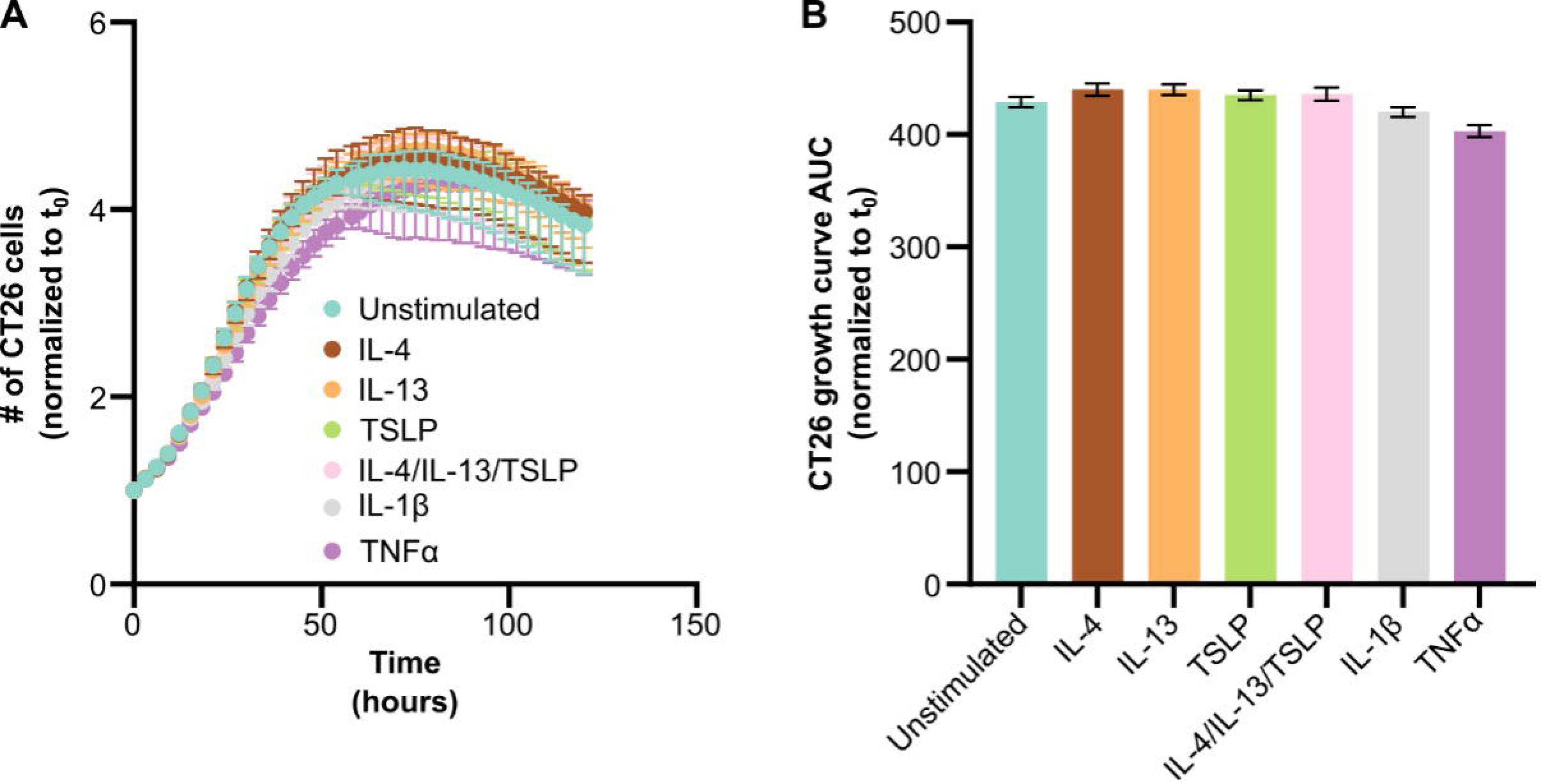
Neither IL-4, IL-13 nor TSLP alter growth of the CT26 cell line *in vitro*. **(A)** Growth curves of CT26 cell line monocultures treated with recombinant murine IL-4, IL-13, TSLP, the combination of IL-4 plus IL-13 plus TSLP, IL-1β or TNFα normalized to the cell count at initiation of the experiment (t_0_). **(B)** AUC of the growth curves from (A). Eight biological replicates were performed on different days using CT26 cells at different passages. Bars indicated the mean and 95% confidence intervals.

**Supplemental Figure 13.**
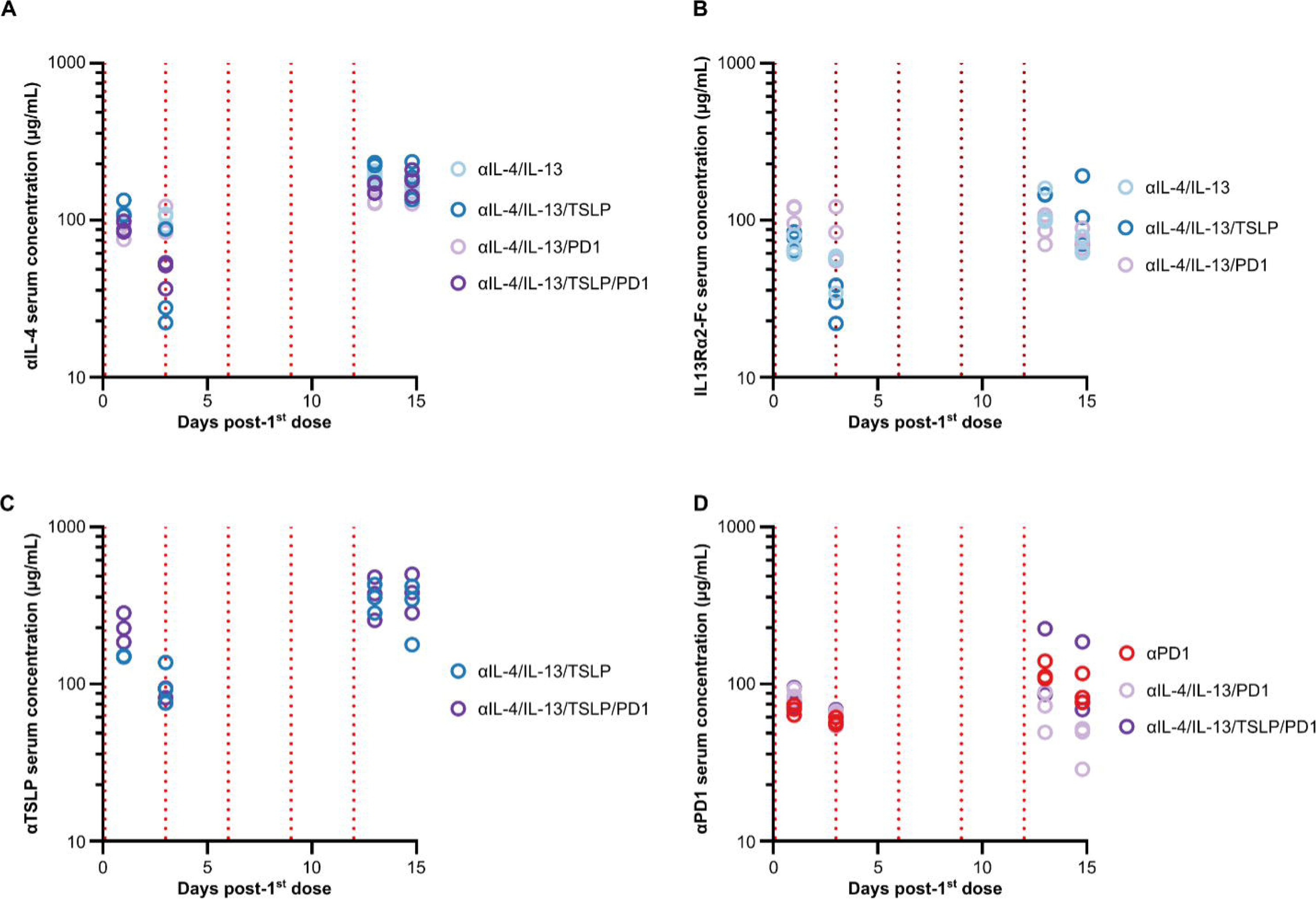
Levels of large molecule antagonists in the serum of CT26-bearing mice after the first and last doses. **(A)** Serum concentrations of the αIL-4 antagonist antibody at one and three days post-first dose and one and three days post-last dose. **(B)** Serum concentrations of the IL13Rα2-Fc antagonist at one and three days post-first dose and one and three days post-last dose. **(C)** Serum concentrations of the αTSLP antagonist antibody at one and three days post-first dose and one and three days post-last dose. **(D)** Serum concentrations of the αPD1 antagonist antibody at one and three days post-first dose and one and three days post-last dose.

**Supplemental Figure 14.**
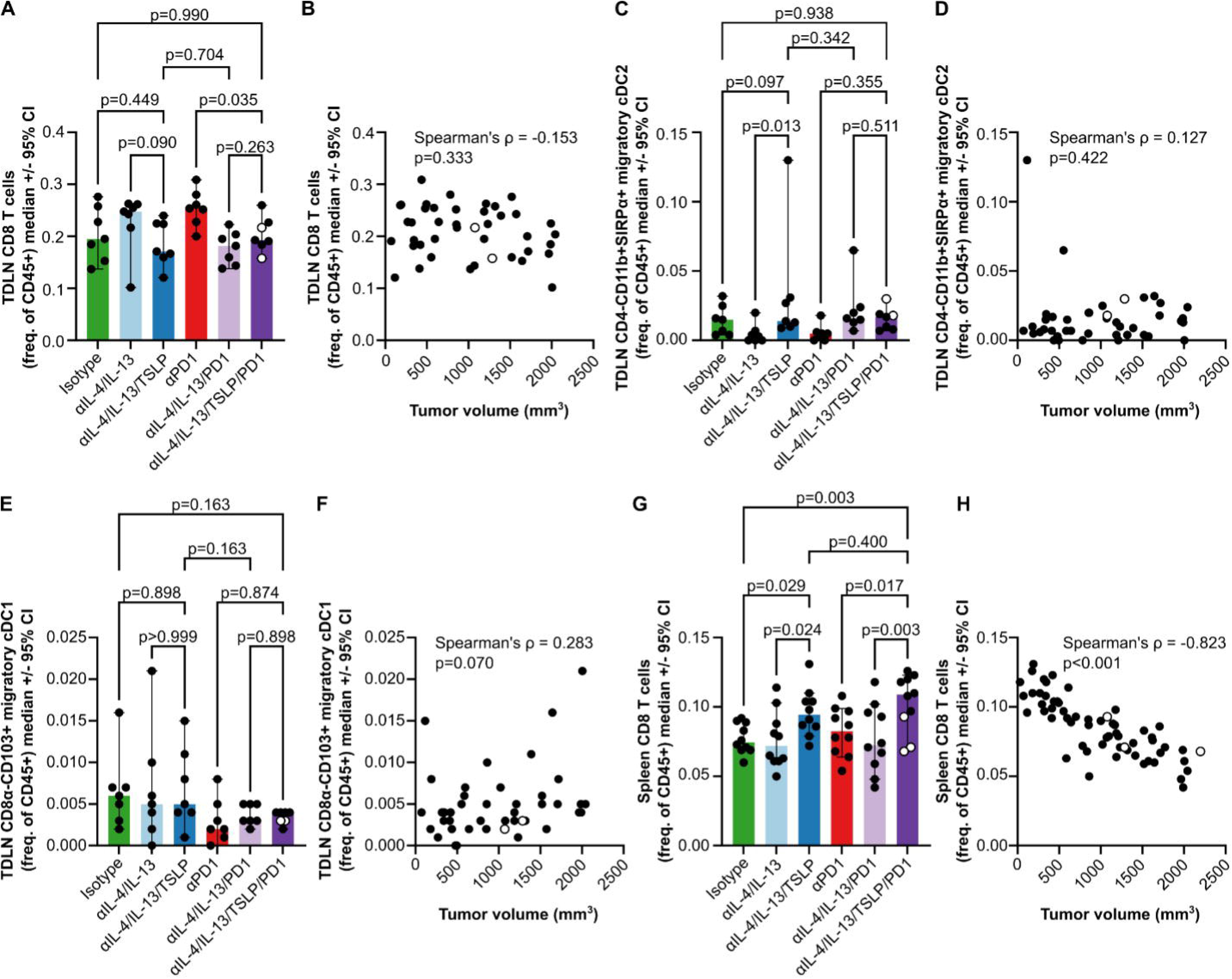
Therapeutic tumor growth inhibition is not associated with changes in CD8 T cells or migratory DC in tumor-draining lymph nodes (TDLN) but is associated with higher frequencies of CD8 T cells in the spleen. **(A)** Frequencies of CD8 T cells (in live CD45+ cells) in TDLN as measured by flow cytometry. **(B)** Correlation of TDLN CD8 T cell frequencies with tumor volume. **(C)** Frequencies of migratory cDC2 (in live CD45+ cells) in TDLN as measured by flow cytometry. **(D)** Correlation of TDLN migratory cDC2 with tumor volume. (E) Frequencies of migratory cDC1 (in live CD45+ cells) in TDLN as measured by flow cytometry. **(F)** Correlation of TDLN migratory cDC1 with tumor volumes. **(G)** Frequencies of splenic CD8 T cells (in live CD45+ cells) as measured by flow cytometry. (H) Correlation of splenic CD8 T cell frequencies with tumor volume. Differences between treatment groups in (A, C, E and G) were tested with one-way ANOVAs (TDLN CD8 T cells p=0.025, TDLN cDC2 p=0.108, TDLN cDC1 p=0.236, splenic CD8 T cells p=0.006) followed by uncorrected Fisher’s LSD tests (p-values shown in figure).

**Supplemental Figure 15.**
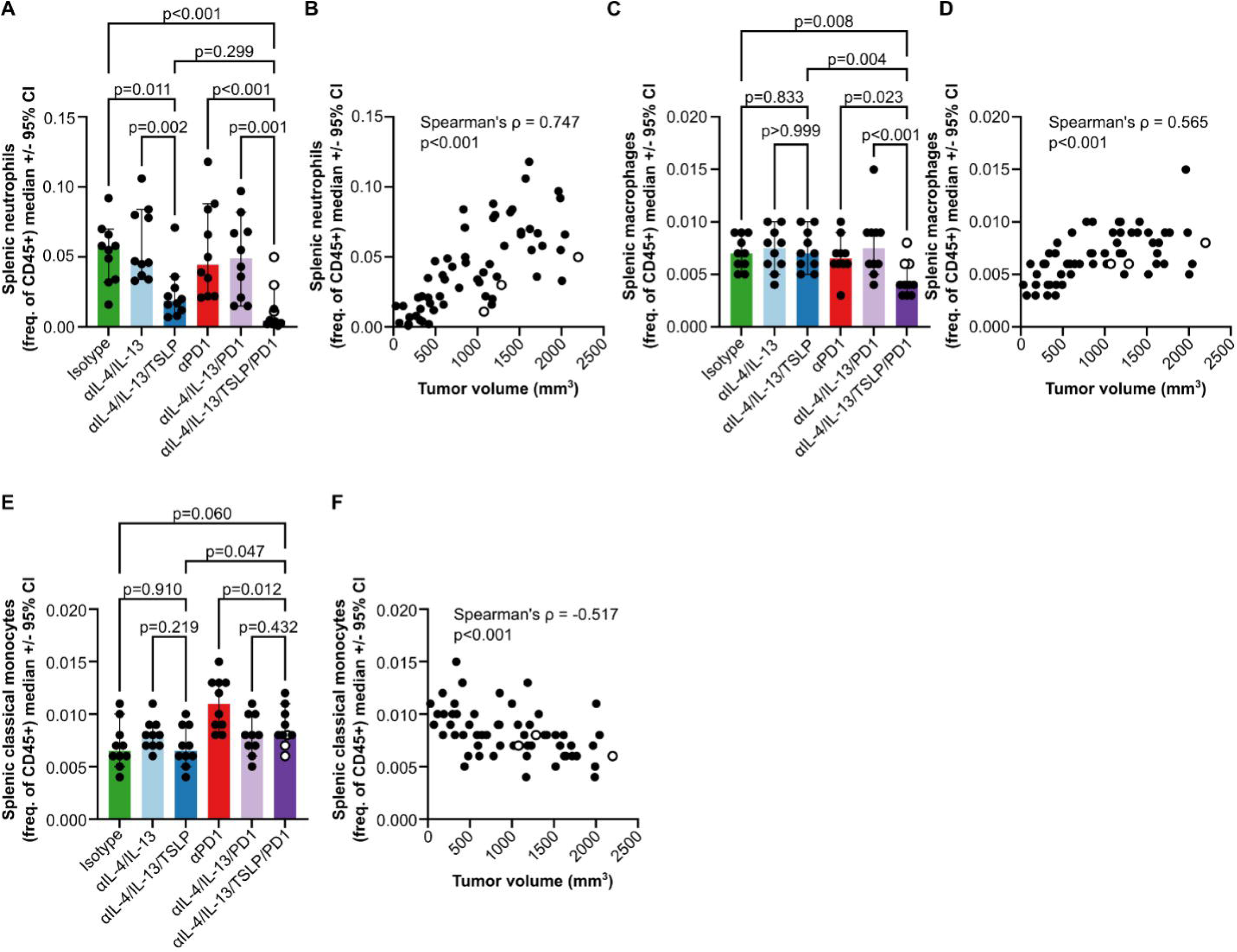
Therapeutic tumor growth inhibition was associated with reduces frequencies of splenic neutrophils and macrophages, and elevated classical monocytes. **(A)** Frequencies of splenic neutrophils (of live CD45+ cells) as measured by flow cytometry. **(B)** Correlation of splenic neutrophil frequencies with terminal tumor volumes. **(C)** Frequencies of splenic macrophages (of live CD45+ cells) as measured by flow cytometry. **(D)** Correlation of splenic macrophage frequencies with terminal tumor volumes. **(E)** Frequencies of splenic classical monocytes (of live CD45+ cells) as measured by flow cytometry. **(F)** Correlation of splenic classical monocyte frequencies with terminal tumor volumes. All bars and whiskers represent the median and 95% confidence intervals of each group, and each dot represents an individual mouse. One-way ANOVAs (neutrophils p<0.001, macrophages p=0.015, classical monocytes p<0.001) followed by uncorrected Fisher’s LSD tests (p-values shown in figure) were used to test for differences between treatment groups. Spearman’s test was used for correlations, and ρ and p-values are shown in the figure.

**Supplemental Figure 16.**
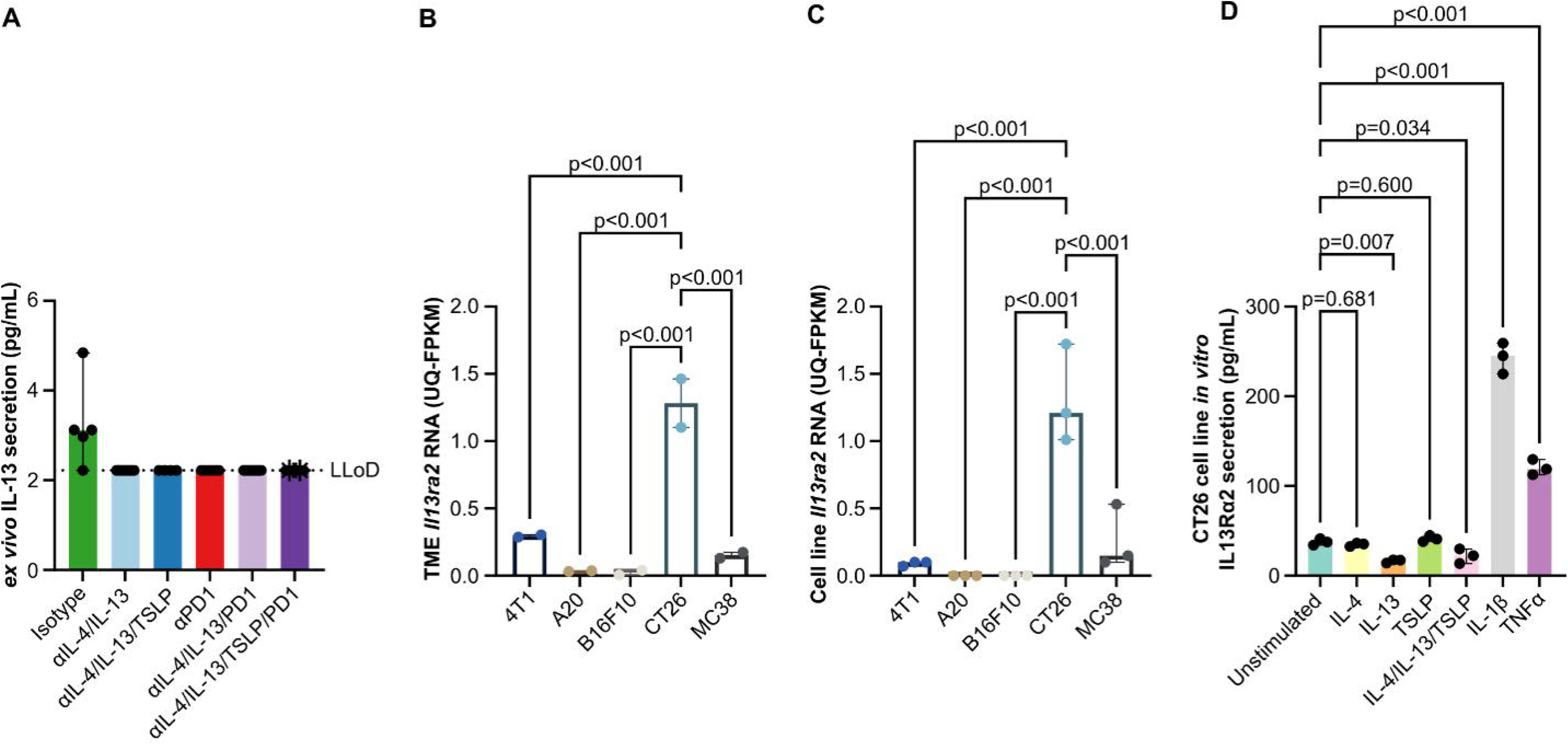
Production of IL13Rα2 likely prohibits detection of IL-13 secretion by restimulated subcutaneous CT26 TME cells *ex vivo*. **(A)** Secreted IL-13 protein detected in supernatants of total cells isolated from subcutaneous CT26 tumors restimulated with PMA and ionomycin *ex vivo*. **(B)** RNA expression of *Il13ra2* in subcutaneous CT26 tumors measured by bulk RNAseq. **(C)** RNA expression of *Il13ra2* by CT26 cell line monocultures *in vitro* measured by bulk RNAseq. **(D)** Secretion of IL-13Rα2 protein by CT26 cell line monocultures *in vitro* stimulated with the recombinant murine cytokines noted on the X-axis measured by enzyme linked immunosorbent assay (ELISA). All bars and whiskers represent the median and 95% confidence intervals of each group, and each dot is an individual mouse or biological replicate. One-way ANOVAs (*ex vivo* IL-13 not tested, TME *Il13ra2* p<0.001, cell line *Il13ra2* p<0.001, CT26 cell line IL-13Rα2 p<0.001) followed by uncorrected Fisher’s LSD tests (p-values shown in figure) were used to test for differences between groups.

**Supplemental Figure 17.**
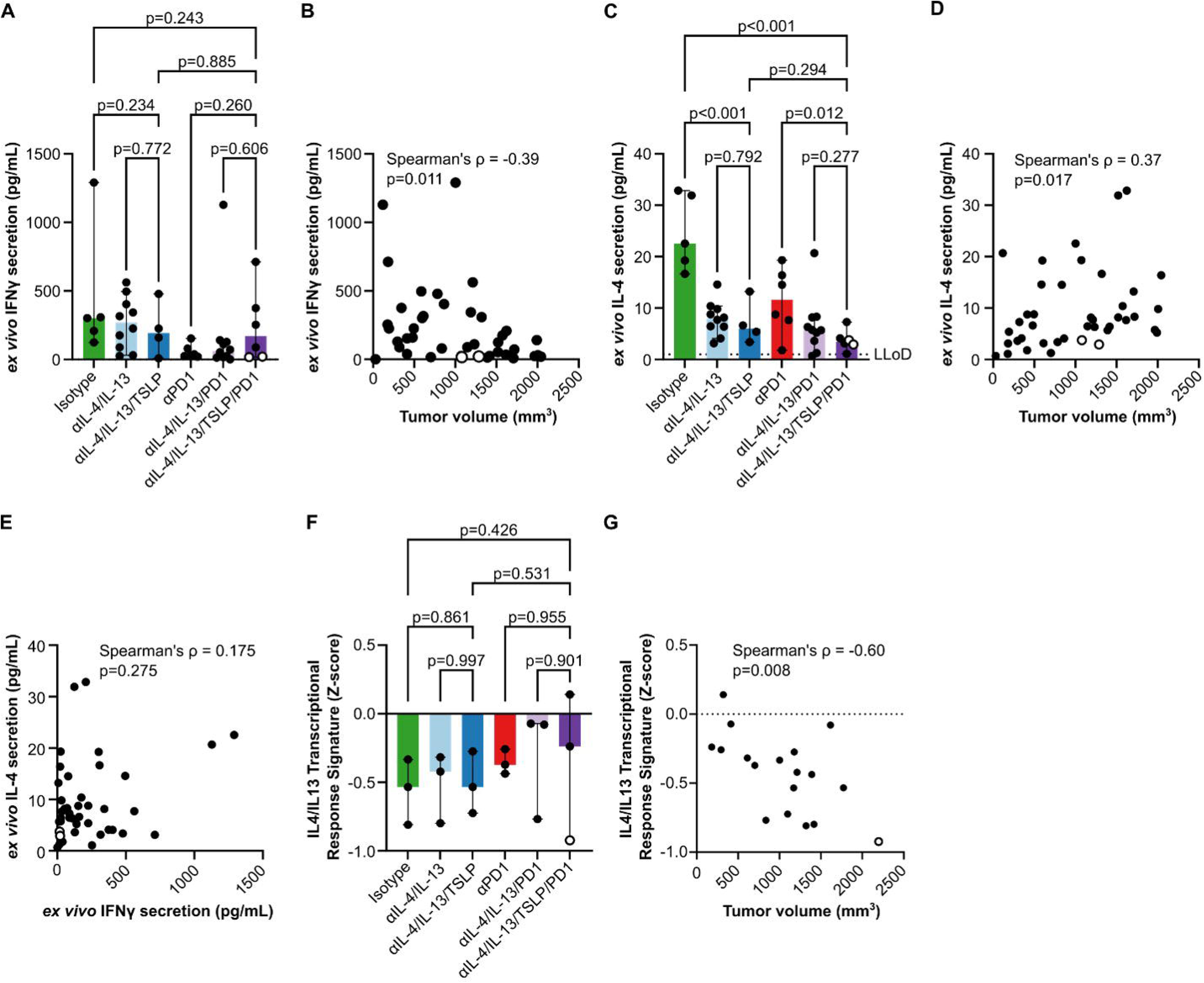
Therapeutic tumor growth inhibition was associated with reduced type 2 polarization rather than increased type 1 polarization. **(A)** Secreted IFNγ protein detected in supernatants of total cells isolated from subcutaneous CT26 tumors restimulated with PMA and ionomycin *ex vivo*. (B) Correlation of IFNγ from (A) with tumor volumes. **(C)** Secreted IL-4 protein detected in supernatants of total cells isolated from subcutaneous CT26 tumors restimulated with PMA and ionomycin *ex vivo*. **(D)** Correlation of IL-4 from (C) with tumor volumes. **(E)** Correlation of IL-4 from (C) with IFNγ from (A). (F) IL-4/IL-13 transcriptional signature levels in a subset of subcutaneous CT26 tumors measured by bulk RNAseq. **(G)** Correlation of IL-4/IL-13 transcriptional signature levels from (F) with tumor volumes. All bars and whiskers represent the median and 95% confidence intervals of each group, and each dot represents an individual mouse. One-way ANOVAs (IFNγ p=0.343, IL-4 p<0.001, IL-4/IL-13 transcriptional signature p=0.882) followed by uncorrected Fisher’s LSD tests (p-values shown in figure) tested for differences between groups. Spearman’s tests were used for correlations and ρ and p-values are shown in the figure.

**Supplemental Figure 18.**
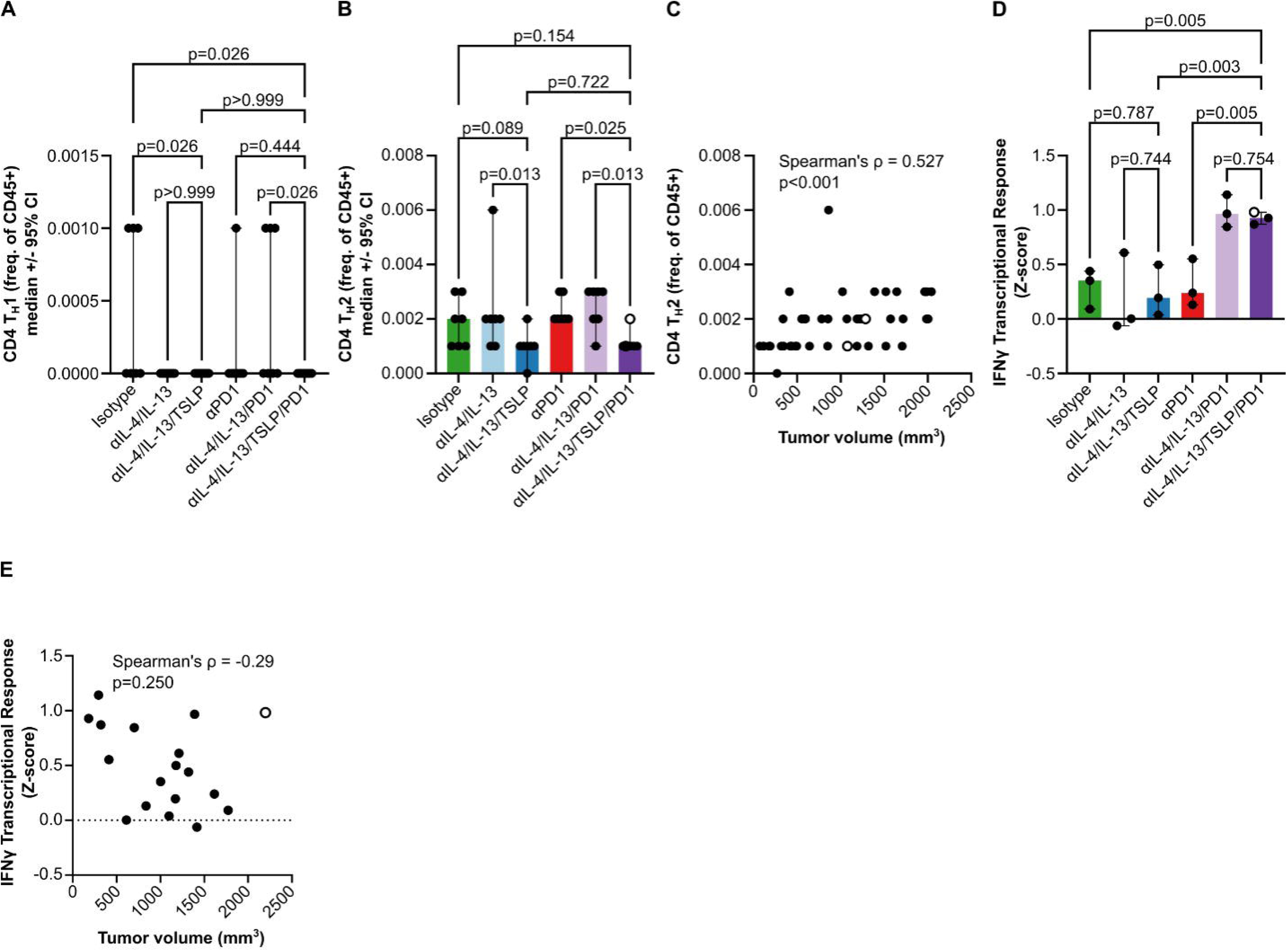
Therapeutic tumor growth inhibition was associated with lower T_H_2 frequencies and higher IFNγ transcriptional response signature expression in TDLN. **(A)** Frequencies of T_H_1 (of live CD45+ cells) in TDLN as measured by flow cytometry. **(B)** Frequencies of T_H_2 (of live CD45+ cells) in TDLN as measured by flow cytometry. **(C)** Correlation of TDLN frequencies of T_H_2 from (B) with tumor volumes. **(D)** IFNγ transcriptional response signature levels measured by bulk RNAseq in a subset of TDLN. **(E)** Correlation of TDLN IFNγ transcriptional response signature levels from (D) with tumor volumes. All bars and whiskers represent the median and 95% confidence intervals of groups, and dots represent individual mice. One-way ANOVAs (T_H_1 p=0.042, T_H_2 p=0.017, IFNγ transcriptional response p=0.002) followed by uncorrected Fisher’s LSD tests (p-values shown in figure) tested for differences between groups. Spearman’s test for correlations was used and ρ and p-values are shown in the figure.

**Supplemental Figure 19.**
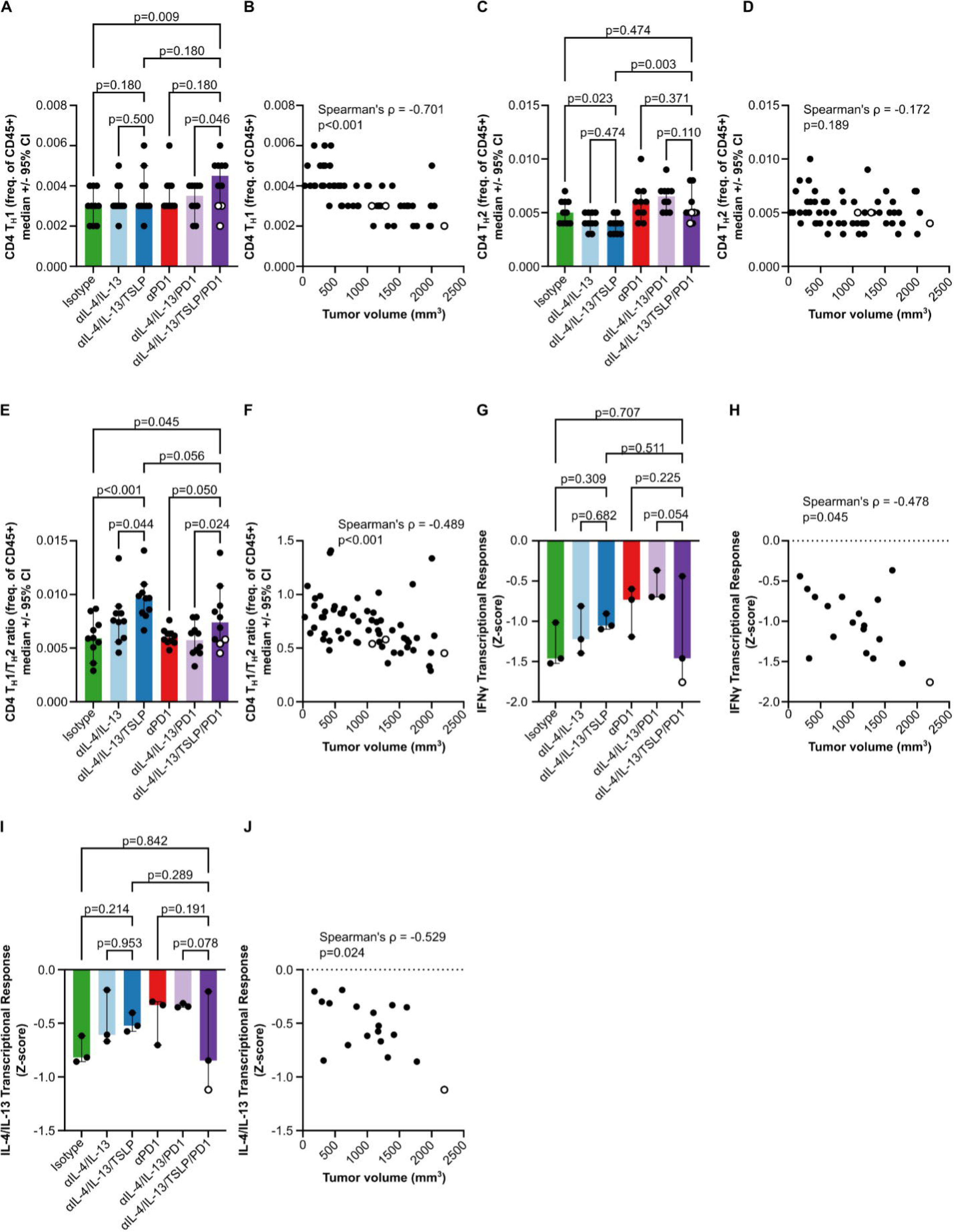
Therapeutic tumor growth inhibition was associated with splenic repolarization away from type 2 towards type 1 immunity. **(A)** Frequencies of splenic T_H_1 (of live CD45+ cells) as measured by flow cytometry. **(B)** Correlation of splenic TH1 frequencies from (A) with tumor volumes. (C) Frequencies of splenic T_H_2 (of live CD45+ cells) as measured by flow cytometry. **(D)** Correlation of splenic T_H_2 frequencies from (C) with tumor volumes. (E) Ratio of splenic T_H_1/T_H_2 frequencies from (A) and (C). (F) Correlation of splenic TH1/TH2 frequency ratios from **(E)** with tumor volumes. **(G)** Splenic IFNγ transcriptional response signature levels as measured by bulk RNAseq in a subset of spleens. (H) Correlation of splenic IFNγ transcriptional response signature levels from (G) with tumor volumes. **(I)** Splenic IL-4/IL-13 transcriptional response signature levels as measured by bulk RNAseq in a subset of spleens. **(J)** Correlation of splenic IL-4/IL-13 transcriptional response signature levels from (I) with tumor volumes. All bars and whiskers represent the median and 95% confidence intervals of each group, and dots represent individual mice.

**Supplemental Figure 20.**
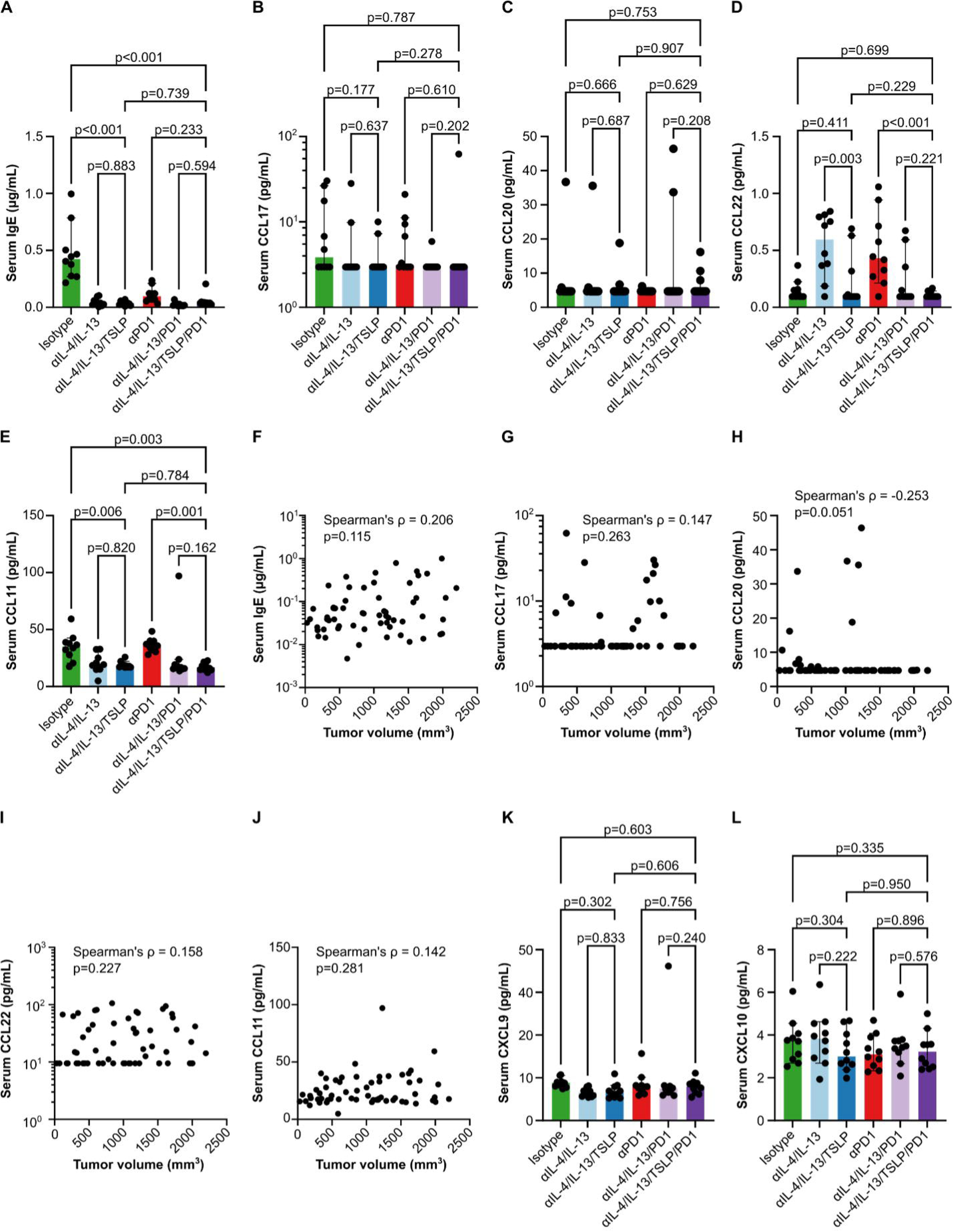
Serum CCL11 is a pharmacodynamic biomarker of IL-4 and IL-13 blockade in the subcutaneous CT26 tumor model. (A-E) Serum levels of IgE (A), CCL17 (B), CCL20 (C), CCL22 (D) or CCL11 (E) per treatment group as measured by fluid phase immunoassay. **(F-J)** Correlations of serum levels of IgE (F), CCL17 (G), CCL20 (H), CCL22 (I) or CCL11 (J) with terminal tumor volumes. **(K-L)** Serum levels of CXCL9 (K) and CXCL10 (L) per treatment group as measured by fluid phase immunoassay. All bars and whiskers represent the median and 95% confidence intervals per group, and each dot represents an individual mouse. One-way ANOVAs (IgE p<0.001, CCL17 p=0.603, CCL20 p=0.615, CCL22 p<0.001, CCL11 p=0.002) followed by uncorrected Fisher’s LSD (p-values shown in figure) were used to test for differences between groups. Spearman’s tests for correlation were used with ρ and p-values shown in the figure.

**Supplemental Figure 21.**
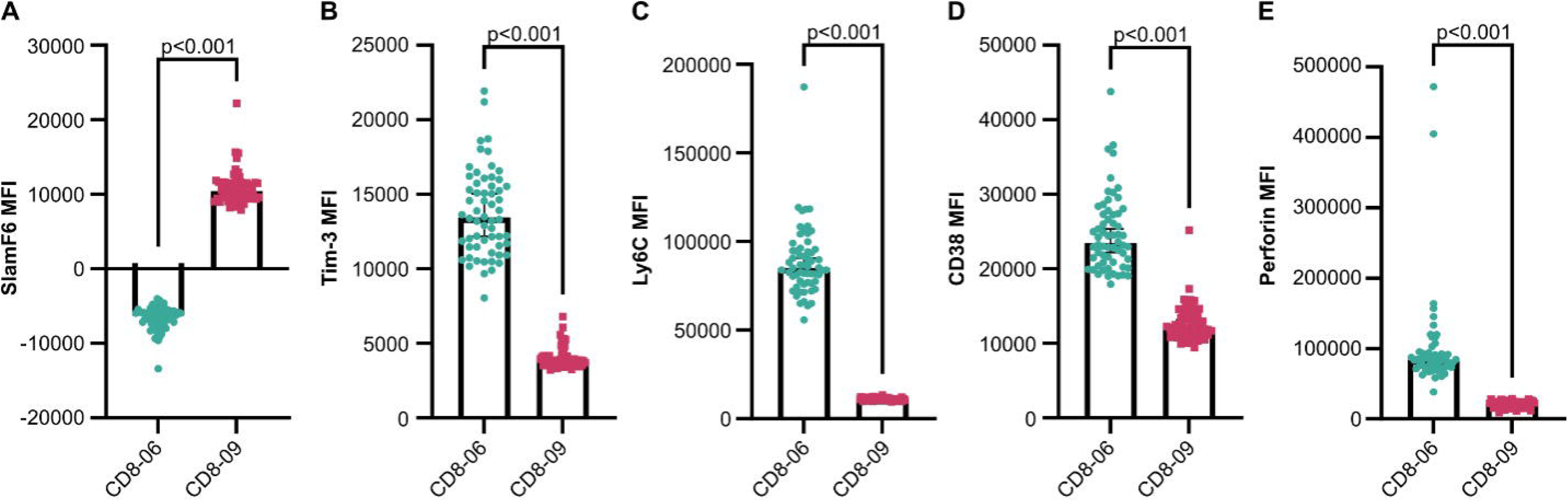
Expression of SlamF6, Tim-3, Ly6C, CD38 and Perforin indicate that tumor subset CD8-06 is effector-like and CD8-09 is stem memory-like. (A-E) Surface expression of SlamF6, Tim-3, Ly6C, CD38 and cytoplasmic expression of Perforin by the tumor-infiltrating CD8 T cell subsets CD8-06 and CD8-09 as measured by flow cytometric median fluorescence intensity (MFI). Each dot represents an individual mouse (mice from all treatment groups were pooled for this analysis), the bars represent the median and whiskers represent the 95% confidence intervals for each CD8 T cell subset. Paired t-tests were used to identify differences between subsets and p-values are shown in the figure.

**Supplemental Figure 22.**
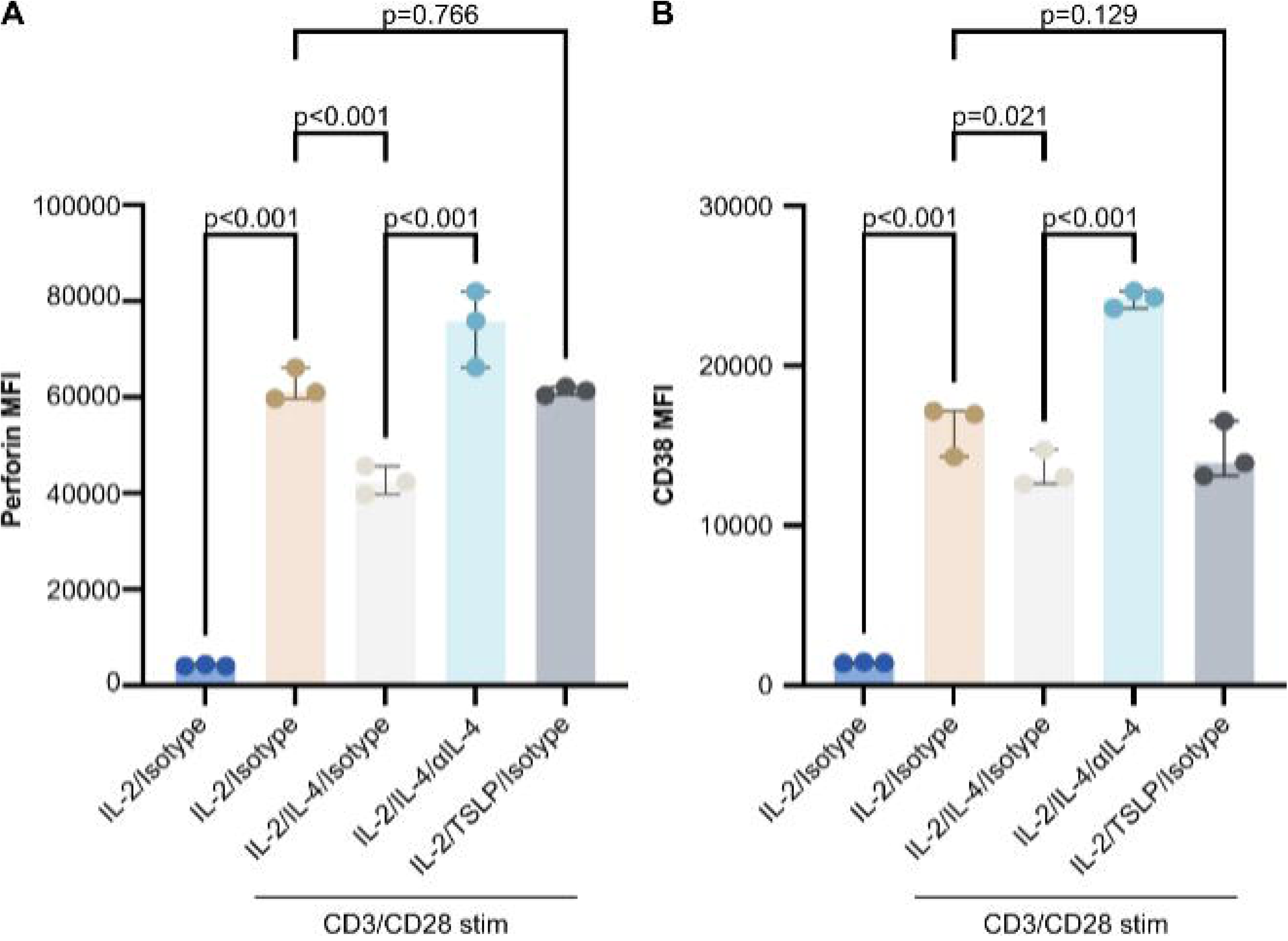
IL-4 impairs activation-induced expression of Perforin and CD38 by primary human CD8 T cells *in vitro*. (A-B) Levels of cytoplasmic Perforin (A) or surface CD38 (B) expression by primary human CD8 T cells after stimulation with agonist αCD3/CD28 antibodies in the presence of the recombinant human cytokines and antagonist antibodies noted on the X-axes. Protein levels were measured by flow cytometry and are presented as median fluorescence intensities (MFI). Bars and whiskers represent the median and 95% confidence intervals of each group. Each dot represents a biological replicate (experiments run on different days) from a single donor. One-way ANOVAs (Perforin p<0.001 and CD38 p<0.001) followed by uncorrected Fisher’s LSD tests (p-values shown in figure) were used.

**Supplemental Figure 23.**
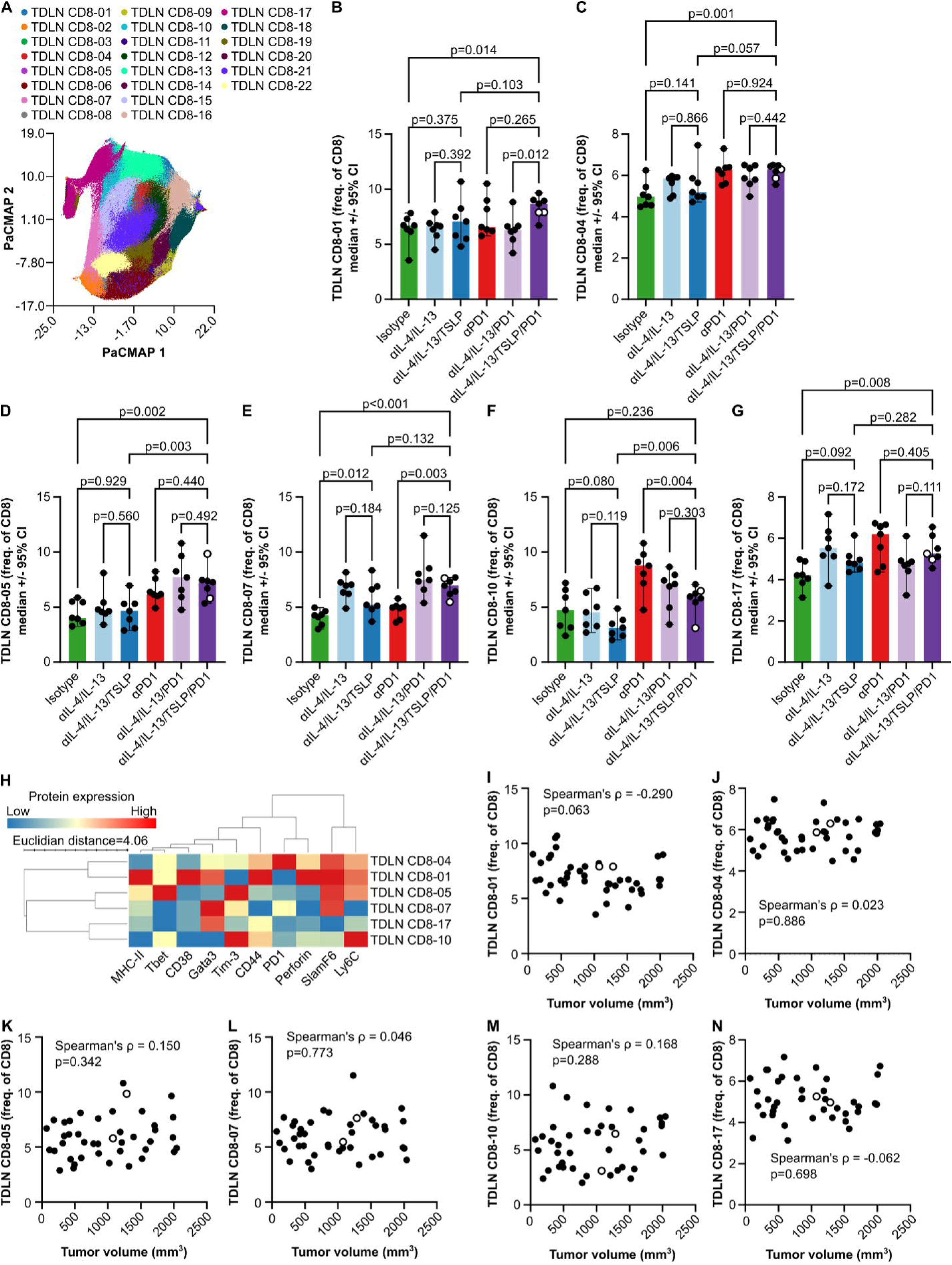
Combined blockade of IL-4, IL-13, TSLP and PD1 altered six CD8 T cell subset frequencies in the TDLN, but none were associated with smaller tumor volumes. **(A)** PaCMAP dimensionality reduction and PARC clustering of TDLN CD8 T cells using CD38, Tbet, MHC-II, Ly6C, Tim-3, CD44, SlamF6, Gata3 and Perforin protein expression. **(B-G)** Frequencies of selected TDLN CD8 T cell subsets from (A) out of total TDLN CD8 T cells. Bars and whiskers represent the median and 95% confidence intervals of each group, and each dot represents a single mouse. One-way ANOVAs (TDLN CD8-01 p=0.073, TDLN CD8-04 p=0.008, TDLN CD8-05 p<0.001, TDLN CD8-07 p<0.001, TDLN CD8-10 p<0.001, TDLN CD8-17 p=0.008) followed by uncorrected Fisher’s LSD tests (p-values shown in figure) were used. **(H)** Heatmap showing the scaled expression of each protein used in the dimensionality reduction and clustering in (A) across selected TDLN CD8 T cell subsets. **(I-M)** Correlations of TDLN CD8 T cell subset frequencies from (B-G) with terminal tumor volumes. Spearman’s correlations were used and ρ and p-values are shown in the figure.

**Supplemental Figure 24.**
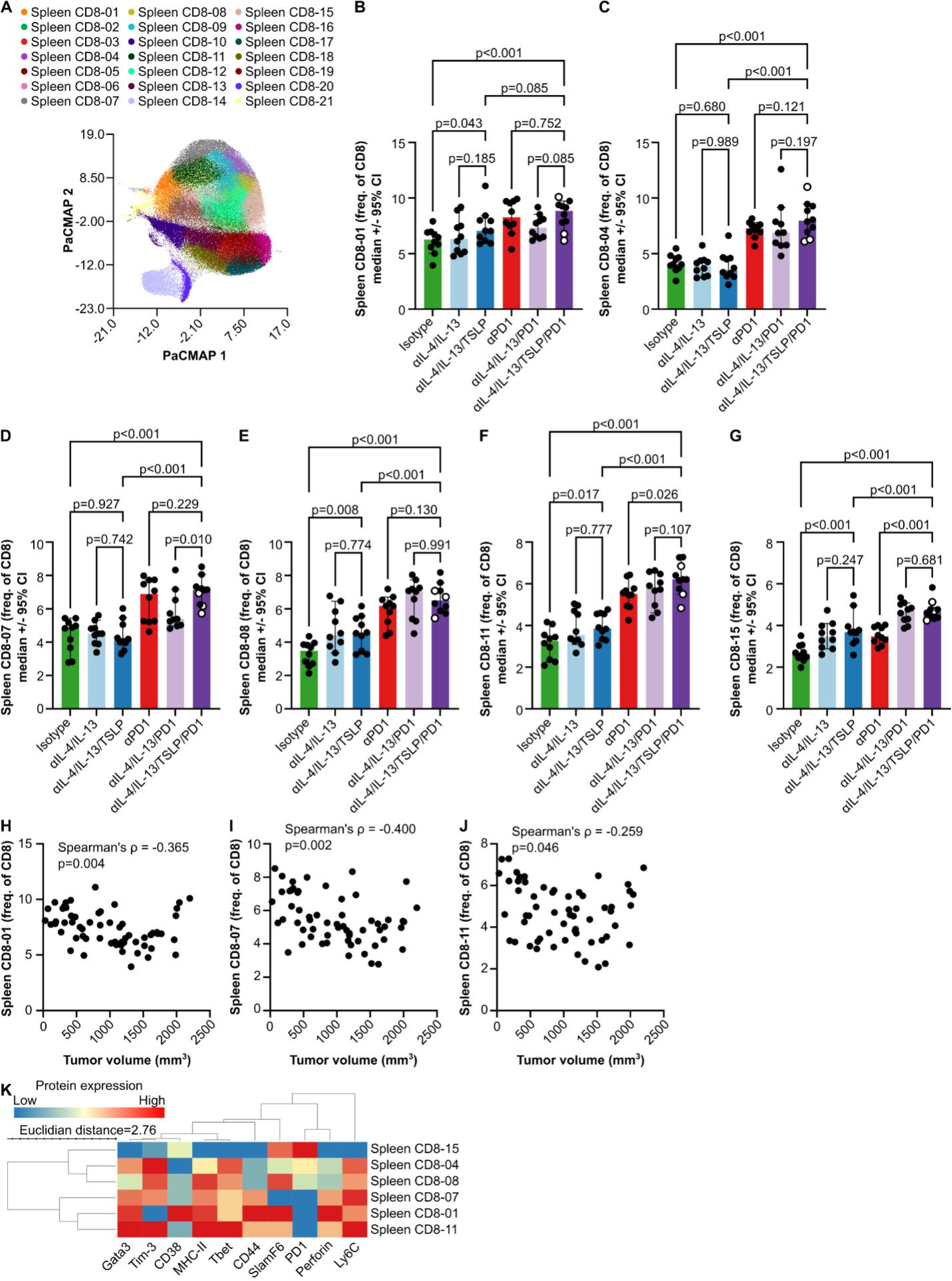
Therapeutic tumor growth inhibition was associated with increases in three splenic CD8 T cell stem- and effector memory-like subsets. **(A)** PaCMAP dimensionality reduction and PARC clustering of splenic CD8 T cells using CD38, Tbet, MHC-II, Ly6C, Tim-3, CD44, SlamF6, Gata3 and Perforin protein expression as measured by flow cytometry. **(B-G)** Frequencies of selected splenic CD8 T cell subsets from (A) out of total splenic CD8 T cells. Bars and whiskers represent the median and 95% confidence intervals of each group, and each dot represents a single mouse. One-way ANOVAs (Spleen CD8-01 p=0.002, Spleen CD8-04 p<0.001, Spleen CD8-07 p<0.001, Spleen CD8-08 p<0.001, Spleen CD8-11 p<0.001, Spleen CD8-15 p<0.001) followed by uncorrected Fisher’s LSD tests (p-values shown in figure) were used. (**H-J)** Correlations of selected splenic CD8 T cell subset frequencies from (B, D, F) with terminal tumor volumes. Spearman’s correlations were used and ρ and p-values are shown in the figure. **(K)** Heatmap showing the scaled expression of each protein used in the dimensionality reduction and clustering in (A) across selected splenic CD8 T cell subsets.

**Supplemental Figure 25.**
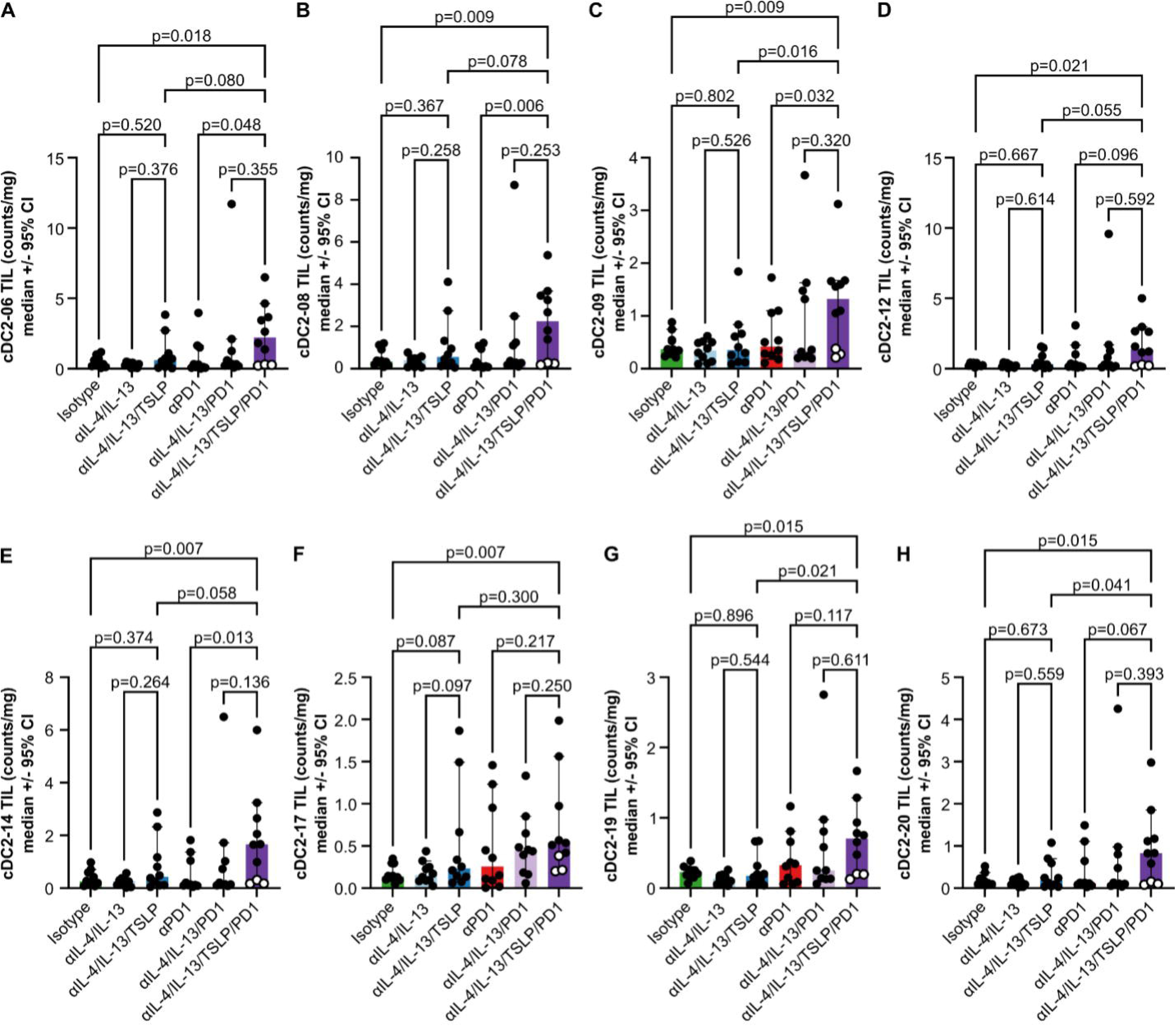
Quadruple blockade of IL-4, IL-13, TSLP and PD1 increased eight tumor-infiltrating cDC2 subsets. (A-H) Mass-normalized counts of tumor-infiltrating PARC cDC2 subsets (from Fig. 7A) as measured by flow cytometry. Bars and whiskers represent the median and 95% confidence intervals of each group, and each dot represents a single mouse. One-way ANOVAs (cDC2-06 p=0.090, cDC2-08 p=0.029, cDC2-09 p=0.025, cDC2-12 p=0.086, cDC2-14 p=0.038, cDC2-17 p=0.055, cDC2-19 p=0.024, cDC2-20 p=0.072) followed by uncorrected Fisher’s LSD tests (p-values shown in figure) were used.

**Supplemental Figure 26.**
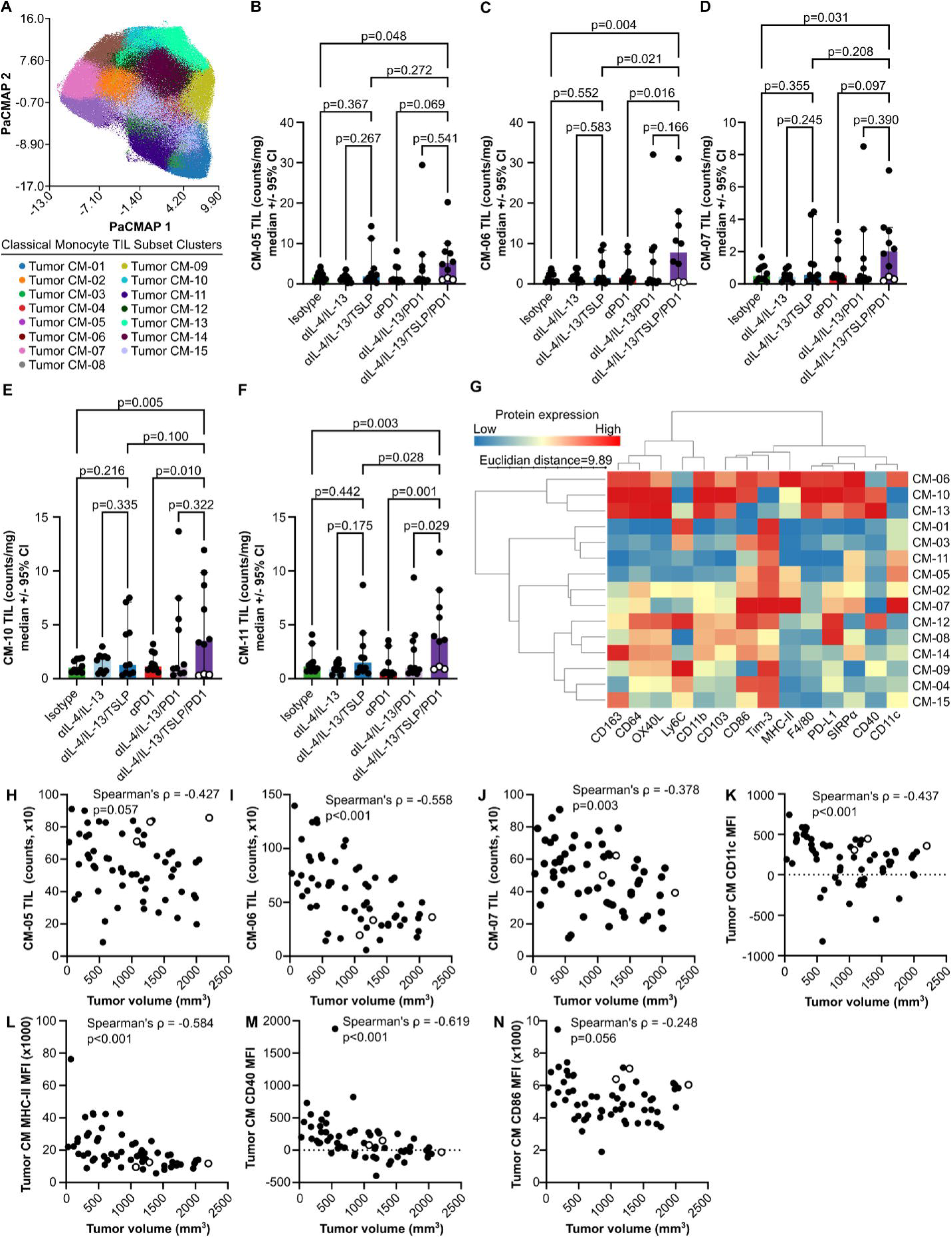
Therapeutic tumor growth inhibition is correlated with increases in three tumor-infiltrating classical monocyte populations that express CD11c, MHC-II and CD86. **(A)** PaCMAP dimensionality reduction and PARC clustering of tumor-infiltrating classical monocytes using CD163, CD64, OX40L, Ly6C, CD11b, CD103, CD86, Tim-3, MHC-II, F4/80, PD-L1, SIRPα, CD40 and CD11c protein expression as measured by flow cytometry. **(B-F)** Mass-normalized counts of tumor-infiltrating PARC classical monocyte subsets from (A) as measured by flow cytometry. Bars and whiskers represent the median and 95% confidence intervals of each group, and each dot represents a single mouse. One-way ANOVAs (CM-05 p=0.188, CM-07 p=0.166, CM-10 p=0.032, CM-11 p=0.007, CM-13 p=0.412) followed by uncorrected Fisher’s LSD tests (p-values shown in figure) were used. (**G)** Heatmap showing the scaled expression of each protein used in the dimensionality reduction and clustering in (A) by all tumor-infiltrating classical monocyte subsets. **(H-J)** Correlations of selected tumor-infiltrating classical monocyte cluster counts with terminal tumor volumes**. (K-N)** Correlations of protein expression by classical monocytes as measured by flow cytometry with terminal tumor volumes. Spearman’s correlations were used and ρ and p-values are shown in the figures.

**Supplemental Figure 27.**
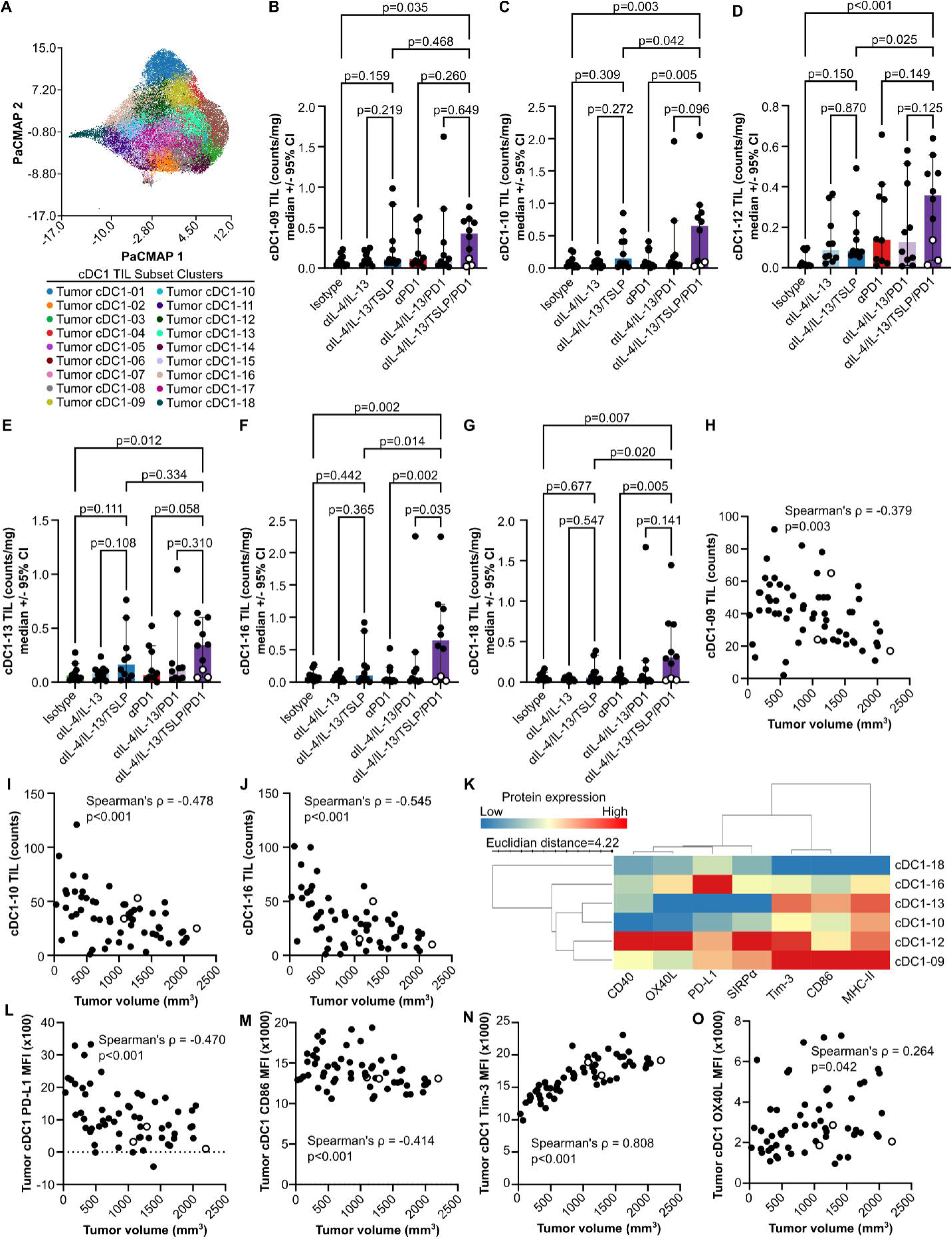
Therapeutic tumor growth inhibition is correlated with increases in three tumor-infiltrating cDC1 subsets. **(A)** PaCMAP dimensionality reduction and PARC clustering of tumor infiltrating (TIL) cDC1 subsets using protein expression of SIRPα, PD-L1, Tim-3, CD86, OX40L, CD40 and MHC-II measured by flow cytometry. **(B-G)** Mass-normalized counts of tumor-infiltrating PARC cDC1 subsets from (A) as measured by flow cytometry. Bars and whiskers represent the median and 95% confidence intervals of each group, and each dot represents a single mouse. One-way ANOVAs (cDC1-09 p=0.220, cDC1-10 p=0.020, cDC1-12 p=0.014, cDC1-13 p=0.074, cDC1-16 p=0.009, cDC1-18 p=0.028) followed by uncorrected Fisher’s LSD tests (p-values shown in figure) were used. **(H-J)** Correlations of selected tumor-infiltrating cDC1 cluster counts with terminal tumor volumes. **(K)** Heatmap showing the scaled expression of each protein used in the dimensionality reduction and clustering in (A) by selected tumor infiltrating cDC1 subsets. **(L-O)** Correlations of protein expression by cDC1 as measured by flow cytometry with terminal tumor volumes. Spearman’s correlations were used and ρ and p-values are shown in the figure.

**Supplemental Figure 28.**
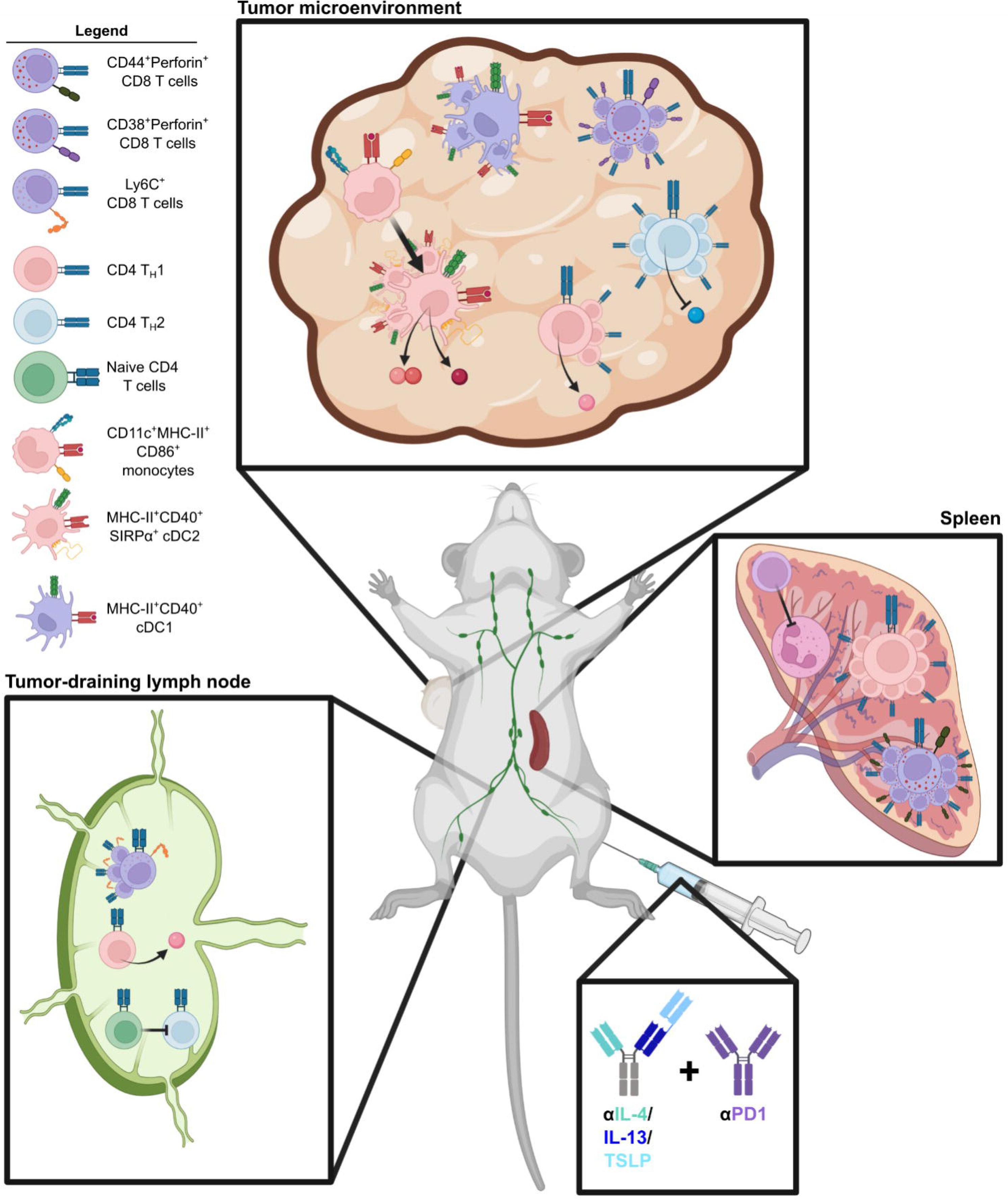
Proposed model of how blocking IL-4, IL-13 and TSLP generates additive anti-tumor immune stimulating synergy with PD1 antagonism. Therapeutic TGI by αIL-4/IL-13/TSLP/PD1 was associated with decreased T_H_2, increased transcriptional response to IFNγ and a shift towards CD8 T cells expressing Ly6C with intermediate levels of Perforin and SlamF6 in TDLN. These changes in TDLN were coincident with increased splenic T_H_1 and CD8 T cells with higher Perforin and CD44 expression, as well as decreased splenic neutrophil frequencies. αIL-4/IL-13/TSLP/PD1 TGI was also concomitant with a type 2 to type 1 TME repolarization including increased infiltration by pre-cDC2-like classical monocytes, cDC2 expressing a type 1-polarized program, increased IFNγ response and reduced IL-4 secretion. TME infiltration by effector-like CD8 T cells expressing high levels of Perforin and CD38 and an expansion of MHC-II and CD40 high cDC1 were also associated with αIL-4/IL-13/TSLP/PD1 TGI. Figure created with BioRender.com.

