## Supplemental Methods and Tables for "Therapeutic Blockade of Type 2 Cytokines and PD1 Unleashes Anti-Tumor Immunity Through Coordinated Reprogramming of Innate and Adaptive Immune Surveillance"

In vitro *T cell priming and tumor growth inhibition assay and cytokine secretion measurement*

Peripheral blood mononuclear cells (PBMC) were isolated from three healthy donors by density gradient centrifugation over Ficoll-Paque Plus (Cytiva) and frozen before use. Total CD4 and CD8 T cells were isolated by negative selection using EasySep kits (Stemcell Technologies) per the manufacturer’s instructions. Apoptosis was induced in A375 human tumor cells by treatment with mitomycin C (Stemcell Technologies). To expand A375-reactive T cells, equal numbers of apoptotic A375 and CD4 or CD8 T cells were combined in G-REX plates (Wilson-Wolfe) in X-Vivo 15 medium (Lonza) supplemented with 10% normal human AB serum (Fisher Scientific), 20ng/mL recombinant human IL-2 improved sequence (Miltenyi Biotec) and 10ng/mL recombinant human IL-7 (Miltenyi Biotec). Recombinant human IL-4 (0.01ng/mL, Biolegend) or IL-12 (2ng/mL, R&D Systems), anti-IL-4 antibody (12pM, clone IL4-1040, Pfizer, Inc.) or isotype antibody (12pM, clone mAb8.8, Pfizer, Inc.) were included as appropriate. T cells were expanded for 11-12 days with cytokines and antibodies refreshed every 2-3 days. At the end of this priming phase conditioned media was collected for cytokine measurements. Cytokines were quantified using V-plex kits (Meso Scale Diagnostics) per the manufacturer’s instructions.

Expanded CD4 and CD8 T cells were harvested from the priming cultures, mixed at a 1:1 ratio and co-cultured with A375 tumor cells expressing nuclear-restricted green fluorescent protein (GFP) at a ratio of 20 T cells:1 A375 cell. Recombinant IL-4, IL-12, isotype and anti-IL-4 antibodies were also included at the same concentrations as above. TGI was measured by longitudinal GFP imaging and counting on an Incucyte S3 microscope (Sartorius). Growth curves were normalized to the initial number of cells and area under the curves (AUC) calculated using Prism (GraphPad). Lower AUC measurements indicate fewer tumor cells and better TGI.

In vitro *human monocyte-derived DC differentiation and stimulation assay*

Frozen CD14+ monocytes from adult blood donors were purchased from Stemcell Technologies. Monocytes were differentiated in Immunocult-ACM (Stemcell Technologies) with DC Differentiation Supplement (Stemcell Technologies) per the manufacturer’s instructions. Differentiated DC were harvested, washed and replated in Immunocult-ACM supplemented with 50ng/mL recombinant human GM-CSF (R&D Systems). Immunocult DC maturation supplement (Stemcell Technologies) and recombinant human IL-4, IL-13 or TSLP (all from Biolegend, used at 0.8ng/mL) were added to the appropriate wells. Monocyte-derived DC were matured for 48 hours after which conditioned media was collected and frozen at -80°C. Secreted proteins were quantified in thawed conditioned media using the human cytokine 2 Legendplex panel (Biolegend).

In vitro *human monocyte-derived macrophage differentiation, polarization, tumor growth inhibition assay and flow cytometry*

Syngeneic monocytes, CD4 and CD8 T cells were isolated from leukopaks (Stemcell Technologies) from three healthy donors using negative selection EasySep kits (Stemcell technologies). T cells were frozen in liquid nitrogen while monocytes were differentiated into macrophages with 20ng/mL recombinant human M-CSF (Biolegend) in cRPMI (RPMI1640 (ThermoFisher Scientific) supplemented with 10% fetal bovine serum (ThermoFisher Scientific), penicillin/streptomycin (ThermoFisher Scientific) and 25mM HEPES (ThermoFisher Scientific)) for seven days. After washing, monocyte-derived macrophages (MDM) were polarized for an additional three days in cRPMI1640 containing recombinant human IL-4, IL-13 or TSLP (all from Biolegend). MDM were washed and 5x10^5^ MDM were co-cultured with 2.5x10^5^ syngeneic CD4 and CD8 T cells (5x10^5^ total T cells) in the presence of soluble CD3 (10μg/mL) and CD28 (2μg/mL) agonist antibodies in cRPMI for an additional three days. Total T cells were isolated by negative selection using the EasySep Human T Cell Isolation Kit (Stemcell Technologies) per the manufacturer’s instructions. Isolated T cells were co-cultured with 5x10^4^ A375 human tumor cells expressing nuclear GFP at a ratio of 8 T cells:1 A375 cell. Co-cultured T cells were restimulated with Immunocult human CD3/CD28 T cell activator (Stemcell Technologies), where appropriate. Longitudinal tumor growth curves were recorded and analyzed as described in the “In vitro *T cell priming and tumor growth inhibition assay and cytokine secretion measurement*” section.

After polarization, MDM were harvested and stained as described in (70). Briefly, MDM were stained with LIVE/Dead Fixable Aqua viability dye (ThermoFisher Scientific) before being fixed with diluted FoxP3/Transcription Factor Fixation/Permeabilization Concentrate (ThermoFisher Scientific) and frozen at -80°C in 10% dimethylsulfoxide/90% 1x Permeabilization Buffer (ThermoFisher Scientific). After thawing and washing in 1x Permeabilization Buffer, Fc receptors were blocked with Human TruStain FcX (Biolegend) and stained with the antibodies listed in Supplemental Table 1 in 1x Permeabilization Buffer containing Brilliant Stain Buffer Plus (BD Horizon). Stained MDM were acquired on a FACSymphony A5 analyzer (BD Biosciences) and analyzed using OMIQ software (Dotmatics).

*Generations of single cell suspensions from murine tissues*

CT26 tumors were processed using the Miltenyi Tumor Dissociation Kit and GentleMACS C tubes with slight modifications to the manufacturer’s instructions. Lyophilized enzymes were reconstituted per the manufacturer’s instructions and each tumor was minced by razor blade before digestion in 2.5mL of RPMI1640 (ThermoFisher Scientific) containing 4% enzyme D, 0.2% enzyme R and 0.5% enzyme A using the m_tdk1 program. The resulting single cell suspensions were filtered through 70μm mesh and washed with phosphate-buffered saline (PBS), counted and split between *ex vivo* restimulation and flow cytometry.

Tumor-draining lymph nodes (TDLN) and spleens were directly mashed through pre-moistened 70μ mesh (FisherBrand) using the plungers of 5mL syringes. The resulting single-cell suspensions either underwent erythrocyte lysis (spleens only) using ACK lysis buffer (ThermoFisher Scientific) or were directly washed and counted for flow cytometry.

*PacMAP dimensionality reduction and PARC clustering of flow cytometry data*

All PacMAP dimensionality reduction was performed using the following settings: nearest neighbors=10; number of output dimensions=2; distance metric=Euclidian; mid-near pairs to nearest neighbor ratio=0.5; further pairs to nearest neighbor ratio=2; random seed=2572; learning rate=1; number of iterations=450; KNN method=HNSW; embedding initialization method=random(1,2). All PARC clustering was performed using the following settings: KNN=30; KNN distance metric=Euclidian; local pruning=prune; standard deviations from mean for pruning=2; edge weighting for Leiden=weighted; N iterations for Leiden=5; small population cutoff=0.02(3).

**Supplemental Table 1.** Flow cytometry antibodies used for human samples.

| **Fluorophore** | **Antigen** | **Vendor** | **Catalog** | **Clone** |
| --- | --- | --- | --- | --- |
| BUV661 | HLA-DR | BD | 565073 | G46-6 |
| BV421 | CD38 | BD | 562444 | HIT2 |
| V450 | CD20 | BD | 561163 | L27 |
| BV510 | CD4 | BD | 562970 | SK3 |
| BV650 | CD8 | BD | 563821 | RPA-T8 |
| AF488 | CD3 | BD | 557694 | UCHT1 |
| PerCP-Cy5.5 | CD45RA | BD | 563429 | HI100 |
| PE-CF594 | CD197/CCR7 | BD | 562381 | 150503 |
| PE-Cy7 | Perforin | Biolegend | 308126 | dG9 |
| AF647 | Granzyme B | Biolegend | 372220 | QA16A02 |
| AF700 | CD45 | BD | 560566 | HI30 |
| BV786 | CD86 | BD | 740990 | 2331 (FUN-1) |
| APC-Cy7 | CD40 | Biolegend | 334324 | 5C3 |

**Supplemental Table 2.** Flow cytometry antibodies used for murine samples.

| **Fluorophore** | **Antigen** | **Vendor** | **Catalog** | **Clone** |
| --- | --- | --- | --- | --- |
| BUV395 | PD-L1 | BD | 745616 | MIH5 |
| BUV496 | CD38 | BD | 741090 | 90/CD38 |
| BUV563 | CD11b | BD | 741242 | M1/70 |
| BUV615 | CD86 | BD | 751557 | GL1 |
| BUV661 | CD11c | BD | 750482 | HL3 |
| BUV737 | F4/80 | BD | 749283 | T45-2342 |
| BUV805 | CD45 | BD | 748370 | 30-F11 |
| Super Bright 436 | SIRPα | ThermoFisher | 62-1721-82 | P84 |
| BV480 | CD4 | BD | 746566 | RM4-4 |
| BV510 | MHC-II | Biolegend | 107636 | M5/114.15.2 |
| BV570 | Ly6C | Biolegend | 128030 | HK1.4 |
| BV650 | Tim-3 | BD | 747623 | 5D12/TIM-3 |
| BV711 | CD40 | BD | 740700 | 3/23 |
| BV750 | CD44 | BD | 747255 | IM7 |
| BV785 | CD103 | Biolegend | 121439 | 2E7 |
| BB515 | OX40L | BD | 565214 | RM134L |
| AF532 | TCRβ | ThermoFisher | 58-5961-82 | H57-597 |
| BB700 | Ly6G | BD | 566435 | 1A8 |
| PE | Ly108 | Biolegend | 134606 | 330-AJ |
| PE-Cy7 | CD163 | Invitrogen | 25-1631-82 | TNKUPJ |
| AF647 | CD64 | Biolegend | 139322 | X54-5/7.1 |
| AF700 | CD8a | Biolegend | 100730 | 53-6.7 |
| APC-Cy7 | PD-1 | Biolegend | 135224 | 29F.1A12 |
| BV421 | T-bet | Biolegend | 644832 | 4B10 |
| PE-CF594 | Gata3 | BD | 563510 | L50-823 |
| APC | Perforin | Biolegend | 154304 | S16009A |
|  | TLR7 |  |  |  |
|  | IRF4 |  |  |  |
|  | IRF8 |  |  |  |
|  | Zbtb46 |  |  |  |
| BUV737 | CD24 |  |  |  |
| BV421 | Axl |  |  |  |
| APC-Fire750 | CD301 |  |  |  |

Ex vivo *restimulation of murine dissociated tissues and secreted protein quantification*

5x106 TME cells or 1x106 splenocytes were aliquoted for *ex vivo* restimulation in 96-well round-bottomed plates. Cells were resuspended in RPMI1640 (ThermoFisher) supplemented with 10% fetal bovine serum (ThermoFisher), penicillin and streptomycin (ThermoFisher), GlutaMAX-I (ThermoFisher), 55μM 2-mercaptoethanol (ThermoFisher Scientific) and 1x Cell Stimulation Cocktail (ThermoFisher Scientific). Conditioned media was harvested after 20 hours of stimulation and frozen at -80°C. Secreted proteins were quantified in thawed conditioned media using the mouse cytokine panel 2, mouse Th cytokine panel (12-plex), mouse hematopoietic stem cell panel (13-plex) and mouse proinflammatory chemokine panel (13-plex) fluid-phase Legendplex immunoassays (all from Biolegend) per the manufacturer’s instructions.

*Human CD8 T cell stimulation*

PBMC were isolated by density gradient centrifugation over Ficoll-Paque PLUS (Cytiva) from whole blood from a healthy human donor per the manufacturer’s instructions. Isolated PBMC were frozen in 10% DMSO/90% FBS in liquid nitrogen. Total T cells were isolated from thawed PBMC by negative selection using the EasySep Human T cell isolation kit (Stemcell Technologies) per the manufacturer’s instructions. 5x10^4^ total T cells were treated with 20ng/mL recombinant human IL-2 (Biolegend) alone, or combinations of agonist CD3 (10μg/mL, Biolegend) and CD28 (5μg/mL, Biolegend) antibodies with 20ng/mL recombinant human IL-2, 50ng/mL recombinant human IL-4 or TSLP (both Biolegend) and 10μg/mL isotype or antagonist IL-4 antibodies (both Biolegend) for two days at 37°C and 5% CO_2_. T cells were then washed and replated in the same conditions without CD3 and CD28 agonist antibodies and incubated for an additional four days.

**Supplemental Table 3.** Gene lists for IL-4/IL-13, TSLP and IFNγ transcriptional response signatures, and the T_H_2 deconvolution signature.

| **Transcriptional Response Signature Gene Lists** | | | |
| --- | --- | --- | --- |
| **IL-4/IL-13** | **TSLP** | **IFNγ** | **T_H_2** |
| *IL4R* | *CRLF2* | *IDO1* | *STAT6* |
| *IL2RG* | *IL7R* | *CXCL10* | *IL7R* |
| *IL13RA1* | *TSLP* | *CXCL9* | *PTGDR2* |
| *IL13RA2* | *BATF3* | *HLADRA* | *LGALS1* |
| *IL4* | *MYC* | *STAT1* | *IL4R* |
| *IL13* | *SLC9B2* | *IFNG* | *IL32* |
| *ATP6V1B2* | *CCT2* |  |  |
| *SERPINB4* | *HSP90AB1* |  |  |
| *CCL18* | *NOLC1* |  |  |
| *CD83* | *COTL1* |  |  |
| *ITK* | *PPM1K* |  |  |
| *POSTN* | *ZEB1* |  |  |
| *CCR2* | *CBLB* |  |  |
| *NFIL3* | *CCL17* |  |  |
| *SOCS1* | *CRLF2* |  |  |
| *NRN1* |  |  |  |
| *CCL11* |  |  |  |
| *BIRC5* |  |  |  |
| *CCL26* |  |  |  |
| *MUC5B* |  |  |  |

| **Day** | **TRT** | **REF** | **TC** | **logTC.se** | **TC.lower** | **TC.upper** | **p.value** | **DF** | **tail** |
| --- | --- | --- | --- | --- | --- | --- | --- | --- | --- |
| Day13 | ANTI-IL-4/IL-13 | ISOTYPE | 0.853 | 0.074 | NA | 1.134 | 0.177 | 53 | 1 |
| Day13 | ANTI-PD1 | ISOTYPE | 0.694 | 0.074 | NA | 0.922 | 0.018 | 53 | 1 |
| Day13 | ANTI-IL-4/IL-13/PD1 | ISOTYPE | 0.690 | 0.074 | NA | 0.918 | 0.017 | 53 | 1 |
| Day13 | ANTI-IL-4/IL-13/TSLP | ISOTYPE | 0.459 | 0.074 | NA | 0.610 | 0.000 | 53 | 1 |
| Day13 | ANTI-IL-4/IL-13/TSLP/PD1 | ISOTYPE | 0.396 | 0.074 | NA | 0.527 | 0.000 | 53 | 1 |
| Day13 | ANTI-IL-4/IL-13 | ANTI-IL-4/IL-13/PD1 | 1.236 | 0.074 | 0.878 | 1.738 | 0.219 | 53 | 2 |
| Day13 | ANTI-PD1 | ANTI-IL-4/IL-13/PD1 | 1.005 | 0.074 | 0.714 | 1.413 | 0.978 | 53 | 2 |
| Day13 | ANTI-IL-4/IL-13 | ANTI-IL-4/IL-13/TSLP | 1.857 | 0.074 | 1.321 | 2.611 | 0.001 | 53 | 2 |
| Day13 | ANTI-IL-4/IL-13 | ANTI-IL-4/IL-13/TSLP/PD1 | 2.152 | 0.074 | 1.530 | 3.027 | 0.000 | 53 | 2 |
| Day13 | ANTI-PD1 | ANTI-IL-4/IL-13/TSLP/PD1 | 1.750 | 0.074 | 1.245 | 2.461 | 0.002 | 53 | 2 |
| Day13 | ANTI-IL-4/IL-13/PD1 | ANTI-IL-4/IL-13/TSLP/PD1 | 1.742 | 0.074 | 1.239 | 2.449 | 0.002 | 53 | 2 |
| Day13 | ANTI-IL-4/IL-13/TSLP | ANTI-IL-4/IL-13/TSLP/PD1 | 1.159 | 0.074 | 0.824 | 1.630 | 0.391 | 53 | 2 |
| Day15 | ANTI-IL-4/IL-13 | ISOTYPE | 0.982 | 0.092 | NA | 1.401 | 0.466 | 53 | 1 |
| Day15 | ANTI-PD1 | ISOTYPE | 0.850 | 0.092 | NA | 1.212 | 0.223 | 53 | 1 |
| Day15 | ANTI-IL-4/IL-13/PD1 | ISOTYPE | 0.776 | 0.092 | NA | 1.107 | 0.118 | 53 | 1 |
| Day15 | ANTI-IL-4/IL-13/TSLP | ISOTYPE | 0.562 | 0.092 | NA | 0.802 | 0.004 | 53 | 1 |
| Day15 | ANTI-IL-4/IL-13/TSLP/PD1 | ISOTYPE | 0.492 | 0.092 | NA | 0.702 | 0.001 | 53 | 1 |
| Day15 | ANTI-IL-4/IL-13 | ANTI-IL-4/IL-13/PD1 | 1.266 | 0.092 | 0.827 | 1.939 | 0.271 | 53 | 2 |
| Day15 | ANTI-PD1 | ANTI-IL-4/IL-13/PD1 | 1.096 | 0.092 | 0.716 | 1.677 | 0.669 | 53 | 2 |
| Day15 | ANTI-IL-4/IL-13 | ANTI-IL-4/IL-13/TSLP | 1.748 | 0.092 | 1.142 | 2.675 | 0.011 | 53 | 2 |
| Day15 | ANTI-IL-4/IL-13 | ANTI-IL-4/IL-13/TSLP/PD1 | 1.997 | 0.092 | 1.305 | 3.057 | 0.002 | 53 | 2 |
| Day15 | ANTI-PD1 | ANTI-IL-4/IL-13/TSLP/PD1 | 1.727 | 0.092 | 1.129 | 2.644 | 0.013 | 53 | 2 |
| Day15 | ANTI-IL-4/IL-13/PD1 | ANTI-IL-4/IL-13/TSLP/PD1 | 1.577 | 0.092 | 1.030 | 2.413 | 0.036 | 53 | 2 |
| Day15 | ANTI-IL-4/IL-13/TSLP | ANTI-IL-4/IL-13/TSLP/PD1 | 1.142 | 0.092 | 0.746 | 1.749 | 0.533 | 53 | 2 |
| Day17 | ANTI-IL-4/IL-13 | ISOTYPE | 0.996 | 0.105 | NA | 1.493 | 0.493 | 53 | 1 |
| Day17 | ANTI-PD1 | ISOTYPE | 0.623 | 0.105 | NA | 0.934 | 0.028 | 53 | 1 |
| Day17 | ANTI-IL-4/IL-13/PD1 | ISOTYPE | 0.612 | 0.105 | NA | 0.918 | 0.024 | 53 | 1 |
| Day17 | ANTI-IL-4/IL-13/TSLP | ISOTYPE | 0.473 | 0.105 | NA | 0.710 | 0.002 | 53 | 1 |
| Day17 | ANTI-IL-4/IL-13/TSLP/PD1 | ISOTYPE | 0.352 | 0.105 | NA | 0.528 | 0.000 | 53 | 1 |
| Day17 | ANTI-IL-4/IL-13 | ANTI-IL-4/IL-13/PD1 | 1.628 | 0.105 | 1.001 | 2.645 | 0.049 | 53 | 2 |
| Day17 | ANTI-PD1 | ANTI-IL-4/IL-13/PD1 | 1.018 | 0.105 | 0.627 | 1.654 | 0.941 | 53 | 2 |
| Day17 | ANTI-IL-4/IL-13 | ANTI-IL-4/IL-13/TSLP | 2.104 | 0.105 | 1.296 | 3.417 | 0.003 | 53 | 2 |
| Day17 | ANTI-IL-4/IL-13 | ANTI-IL-4/IL-13/TSLP/PD1 | 2.826 | 0.105 | 1.739 | 4.591 | 0.000 | 53 | 2 |
| Day17 | ANTI-PD1 | ANTI-IL-4/IL-13/TSLP/PD1 | 1.768 | 0.105 | 1.089 | 2.872 | 0.022 | 53 | 2 |
| Day17 | ANTI-IL-4/IL-13/PD1 | ANTI-IL-4/IL-13/TSLP/PD1 | 1.736 | 0.105 | 1.069 | 2.820 | 0.026 | 53 | 2 |
| Day17 | ANTI-IL-4/IL-13/TSLP | ANTI-IL-4/IL-13/TSLP/PD1 | 1.343 | 0.105 | 0.826 | 2.183 | 0.228 | 53 | 2 |
| Day20 | ANTI-IL-4/IL-13 | ISOTYPE | 0.845 | 0.134 | NA | 1.416 | 0.294 | 53 | 1 |
| Day20 | ANTI-PD1 | ISOTYPE | 0.605 | 0.134 | NA | 1.013 | 0.054 | 53 | 1 |
| Day20 | ANTI-IL-4/IL-13/PD1 | ISOTYPE | 0.558 | 0.134 | NA | 0.936 | 0.032 | 53 | 1 |
| Day20 | ANTI-IL-4/IL-13/TSLP | ISOTYPE | 0.516 | 0.134 | NA | 0.865 | 0.018 | 53 | 1 |
| Day20 | ANTI-IL-4/IL-13/TSLP/PD1 | ISOTYPE | 0.363 | 0.134 | NA | 0.608 | 0.001 | 53 | 1 |
| Day20 | ANTI-IL-4/IL-13 | ANTI-IL-4/IL-13/PD1 | 1.514 | 0.134 | 0.815 | 2.813 | 0.185 | 53 | 2 |
| Day20 | ANTI-PD1 | ANTI-IL-4/IL-13/PD1 | 1.083 | 0.134 | 0.583 | 2.012 | 0.796 | 53 | 2 |
| Day20 | ANTI-IL-4/IL-13 | ANTI-IL-4/IL-13/TSLP | 1.638 | 0.134 | 0.883 | 3.040 | 0.115 | 53 | 2 |
| Day20 | ANTI-IL-4/IL-13 | ANTI-IL-4/IL-13/TSLP/PD1 | 2.329 | 0.134 | 1.254 | 4.324 | 0.008 | 53 | 2 |
| Day20 | ANTI-PD1 | ANTI-IL-4/IL-13/TSLP/PD1 | 1.666 | 0.134 | 0.898 | 3.093 | 0.104 | 53 | 2 |
| Day20 | ANTI-IL-4/IL-13/PD1 | ANTI-IL-4/IL-13/TSLP/PD1 | 1.538 | 0.134 | 0.829 | 2.854 | 0.168 | 53 | 2 |
| Day20 | ANTI-IL-4/IL-13/TSLP | ANTI-IL-4/IL-13/TSLP/PD1 | 1.422 | 0.134 | 0.765 | 2.641 | 0.260 | 53 | 2 |
| Day22 | ANTI-IL-4/IL-13 | ISOTYPE | 0.815 | 0.156 | NA | 1.489 | 0.286 | 53 | 1 |
| Day22 | ANTI-PD1 | ISOTYPE | 0.645 | 0.156 | NA | 1.179 | 0.115 | 53 | 1 |
| Day22 | ANTI-IL-4/IL-13/PD1 | ISOTYPE | 0.552 | 0.156 | NA | 1.009 | 0.052 | 53 | 1 |
| Day22 | ANTI-IL-4/IL-13/TSLP | ISOTYPE | 0.533 | 0.156 | NA | 0.974 | 0.043 | 53 | 1 |
| Day22 | ANTI-IL-4/IL-13/TSLP/PD1 | ISOTYPE | 0.337 | 0.156 | NA | 0.616 | 0.002 | 53 | 1 |
| Day22 | ANTI-IL-4/IL-13 | ANTI-IL-4/IL-13/PD1 | 1.477 | 0.157 | 0.717 | 3.045 | 0.284 | 53 | 2 |
| Day22 | ANTI-PD1 | ANTI-IL-4/IL-13/PD1 | 1.169 | 0.157 | 0.568 | 2.409 | 0.666 | 53 | 2 |
| Day22 | ANTI-IL-4/IL-13 | ANTI-IL-4/IL-13/TSLP | 1.529 | 0.156 | 0.743 | 3.148 | 0.243 | 53 | 2 |
| Day22 | ANTI-IL-4/IL-13 | ANTI-IL-4/IL-13/TSLP/PD1 | 2.417 | 0.156 | 1.173 | 4.979 | 0.018 | 53 | 2 |
| Day22 | ANTI-PD1 | ANTI-IL-4/IL-13/TSLP/PD1 | 1.913 | 0.156 | 0.929 | 3.940 | 0.077 | 53 | 2 |
| Day22 | ANTI-IL-4/IL-13/PD1 | ANTI-IL-4/IL-13/TSLP/PD1 | 1.636 | 0.156 | 0.795 | 3.368 | 0.177 | 53 | 2 |
| Day22 | ANTI-IL-4/IL-13/TSLP | ANTI-IL-4/IL-13/TSLP/PD1 | 1.580 | 0.157 | 0.767 | 3.258 | 0.210 | 53 | 2 |

**Supplemental Table 4.** Longitudinal tumor volume ANCOVA results demonstrates sustained growth inhibition by combined blockade of IL-4, IL-13, TSLP and PD1.

|  |  | ***IL4* RNA** | | ***IL13* RNA** | | ***TSLP* RNA** | | **IL4/IL13 transcriptional response signature** | | **TSLP transcriptional response signature** | | **T_H_2 deconvolution transcriptional signature** | |
| --- | --- | --- | --- | --- | --- | --- | --- | --- | --- | --- | --- | --- | --- |
| **Indication** | **TCGA abbreviation** | **HR** | **p-value** | **HR** | **p-value** | **HR** | **p-value** | **HR** | **p-value** | **HR** | **p-value** | **HR** | **p-value** |
| Testicular germ cell cancer | TGCT | 1.16 | 0.891 | 1.41 | 0.361 | 1.13 | 0.724 | 36 | 0.03 | 11.9 | 0.112 | 10.1 | 0.13 |
| Papillary kidney carcinoma | KIRP | 1.14 | 0.74 | 0.66 | 0.343 | 0.84 | 0.016 | 3.39 | 0.009 | 10.5 | <0.001 | 1.1 | 0.818 |
| Uveal melanoma | UVM | 0.64 | 0.628 | 1.07 | 0.939 | 1.04 | 0.774 | 4.3 | 0.146 | 8.94 | <0.001 | 3.28 | 0.016 |
| Prostate adenocarcinoma | PRAD | 0.02 | 0.235 | 1.05 | 0.869 | 0.93 | 0.766 | 0.4 | 0.421 | 8.96 | 0.107 | 0.36 | 0.176 |
| Lower Grade Glioma | LGG | 2.61 | 0.017 | 1.04 | 0.768 | 1.07 | 0.378 | 8.4 | <0.001 | 0.73 | 0.426 | 2.42 | <0.001 |
| Thymoma | THYM | 0.55 | 0.135 | 1.94 | 0.014 | 0.37 | 0.01 | 7.87 | 0.019 | 0.31 | 0.257 | 0.46 | 0.478 |
| Papillary thyroid carcinoma | THCA | 0.86 | 0.855 | 2.04 | 0.109 | 1.14 | 0.434 | 0.98 | 0.976 | 4.58 | 0.034 | 0.54 | 0.227 |
| Chromophobe renal cell carcinoma | KICH | <0.001 | 0.406 | 0.86 | 0.842 | 1.09 | 0.75 | 4.49 | 0.151 | 0.97 | 0.971 | 1.15 | 0.83 |
| Clear cell kidney carcinoma | KIRC | 1.86 | 0.025 | 1.39 | 0.002 | 1.19 | 0.002 | 2.81 | <0.001 | 1.76 | 0.025 | 2.02 | 0.004 |
| Mesothelioma | MESO | 1.26 | 0.202 | 0.96 | 0.764 | 0.95 | 0.552 | 1.92 | 0.047 | 1.62 | 0.131 | 2.11 | 0.014 |
| Stomach adenocarcinoma | STAD | 1.11 | 0.451 | 0.98 | 0.877 | 1.08 | 0.135 | 1.59 | 0.09 | 1.95 | 0.006 | 1.29 | 0.302 |
| Glioblastoma multiforme | GBM | 1.03 | 0.85 | 0.92 | 0.378 | 1.1 | 0.074 | 2.52 | 0.001 | 0.89 | 0.687 | 1.32 | 0.102 |
| Bladder Urothelial Carcinoma | BLCA | 1.00 | 0.986 | 0.991 | 0.926 | 1.09 | 0.05 | 0.99 | 0.94 | 1.33 | 0.123 | 0.94 | 0.659 |
| Lung squamous cell carcinoma | LUSC | 0.83 | 0.482 | 1.09 | 0.315 | 0.95 | 0.18 | 1.16 | 0.474 | 1.15 | 0.567 | 1.39 | 0.037 |
| Adrenocortical carcinoma | ACC | 1.76 | 0.066 | 1.37 | 0.046 | 0.94 | 0.528 | 1.61 | 0.504 | 0.571 | 0.322 | 0.99 | 0.976 |
| Liver hepatocellular carcinoma | LIHC | 1.4 | 0.202 | 0.96 | 0.605 | 1 | 0.943 | 0.9 | 0.647 | 1.27 | 0.299 | 0.58 | 0.01 |
| Ovarian serous cystadenocarcinoma | OV | 0.75 | 0.308 | 0.89 | 0.458 | 0.97 | 0.53 | 1.04 | 0.848 | 0.97 | 0.914 | 1.01 | 0.938 |
| Lung adenocarcinoma | LUAD | 0.97 | 0.845 | 0.94 | 0.528 | 0.86 | 0.004 | 0.82 | 0.299 | 1.15 | 0.518 | 0.86 | 0.458 |
| Esophageal cancer | ESCA | 1.44 | 0.391 | 1.22 | 0.328 | 1.04 | 0.53 | 1.08 | 0.832 | 0.85 | 0.633 | 0.95 | 0.889 |
| Colon adenocarcinoma | COAD | 1.02 | 0.975 | 0.83 | 0.173 | 1.06 | 0.414 | 0.96 | 0.871 | 0.94 | 0.84 | 0.72 | 0.297 |
| Head and neck squamous cell carcinoma | HNSC | 1.17 | 0.602 | 0.95 | 0.495 | 0.97 | 0.467 | 0.74 | 0.136 | 1.07 | 0.744 | 1.01 | 0.966 |
| Uterine corpus endometrial carcinoma | UCEC | 1.01 | 0.977 | 0.91 | 0.731 | 0.98 | 0.765 | 0.65 | 0.181 | 1.13 | 0.704 | 0.44 | 0.002 |
| Sarcoma | SARC | 0.42 | 0.005 | 0.73 | 0.024 | 1 | 0.98 | 0.52 | 0.004 | 1.2 | 0.473 | 0.47 | <0.001 |
| Cervical cancer | CESC | 0.77 | 0.672 | 0.79 | 0.174 | 1.03 | 0.645 | 0.59 | 0.09 | 1.08 | 0.808 | 1.23 | 0.652 |
| Cholangiocarcinoma | CHOL | 0.33 | 0.209 | 0.818 | 0.539 | 0.98 | 0.884 | 0.64 | 0.462 | 1.03 | 0.949 | 0.37 | 0.119 |
| Uterine carcinosarcoma | UCS | 0.86 | 0.85 | 1.01 | 0.962 | 0.99 | 0.911 | 1.13 | 0.864 | 0.54 | 0.211 | 0.9 | 0.8 |
| Pancreatic ductal adenocarcinoma | PAAD | 0.49 | 0.039 | 0.8 | 0.142 | 1.04 | 0.707 | 0.76 | 0.379 | 0.81 | 0.468 | 0.61 | 0.2 |
| Rectum adenocarcinoma | READ | 0.5 | 0.677 | 1.19 | 0.571 | 1.2 | 0.341 | 0.99 | 0.991 | 0.48 | 0.33 | 0.95 | 0.955 |
| Breast cancer | BRCA | 0.56 | 0.142 | 0.83 | 0.132 | 0.83 | 0.001 | 0.671 | 0.057 | 0.76 | 0.182 | 0.74 | 0.126 |
| Diffuse large B-cell lymphoma | DLBC | 0.81 | 0.39 | 1.57 | 0.08 | 0.8 | 0.421 | 0.44 | 0.357 | 0.89 | 0.883 | 0.5 | 0.541 |
| Cutaneous melanoma | SKCM | 0.45 | 0.014 | 0.74 | 0.048 | 0.94 | 0.14 | 0.6 | 0.002 | 0.54 | 0.002 | 0.71 | 0.008 |
| Pheochromocytoma & Paraganglioma | PCPG | 1.24 | 0.865 | 1.21 | 0.789 | 0.61 | 0.051 | 0.68 | 0.748 | 0.32 | 0.349 | 0.17 | 0.049 |

**Supplemental Table 5.** Median split hazard ratios and p-values of *IL4*, *IL13* and *TSLP* RNA, and transcriptional response and deconvolution signatures in individual TCGA cohorts.
